# Odyssey: reconstructing evolution through emergent consensus in the global proteome

**DOI:** 10.1101/2025.10.15.682677

**Authors:** Ankit Singhal, Shyam Venkatasubramanian, Sean Moushegian, Steven Strutt, Michael Lin, Connor Lee

**Affiliations:** Anthrogen, PBC. San Francisco, CA

## Abstract

We present Odyssey, a family of multimodal protein language models for sequence and structure generation, protein editing and design. We scale Odyssey to more than 102 billion parameters, trained over 1.1 *×* 10^23^ FLOPs. The Odyssey architecture uses context modalities, categorized as structural cues, semantic descriptions, and orthologous group metadata, and comprises two main components: a finite scalar quantizer for tokenizing continuous atomic coordinates, and a transformer stack for multimodal representation learning. Odyssey is trained via discrete diffusion, and characterizes the generative process as a time-dependent unmasking procedure. The finite scalar quantizer and transformer stack leverage the consensus mechanism, a replacement for attention that uses an iterative propagation scheme informed by local agreements between residues. Across various benchmarks, Odyssey achieves landmark performance for protein generation and protein structure discretization. Our empirical findings are supported by theoretical analysis.

## 1 Introduction

It is difficult to overstate the profound ubiquity of proteins as the universal workhorses that govern the living world; they operate as catalysts [70, 18, 42, 63], sensors [20, 47, 50], transporters [67, 57, 66], scaffolds [2, 77], switches [76, 52], motors [3], regulators [9, 4, 49], host defense effectors [11, 17, 78], and exist as a broad class of living *molecular machines*. The global proteome, the collection of all proteins that exist today, has emerged over billions of years of biological evolution molded by mutation, drift, recombination, and the biophysical constraints that govern folding and function.

The resulting distribution is neither random nor uniformly smooth: it is a highly structured, sparsely populated manifold in an astronomically large feature space.

Advances in modern genomic sequencing, structural imaging techniques, and computational modeling have allowed us to subsample billions of sequences [1, 19, 46, 64] along with hundreds of millions of structures [33, 41] and functional annotations [32] from this manifold. Current efforts focus on using this influx of data to uncover the latent grammar underpinning evolution [45, 13, 62], and to leverage that understanding in the design of these molecular machines, subject to our own specifications.

Much of the work done toward this end has mirrored recent advancements in natural language processing (NLP), particularly in regard to the transformer architecture [74], which has seen widespread adoption across various domains. Over the last decade, NLP has largely adopted the view that natural language can be modeled as tokens arranged as a one-dimensional object, with meaning emerging from distributional regularities, captured by self-attention, and learned via next-token (autoregressive modeling) or masked-token (masked-language modeling) objectives [21, 74, 6, 27]. This framing reflects the character of human language: predominantly local syntax, punctuated by sparse long-range dependencies, with limited invariance beyond word order, and semantics that are compositionally expressed in sequences [24]. Proteins, by contrast, are intrinsically multimodal and geometric: a one-dimensional amino-acid sequence maps — epistatically — to a three-dimensional structure whose function depends on local, distance-dependent, and orientation-dependent interactions under thermodynamic constraints.

Despite this, frontier generative protein language models have largely imported NLP defaults with minimal adaptation [28, 10].

We posit that two of the primary architectural choices and training methodologies employed by NLP models — the self-attention mechanism and the autoregressive/masked language modeling objective —necessitate substantial redesign for improved application to data classes such as proteins. We do so with the express intent of preserving the scaling properties of transformer architectures — properties often lost in overly inductive models built for hyperspecific applications [31, 72].

We now frame our methodology with the following intuitions:

1. Unlike text, where sparse, long-range relations can be made *without* consequence to intervening tokens (e.g., “Rosalind” at the start of a paragraph and “she” at the end), proteins realize long-range dependencies only through the geometry that is constrained by a covalent backbone. When two residues, *i* and *j*, come into proximity in three-dimensions, their intervening span is necessarily co-perturbed. As such, protein dependencies are many-bodied and locally cooperative rather than arbitrarily pairwise across a sequence [22, 55]. In other words, text tolerates ‘teleportations’ in dependency with no cost to tokens in between; proteins cannot. Intuitively, amino acids in chains are more similar ‘pearls on a string’ rather than independently moving, free-floating pieces. Motivated by this simple inductive bias of local adherence, we introduce the *consensus* mechanism, which replaces global self-attention with repeated local agreement on a sparse graph. This bias is not protein-specific and is observed in various domains (e.g., other biomolecules).
2. During training, the learning rate quantifies the stride of the optimizer — too short and it shuffles; too long and it stumbles. Scaling to large model sizes renders learning rate tuning more difficult, and more brittle. We hypothesize that, benchmarked against attention, consensus offers greater robustness to learning rate misspecification. We further note consensus scales with 𝒪(*L*) complexity with respect to the length of the sequence, *L*, whereas attention scales with 𝒪(*L*^2^) complexity, opening the door to training larger, longer-context models that can generate more intricate proteins.
3. Evolution can be viewed, at its coarsest scale, as a stochastic proposal (mutation) coupled with a selection operator that prunes unfavorable variants over time. Discrete diffusion mirrors these dynamics: it corrupts sequences with injected noise and learns a reverse-time denoiser that concentrates probability mass on functionally consistent proteins under an objective [44]. We hypothesize that this interplay between technology and biology enhances *de novo* protein generation.

While such intuition far from guarantees superior performance (or, in fact, performance of any kind), it motivates our hypothesis of using consensus and discrete diffusion as the generative backbone for proteins. The organization of this paper is as follows. In Section 2, we delineate Odyssey, including consensus, discrete diffusion, architecture, and alignment. In Section 3, we present empirical results comparing consensus with attention and discrete diffusion with masked language modeling, alongside ablations and scaling analyses. In Section 4, we present empirical results for alignment, and finally, in Section 5, we summarize our paper. The main contributions of this paper are as follows:

1. We introduce *Odyssey*, a family of multimodal generative protein language models that extends to 102B parameters and supports sequence and structure generation, conditional design, and editing.
2. We develop a finite scalar quantizer (FSQ) for atomic structure coordinates that achieves benchmark protein discretization performance.
3. We propose the *consensus* mechanism — a drop-in replacement for attention that demonstrates properties favorable for the scaling of transformer architectures.
4. We demonstrate that training with discrete diffusion yields lower perplexities than masked language modeling in joint protein sequence and structure prediction.

## 2 Odyssey

The Odyssey architecture comprises two primary components — a *finite scalar quantizer* (FSQ) for tokenizing protein structure, and a *transformer stack* for multimodal representation learning. The FSQ and transformer stack leverage transformer blocks derived through the *consensus* mechanism, a novel, drop-in replacement for attention. We provide a short introduction to consensus in Section 2.1, with comprehensive details provided in Section D of the Appendix. The FSQ quantizes continuous atomic coordinates into discrete structure tokens, which are embedded and summed with the embedded sequence tokens (context is provided). The transformer stack operates on this joint representation, and is trained using discrete diffusion [44], which we discuss in Section 2.2 (see Section E of the Appendix for full derivation). We outline the Odyssey architecture, training, generation, and alignment schemes in Sections 2.3, 2.4, and 2.5, with full details provided in the Appendix. For the derivations that follow, we consider *L* to be the maximum sequence length.

### 2.1 Consensus

We replace the attention mechanism with a local, iterative propagation rule motivated by protein data, where nearby residues tend to agree. Rather than mixing all tokens in one hop, consensus mixes over a sparse graph and propagates information outward across layers. A single consensus layer performs one gradient step that resolves local *disagreements* between neighboring tokens while preserving sharp boundaries. The comprehensive outline of consensus is presented in Section D of the Appendix, and we present an abridged version of the self-consensus mechanism as follows.

We invoke a flattened batch with *B* = 1. Given token embeddings *y* = [*y*^(0)^, …, *y*^(*L−*1)^]^⊤^ ∈ ℝ^*L×d*^, we first project to a *consensus feature space* by an affine map, ∀*i* ∈ *{*0, …, *L* − 1*}*:

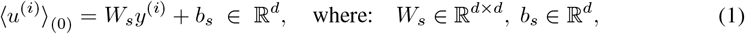

and denote *𝒰*_(0)_ = [⟨*u*^(0)^⟩_(0)_, …, ⟨*u*^(*L−*1)^⟩_(0)_]^*⊤*^. We connect each position *i* to neighbors *j* within a window *w*, forming a directed edge set *E* = *{*(*i, j*) : 0 ≤ *i, j* ≤ *L* − 1, 0 *<* |*i* − *j*| ≤ *w}*, such that 𝒩^(*i*)^ = *{j* : (*i, j*) ∈ *E}*. Each directed edge (*i, j*) forms a positive-definite *consensus weight matrix*:

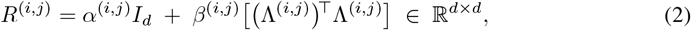

where *α*^(*i,j*)^, *β*^(*i,j*)^ ∈ ℝ_*>*0_, Λ^(*i,j*)^ ∈ ℝ^*r×d*^ are produced using small MLPs from the pairwise context, [*y*^(*i*)^; *y*^(*j*)^] ∈ ℝ^2*d*^. The *α*^(*i,j*)^*I*_*d*_ matrix induces *isotropic* coupling that mixes all features equally, and penalizes the pairwise difference uniformly in every direction. Contrastingly, the *β*^(*i,j*)^(Λ^(*i,j*)^)^*⊤*^Λ^(*i,j*)^ matrix is *anisotropic*, and confines propagation to at most *r* informative directions. When *r* = 0, the consensus weight matrix reduces to purely isotropic coupling. The index [*R*^(*i,j*)^]^(*a,b*)^ measures how the *b*-th feature difference influences the *a*-th feature update at node *i*.

We subsequently define the following quadratic energy over the graph:

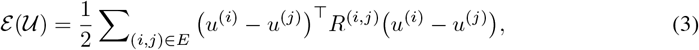

which depends only on pairwise differences, and is thus invariant to adding the same vector *c* ∈ ℝ^*d*^ to all *u*^(*i*)^ (consensus subspace). As such, ℰ is convex and strictly convex on the orthogonal complement of this subspace. The mechanism takes one explicit step via the *consensus gradient update*:

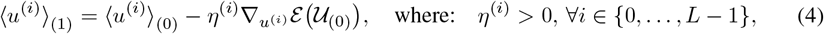

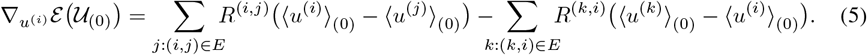

Intuitively, node *i* exchanges its current disagreement with each neighboring node, and moves by a weighted average of these differences. We use a learned step size to accelerate mixing in homogeneous regions (large *η*^(*i*)^) and preserve sharp boundaries (small *η*^(*i*)^). We map back via an output projection:

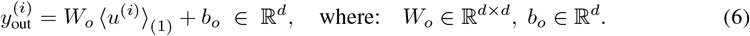

For edge hidden dimension *ξ* (see Section D.1 of the Appendix), the cost of self-consensus scales as:

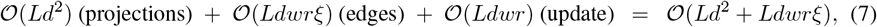

which is linear in sequence length, *L*. This contrasts with the quadratic token-pairing of self-attention. Intuitively, consensus is a content-adaptive propagation step on a graph Laplacian with matrix-valued edge couplings [38]: *α*^(*i,j*)^*I*_*d*_ provides a stable baseline, and *β*^(*i,j*)^(Λ^(*i,j*)^)^*⊤*^Λ^(*i,j*)^ selects at most *r* principal directions of agreement, learned from embedding pairs. We illustrate the self-consensus mechanism via a simplified example in Figure 2, where *d* = 2.

**Figure 1:**
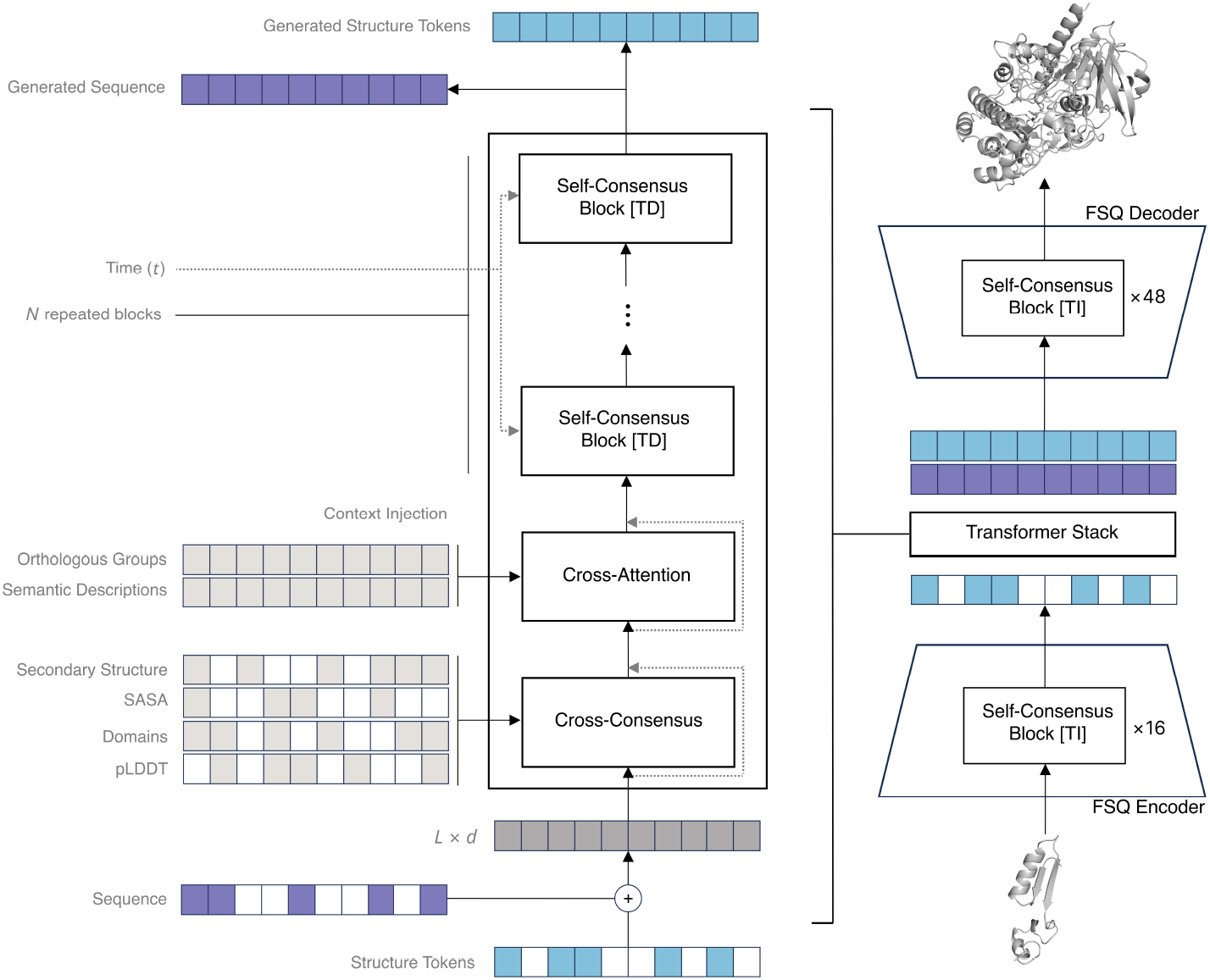
Illustration of the Odyssey architecture. Odyssey encompasses an FSQ for tokenizing continuous atomic coordinates, and a transformer stack for multimodal representation learning. The corrupted atomic coordinates are fed to the FSQ encoder and positionally summed with the corrupted sequence, before being passed to the transformer stack to generate a complete sequence and structure. The generated structure tokens are decoded into predicted atomic coordinates via the FSQ Decoder.

**Figure 2:**
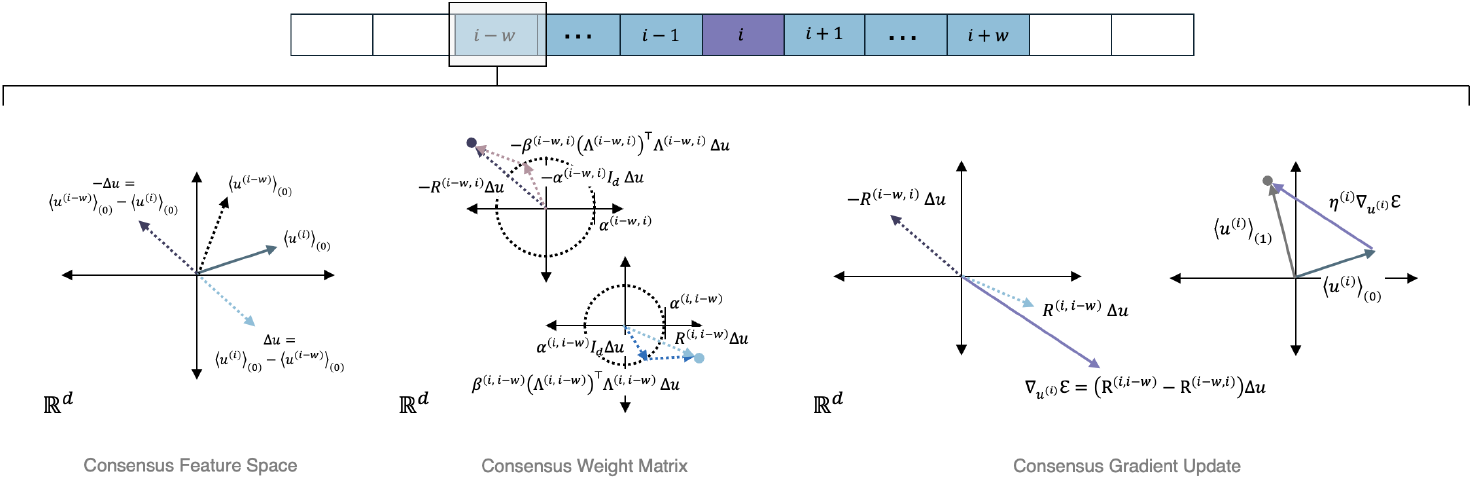
Illustration of self-consensus for *d* = 2, depicting the local neighborhood principle and the operative steps of self-consensus. We show an abridged consensus gradient update for position index *i* via index *i* − *w*. We repeat this process ∀*j* ∈ {*i* − *w*, …, *i* + *w*}.

We formally outline self-consensus (source tokens negotiate with local neighbors) in Section D.1 of the Appendix, cross-consensus (source tokens negotiate with nearby context tokens while holding context fixed) in Section D.2 of the Appendix, multi-head consensus (through *H* parallel subspaces) in Section D.3 of the Appendix, and rotary positional embeddings (RoPE) [71] for consensus in Section D.4 of the Appendix. The Odyssey architecture leverages these outlined mechanisms.

### 2.2 Discrete diffusion

As we delineate in Section E of the Appendix, the Odyssey transformer stack leverages score-entropy discrete diffusion [44] in its training and generation methodology. We begin by formally defining the diffusion process that applies masks, which we term the *forward process*.

Under the forward process, each token of the corruptible tracks is transformed via a continuous-time Markov chain (CTMC). At each time, *t* ∈ [0, *T*], a residue remains the same *or* transitions to `MASK` (for source tracks) or `IGN` (for context tracks). The `MASK`/`IGN` tokens are *absorbing* — once a position is corrupted to `MASK`/`IGN`, it remains corrupted for the remainder of the forward CTMC.

To simulate the forward process, we define an *instantaneous noise schedule, σ*(*t*), which is monotonically increasing in *t*. The forward process corrupts token 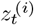 using the following procedure:

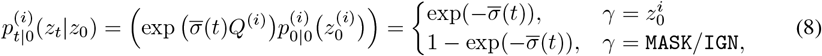

where 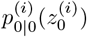 is the one-hot vector corresponding to 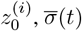 is the cumulative noise level at time *t*, and where *Q* is given in Eq. (64). We depict the forward process for a simplified protein sequence in Figure 3. The forward process of Odyssey is detailed further in Section E.1 of the Appendix.

**Figure 3:**
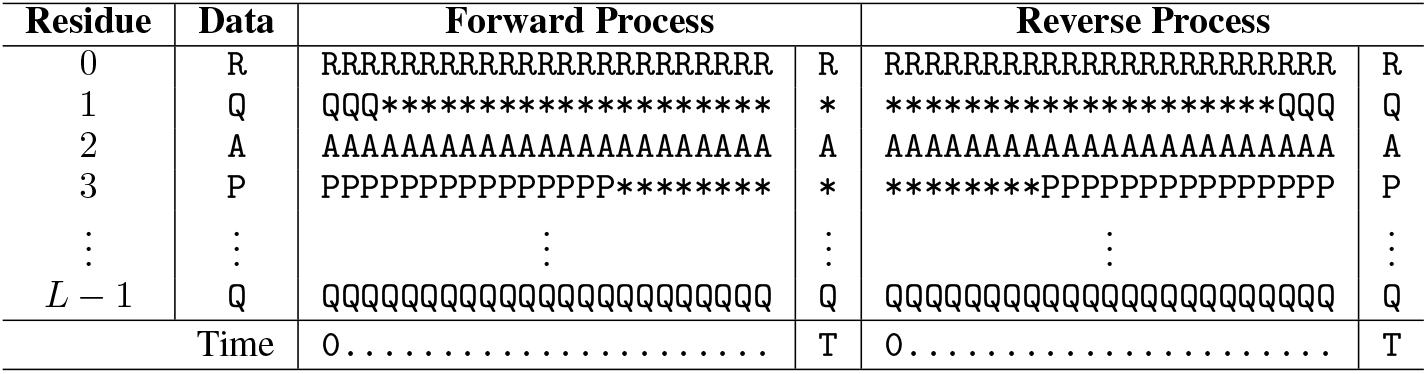
Discrete diffusion forward and reverse processes of the sequence track under the ideal transformer stack for a length-*L* protein, omitting all special tokens except for `MASK` (denoted by *). The sequence `RQAP · · · Q` is corrupted to `R*A* · · · Q` in the forward process, and the ideal transformer stack reconstructs `RQAP · · · Q`. The final timestep of each process is displayed in the adjacent column.

Odyssey aims to simulate the reverse of the forward process, diffusing the `MASK`s comprising the source tracks into content tokens. We denote this as the *reverse process*. Letting *p*_source,*t*_ denote the probability law over source track tokens at time *t* during the forward process, we have that:

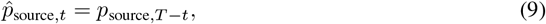

where 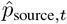 denotes the probability law of all source track tokens at time *t* of the reverse process. For position *i*, the *i*-th column of the output, 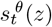, from Odyssey’s transformer stack is:

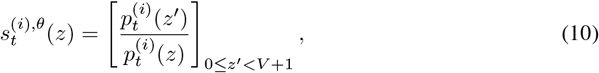

where *V* is the size of the vocabulary. This probability ratio characterizes the requisite quantity to “adjust” 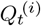 into a infinitesimal generator matrix 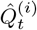 of the reverse-time CTMC:

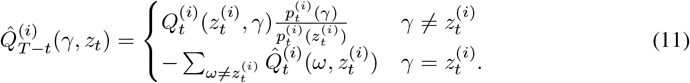

Using this infinitesimal generator matrix, the continuous-time reverse process can be simulated as the solution of a vector-valued ordinary differential equation. Odyssey utilizes Tweedie’s method to numerically integrate the reverse process. We further detail the Odyssey reverse process in Section E.2 of the Appendix, and illustrate the reverse process for a simplified protein sequence in Figure 3.

### 2.3 Architecture and training

As detailed previously, the Odyssey architecture comprises an FSQ for quantizing continuous atomic coordinates into discrete structure tokens, which can be embedded and summed with the embedded sequence tokens (with the embedded context tracks introduced through cross-consensus and cross-attention). The transformer stack operates on this joint representation. We provide a comprehensive description of the FSQ in Section F.2 of the Appendix, and of the transformer stack in Section F.3 of the Appendix. We schematize Odyssey in Figure 1, and provided an abridged summary as follows.

#### Finite scalar quantizer

To discretize continuous atomic coordinates into structure tokens, we use an FSQ [48] to avoid the codebook collapse and auxiliary components typical of VQ-VAEs [73]. The FSQ partitions coordinate space by an axis-aligned lattice with codebook ℋ; in Odyssey we instantiate ℋ = {0, …, 6} × {0, …, 4}^4^, yielding the structural vocabulary 𝒱_struct_ = {0, …, |*ℋ*| − 1}. The encoder maps per-residue coordinates to codes using a projector schematic (two convolutional blocks followed by 16 *time-independent* self-consensus blocks), then uses lattice quantization and lexicographic indexing to produce 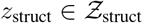. Decoding proceeds by inverse indexing and lattice dequantization, followed by an expander (16 time-independent self-consensus blocks (see Section F.1 of the Appendix) in stage-1 and 48 time-independent blocks in stage-2, preceding two convolutional blocks) to reconstruct coordinates. In FSQ stage-1 training, the model learns to reconstruct the atomic coordinates in the 3-backbone scheme. In stage-2 training, the trained encoder is frozen, and a larger decoder learns to reconstruct the atomic coordinates in the atom-14 scheme [33]. We further detail FSQ training in Section G.1 of the Appendix.

#### Transformer stack

The transformer operates on a joint source representation formed by summing the embedded sequence tokens 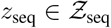, with the embedded structure tokens, 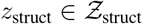 where 𝒱_seq_ = 0, …, *V*_seq_ 1 (see Section C of the Appendix). The auxiliary context is injected residually through two pathways: position-aligned tracks (secondary structure, solvent accessibility, domains, pLDDT) enter via cross-consensus, while global tracks (orthologous groups, semantic text) enter via cross-attention. The transformer stack uses *time-dependent* self-consensus blocks (see Section F.1 of the Appendix) (conditioned on the diffusion time step *t*) in place of self-attention blocks, enabling local, direction-aware propagation. Training follows the method of score-entropy discrete diffusion [44]: we sample a single time step, *t* ∼ 𝒰([0, *T*]), per protein, apply the forward process to overwrite a *t*-dependent subset of tokens using corruption symbols, and train the transformer stack to predict probability ratios at the masked sites. The objective is the diffusion-weighted denoising score entropy, applied separately to the sequence and structure tracks via an estimator and combined with a weighted sum. This objective upper-bounds the negative log-likelihood of the ground-truth token. We detail transformer training in full in Section G.2 of the Appendix.

### 2.4 Generation

Generation begins from an *original* protein and a designer-specified binary mask 𝕄 over residues and tracks. Positions marked to be kept retain their ground-truth tokens, while the remainder are replaced by corruption symbols (`MASK` for source tracks; `IGN` for context tracks), yielding designer-corrupted start state, *z*_*T*_. Equivalently, letting *z*_0_ denote the uncorrupted tokens, we have:

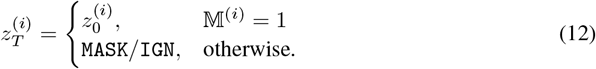

For positions with 𝕄^(*i*)^ = 0, replacing content tokens by `MASK/IGN` approximates (for a sufficiently large value of *T*) sampling from the stationary distribution of the forward-process 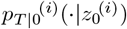. From this *z*_*T*_, we simulate the reverse CTMC in discrete steps with stride Δ*t* from *t* = *T* to *t* = 0.At each step, masked positions are updated by sampling from a learned, time-dependent reverse kernel, 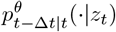, that is assembled from the per-position conditionals 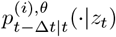 of Remark H. If any `MASK` tokens remain immediately before the final timestep (*t* = Δ*t*), we sample from the conditional 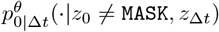. The mask is updated monotonically as tokens are un`MASK`ed so that previously preserved positions remain fixed. After obtaining *z*_0_, we decode structure tokens with the stage-2 FSQ to atom-14 coordinates, while sequence tokens are mapped back to amino acids. We outline the full generation scheme using the Odyssey transformer stack in Section H of the Appendix.

### 2.5 Alignment

We align only the sequence reverse kernel, treating all other tracks as conditioning, using D2-DPO [12]. Each batch contains a naturally-occurring wild type *z*_seq,0_ and *B* mutants with scalar scores, where each score denotes the measured fitness of the mutant over the wild type. We enumerate within-batch pairs, keep those with positive gaps, sort by gap magnitude, and select the top-*k* (winner, loser) pairs. Let *j* ∈ *{*0, …, *k* − 1*}*. For the *j*-th pair 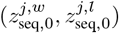, we sample a time step, *t*_*j*_ ∼ 𝒰([0, *T*]), and compute (for each protein) the preference difference, 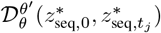 of the aligned model, *θ*^*′*^, over the reference model, *θ*. We form the preference margin:

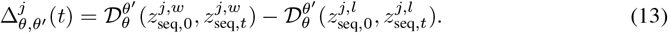

The empirical D2-DPO risk averages this objective over admissible pairs, and is given by:

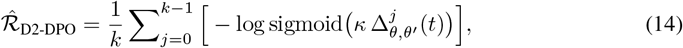

with temperature parameter *κ >* 0. Post-alignment, we report the validation score, 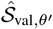, by treating each generated sequence, 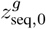, as a mutant against the wild type, *z*_seq,0_. We delineate the complete alignment methodology, with all expressions defined, in Section I of the Appendix.

## 3 Evaluations of unaligned capabilities

The following experiments invoke normalized per-token estimates of perplexity for discrete diffusion and masked language modeling. Discrete diffusion estimates the joint probability of the full protein; we compute the per-token perplexity by first normalizing score entropy with protein length before exponentiation. Masked language models compute the conditional probability of masked residues given unmasked residues; we compute per-token perplexity by first normalizing cross-entropy via the number of masked positions before exponentiation, with ℐ = *{i* : *i* ∈ *{*0, …, 2*L* − 1*}*, 𝕄^(*i*)^ = 0*}*:

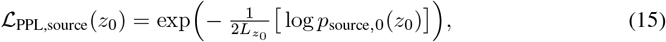

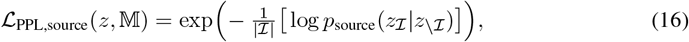

where 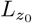 denotes the number of residues in the protein *z*_0_. Eq. (15) and Eq. (16) aim to estimate per-token perplexity. As the perplexity of Eq. (16) is conditioned upon unmasked positions, we do not claim a strict correspondence between these respective metrics.

We present empirical results demonstrating the data-efficient scaling properties exhibited by Odyssey in Section 3.1, the robustness of consensus to variable learning rates in Section 3.2, and the improved inference-time results yielded by discrete diffusion over masked language modeling in Section 3.3. For masked language modeling, we consider the fixed 15% schedule in [21] (*simple masking*), and the mixed betalinear-30/cosine schedule from [28] (*complex masking*).

### 3.1 Odyssey scales incredibly data-efficiently

The Odyssey family exhibits a pronounced ability to scale data-efficiently across orders of magnitude in model size. We validate this property over a sweep of smaller Odyssey variants to motivate scaling to production-ready 1.2B, 12B, and 102B parameter models. All models for the scaling experiments were trained with flat learning rates, proportional to their respective batch sizes (see Table 11). The dataset on which these scaling experiments were compiled is detailed in Section B of the Appendix.

Following the guidelines of [37], we fit a power law linking model size, 𝒦, tokens, ℬ, and validation loss, 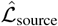, utilizing a sweep of smaller Odyssey variants (see Fig. 4). From this power law, the compute-optimal allocation of parameters for a given dataset size is given by:

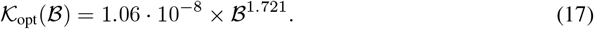

**Figure 4:**
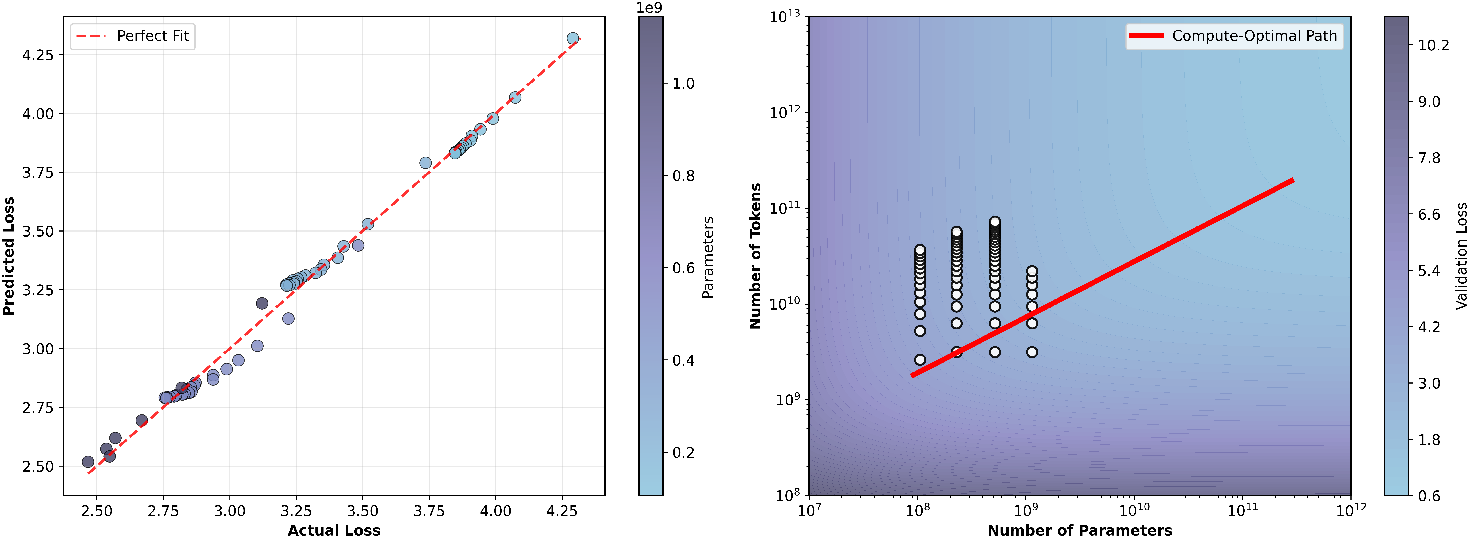
Scaling results for Odyssey variants across 142M — 1.2B parameters and up to 80B tokens. We depict the validation DWDSE loss (mean of sequence/structure DWDSE losses) from Section G.2. (a) Actual vs predicted losses using the modified Kaplan Fit (*r*^2^ = 0.993). (b) Compute-optimal frontier plotted with trained parameterizations and dataset size pairs, with a loss landscape.

Eq. (17) shows that for a fixed target loss, Odyssey needs far fewer tokens than would be required by proportionally smaller models trained on larger corpora [10, 28]. Guided by this analysis, we scale to Odyssey-1.2B, 12B, and 102B, where the 1.2B and 12B models lie on the frontier predicted by Eq. (17). Due to practical constraints relating to the compute availability used to train Odyssey, we scale our largest production model — trained across 1.1 × 10^23^ FLOPs — to 102B parameters as opposed to the optimal 142B parameters predicted by our scaling law (see Table 1). The training configurations for these models are detailed in Table 10.

**Table 1:**
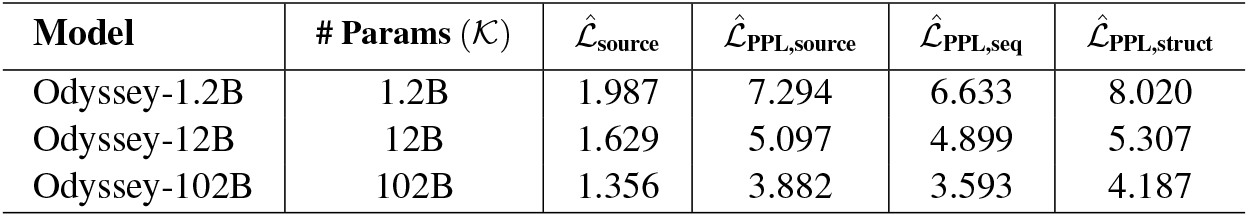
Validation perplexities, 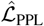 (of Eq. (15)), for Odyssey-1.2B, 12B, 102B.

### 3.2 Consensus exhibits properties favorable for the scaling of transformers

In large-scale transformer training, hyperparameter selection quickly becomes a bottleneck. Exhaustive sweeps that are tractable for small models are prohibitively expensive at billion-parameter scales [80], so practitioners fall back on heuristic extrapolations from smaller runs [25]. For models that are sensitive to these choices, even mild misspecification can blunt or erase the expected gains from scaling [56]. In such regimes, mechanisms that remain performant under mild misspecification of the learning rate, Γ, are disproportionately valuable.

We train several masked language models (see Figure 5) using simple masking via self-consensus and cross-consensus, and separately via self-attention and cross-attention. As the model size increases, consensus backbones sustain near-optimal perplexity across a *broader interval*, 𝒲, of Γ than attention backbones. When Γ exceeds its optimum, consensus degrades gradually, whereas attention exhibits an abrupt cliff, alongside large intermittent spikes in both training and validation loss. Qualitatively, 𝒲 remains relatively stable for consensus across scales, but contracts rapidly for attention. At 35M parameters, both mechanisms tolerate a range of learning rates. By 552M parameters, attention’s 𝒲 narrows to a small band with visible loss excursions, while consensus maintains its original span.

**Figure 5:**
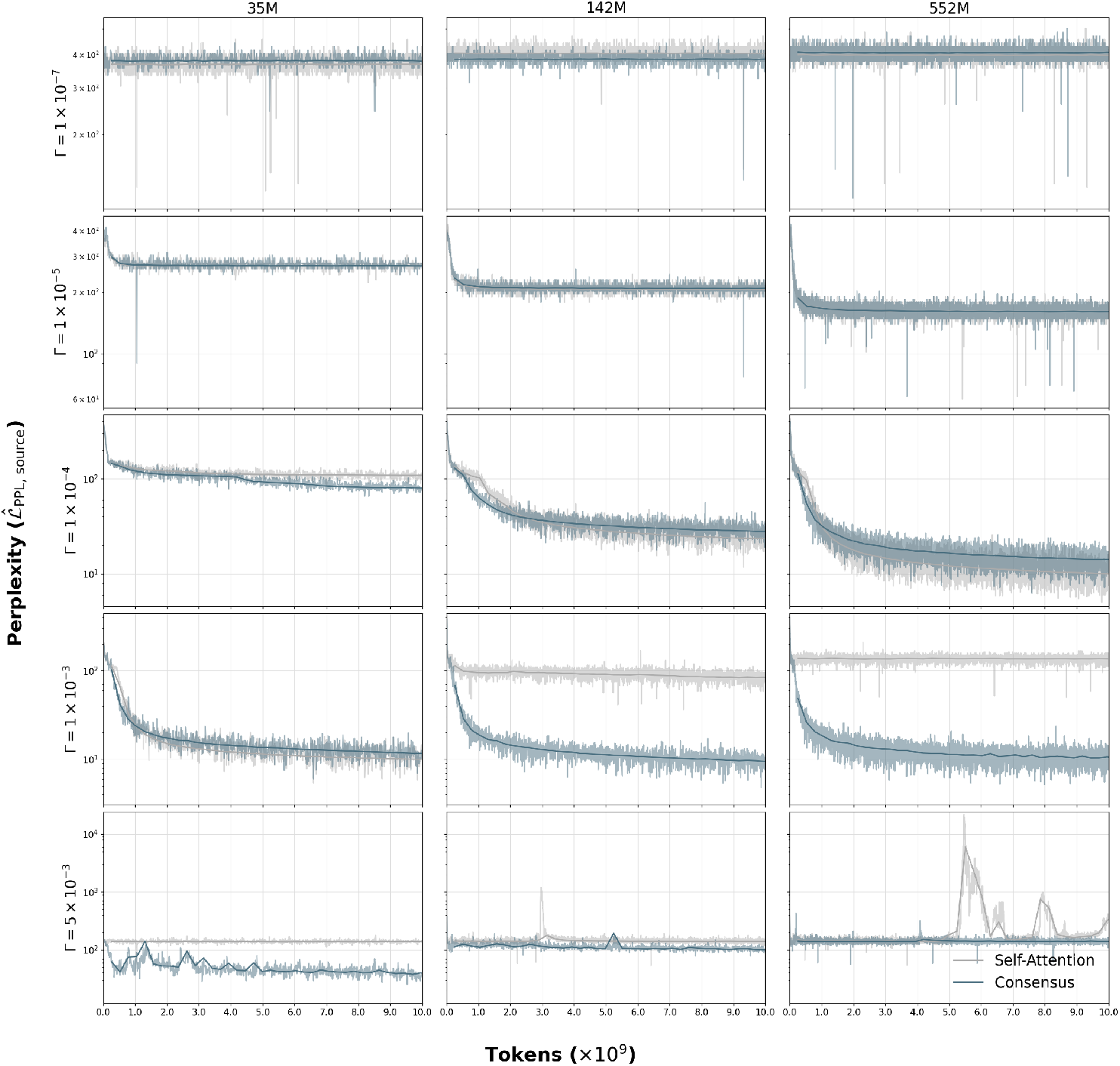
We train several masked language models (35M, 142M, 552M parameters) with consensus (blue) or attention (gray) using simple masking. The training perplexities form the background, and the validation perplexities are the solid lines in the foreground. For variable learning rate, Γ, we see improved robustness using consensus vs attention, with heightened effects at larger model sizes.

The wider 𝒲 under consensus has three major consequences particularly valuable for the scaling of transformers: (1) higher odds of near-optimal training at fixed total compute budget, (2) faster convergence when training with aggressive schedules, and (3) operational robustness.

Let *p*^⋆^ denote the probability that a coarse sweep (few Γ and a fixed seed) hits in the interval. Since *p*^⋆^ ∝ 𝒲 for a fixed proposal distribution over Γ, the larger 𝒲 for consensus implies a materially higher hit-rate than 𝒲 for attention, and the gap widens with scale. For a fixed budget allocated across ablation and scale, consensus is more likely to deliver models trained close to their optimal Γ.

Instability in attention becomes pronounced at high learning rates [35, 83, 79], which in turn constrains both warmup length and peak Γ; consensus, by contrast, tolerates higher peaks. This tolerance enables more aggressive learning rate schedules that reduce tokens-to-target perplexity—i.e., faster convergence for a fixed optimizer and batch size [81, 25].

Wider learning rate tolerance (larger 𝒲) makes training less sensitive to incidental drift that perturbs the ‘felt’ learning rate 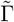 (e.g., microbatch changes, dataloader variance, minor optimizer tweaks). As 𝒲 is broader for consensus, small shifts in 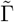 remain inside 𝒲, reducing failed runs at scale.

While consensus was motivated by the data modality Odyssey was trained for, we offer no intuitive reasoning as to whether this scaling favorability can be ascribed to the consensus mechanism being better suited for protein generation specifically, or whether it is inherent to the mechanism itself and can be replicated in transformer architectures over a wide variety of other tasks.

### 3.3 Discrete diffusion yields better inference-time results than masked language modeling

From an evolutionary perspective, protein sequences arise from mutations filtered by selection. In discrete diffusion, the forward process induces mutations, and the learned reverse process concentrates probability on sequences that satisfy global compatibility constraints, akin to selection. As the model must denoise from progressively degraded contexts, it learns coordinated, multi-residue corrections.

We benchmark discrete diffusion versus masked language modeling across three scales under matched architectures with consensus, and optimization (see Table 13). We compute perplexities using Eq. (15) and Eq. (16), and illustrate our results in Figure 6. Across all model sizes, we observe that discrete diffusion yields lower training perplexities than complex masking, and has lower or comparable training perplexities than simple masking. The training of discrete diffusion models, however, exhibits greater stochasticity than that of either masked language model.

**Figure 6:**
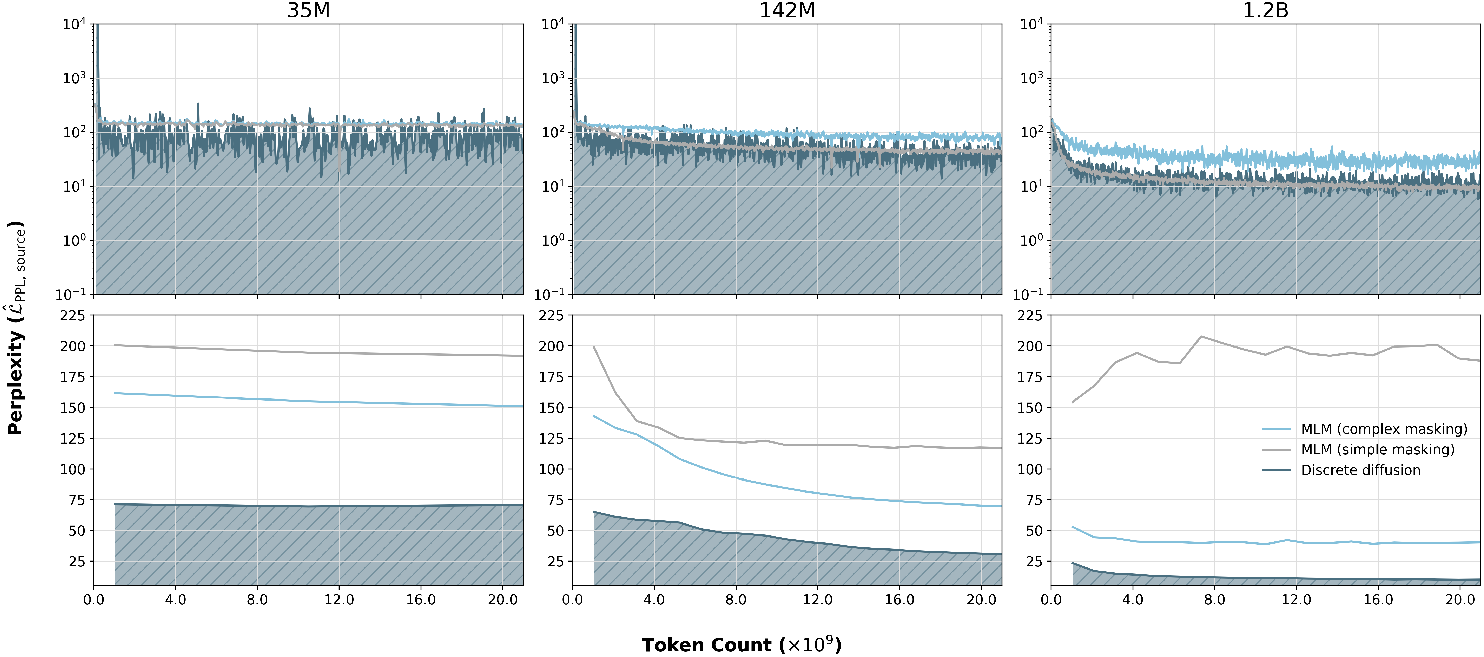
We train several models (35M, 142M, 1.2B parameters) with either masked language modeling (gray for simple masking, light blue for complex masking) or discrete diffusion (dark blue). We hatch the area under the discrete diffusion perplexities to signify the DWDSE loss as an upper bound (see Section G.2 of the Appendix). The masked language model perplexities were derived by taking the exponential of the sum of the sequence and structure cross-entropy losses. We plot the training perplexities in (a) and the validation perplexities in (b), as described in Section 3.3.

For validation, we benchmark all models on the discrete diffusion schedule, to evaluate robustness to the varying levels of corruption seen during generation. We observe that the discrete diffusion models are more performant than their masked language counterparts, with the simple-masking-trained 1.2B parameter model overfitting considerably to its own masking scheme.

We note that the validation loss, 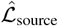, for discrete diffusion is an *upper bound* of the loss 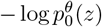 representing entire-protein generation, as shown by the hatched area in Figure 6. Consequently, the difference in validation loss between discrete diffusion and masked language modeling is a conservative underestimate of the true gap in perplexities; the real gap can only be larger.

## 4 Alignment dramatically improves protein fitness predictive power

We train three model sizes — 1.2B, 12B, and 102B — for production-ready inference. To demonstrate alignment capabilities, we test our most lightweight production-ready model on an adapted tertiary coordination task proposed by [28], wherein we examine the ability of Odyssey to predict the fitness of generated proteins toward a multi-objective end.

Six held-out enzymes were curated from the Mechanism and Catalytic Site Atlas [60]. We prompt the pre-trained Odyssey-1.2B with a total of 16 generation prompts per enzyme, in which active site residues were kept unmasked as both sequence and structure context. The length of the protein was randomly sampled to be between 200-600 residues, with the inter-active-site sequence distances kept constant between masks of a given protein. Apart from the active site, the remaining residue positions are masked, and each prompt is used to generate 128 sequences, for a total of 2048 generated sequences per protein. Each protein is then folded and scored by pTM (predicted TM-score) and cRMSD (constrained-site RMSD). Collectively, these two scores answer the questions of global fold quality (i.e., as a proxy for the ability for the protein to successfully fold *in vitro*) and active site conformation, both important yet independent objectives in enzyme design.

The true score was computed from these two inputs for each generated sequence:

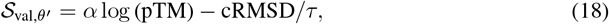

where *α* and *τ* were tuned for the relative weighting of pTM and cRMSD.

Alignment following the methods outlined in Section I was performed using the true scores computed above to construct preference pairs. After alignment using D2-DPO, we observe that the model is able to learn strong positive relationships correlating the model’s internal predicted score 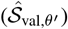 with the true score 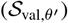, turning into a powerful predictor of *de novo* generated enzyme fitness. Remarkably, this emerges without explicit pretraining on tertiary-coordination or fitness prediction, underscoring out-of-distribution generalization from the model’s internal priors. These results suggest that the latent representation captures hierarchical protein “grammar” — sequence–structure–function constraints — that D2-DPO can surface and align to downstream fitness.

## 5 Discussion

We presented the Odyssey family of protein language models, which ranges in size from 35 million to more than 102 billion parameters. Odyssey advances protein generative modeling along three axes — representation, mechanism, and training objective. Our results suggest these decisions reinforce one another. Odyssey comprises a finite scalar quantizer for high-fidelity discretization of atomic coordinates and a transformer stack for multimodal learning. The FSQ avoids the codebook dynamics and auxiliary losses typical of VQ-VAEs, while supporting stage-wise training that first reconstructs 3-backbone coordinates and then atom-14 coordinates with a frozen encoder. In practice, this delivers a stable structure vocabulary and simplifies joint embedding with sequence; in our empirical results, we find state-of-the-art performance of our FSQ as a protein structure tokenizer.

Within the transformer stack, *consensus*, a content-adaptive local propagation rule, replaces global attention. By propagating agreement over a sparse neighborhood graph using an anisotropic energy function in feature space, consensus selects feature directions in which to accelerate mixing while preserving differences in the complementary directions. Furthermore, the anisotropy depends on the content, allowing for more agreement in homogeneous regions while preserving sharp boundaries as required across the sequence. Two observations matter for scale:

1. 𝒪(*L*) cost enables longer contexts, laying the groundwork for longer, more complex protein generation tasks and/or faster parallel exploration of many potential generated protein conformations.
2. The interval of near-optimal learning rates, 𝒲, diminishes more gradually for consensus versus attention as model size increases. Practically, this widens the operating envelope for large protein models, where exhaustive sweeps are infeasible, without sacrificing downstream performance. It similarly enables more aggressive learning rate schedules for faster training convergence.

We train Odyssey with a discrete diffusion learning objective and generation scheme that more closely mirrors evolution itself (under the coarse ‘proposal and selection’ model), which we empirically show improves validation perplexity under corruption schedules that reflect the reverse-time denoising task. Due to the DWDSE loss being an upper bound, we note this measured gap is conservative.

Altogether, these strategies admit powerful data-efficient scaling properties relevant to the training and generative capabilities of protein models, while enabling post-hoc alignment that surfaces latent sequence–structure–function constraints for both generalizable and specialized task performance.

## Acknowledgments

We thank Vishvajit Kher, Alex Lourenco, and the rest of the Andromeda AI cluster team for their help with wrangling GPUs, and we thank Gustaf Alströmer, Dan Fitzgerald, and Claire Goldsmith among others for their overarching support. We also thank the experts who reviewed and gave feedback on Odyssey and this manuscript prior to release.

## Competing Interests

All authors are current or former employees, executives, and/or directors of Anthrogen, PBC, and may hold shares in Anthrogen, PBC. Patents have been filed related to aspects of this work.

## Safety Considerations

Protein foundation models are inherently dual-use: the same capabilities that advance human health, sustainability, and basic science could, in principle, be misapplied. Our work is informed by the [Principles for the Responsible Development of AI for Biological Design](https://responsiblebiodesign.ai/), and we support continued community development of evaluations, screening standards, and access norms as capabilities evolve. With these considerations in mind, we find the benefits of releasing this research publicly far outweigh the potential risks.

## Appendix

### A Finite scalar quantizer benchmarks

We evaluate the reconstruction fidelity of our finite scalar quantizers (FSQs) (see Section F) trained in two stages (stage-1, which utilizes a lightweight encoder and decoder, and stage-2, where the stage-1 encoder is frozen while training a large decoder). Reconstructions are assessed by RMSD on CAMEO [26], CASP15 [39], and CASP16 [82] for the (a) 3-backbone and (b) atom-14 representations. Figure 8 summarizes these evaluations.

**Figure 7:**
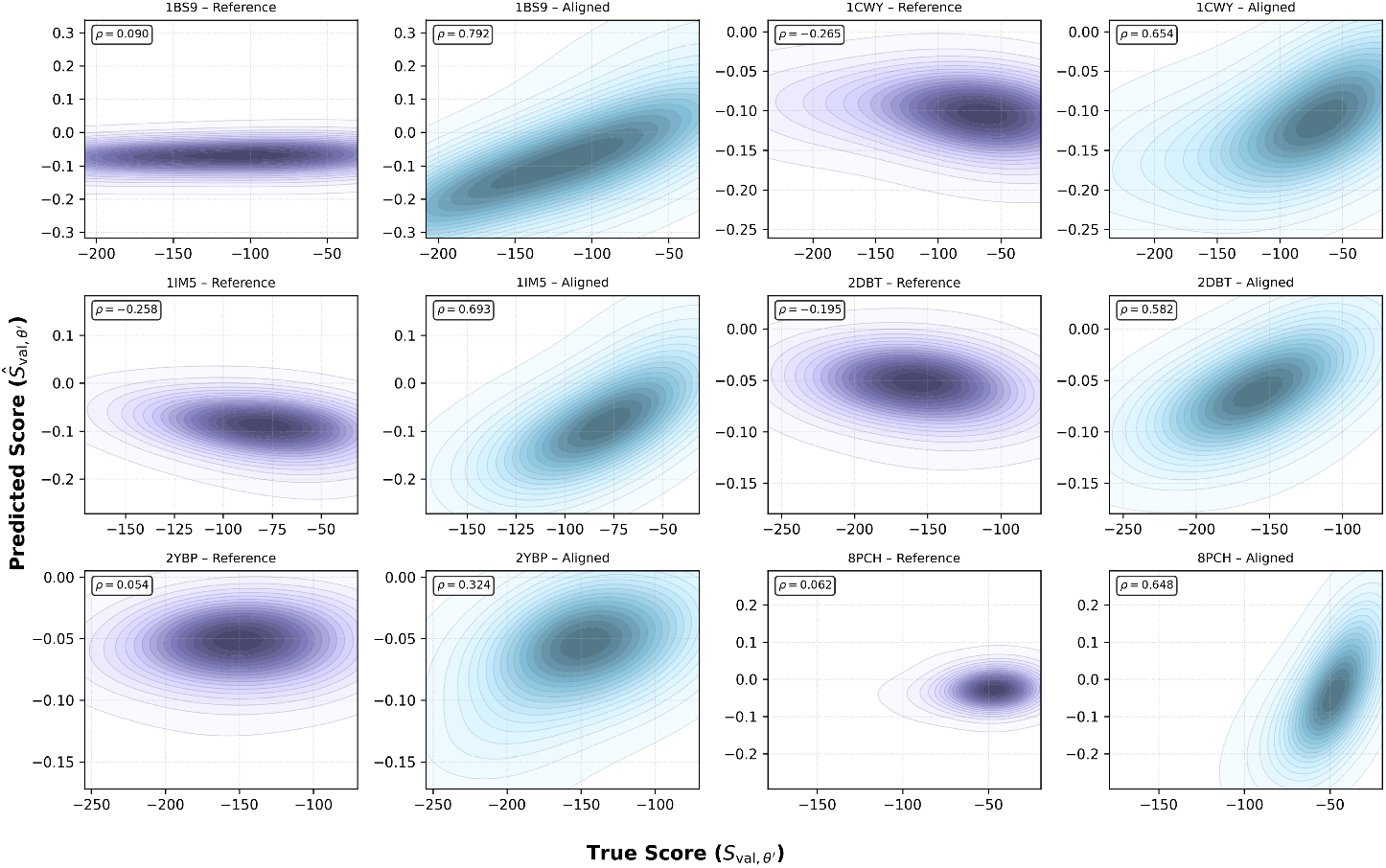
Alignment performance on six held-out enzymes from M-CSA. We observe that the model has aligned to the true fitness score (measured by Spearman correlation coefficient, *ρ*)

**Figure 8:**
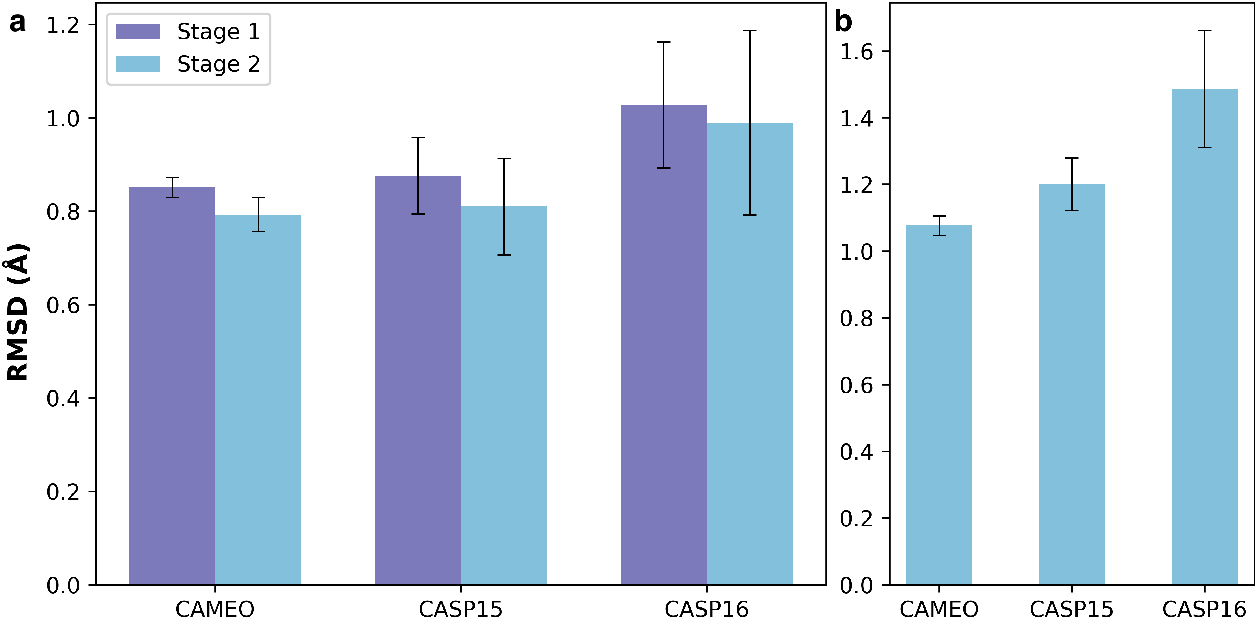
Benchmarking FSQ reconstruction performance (RMSD) on CAMEO, CASP15 and CASP16. (a) 3-backbone reconstruction performance of stage-1 and stage-2 FSQ. (b) Atom-14 reconstruction performance of stage-2 FSQ.

We observe that our stage-1 and stage-2 FSQs achieve state-of-the-art 3-backbone reconstruction performance on CAMEO, CASP15, and CASP16, with the stage-2 FSQ further achieving state-of-the-art atom-14 reconstruction performance on CAMEO, CASP-15, and CASP-16. We further visualize twelve randomly selected proteins from the CAMEO dataset in Figure 9 to illustrate the FSQ’s ability to learn and retain structure at an atomic scale, even without a learned codebook.

**Figure 9:**
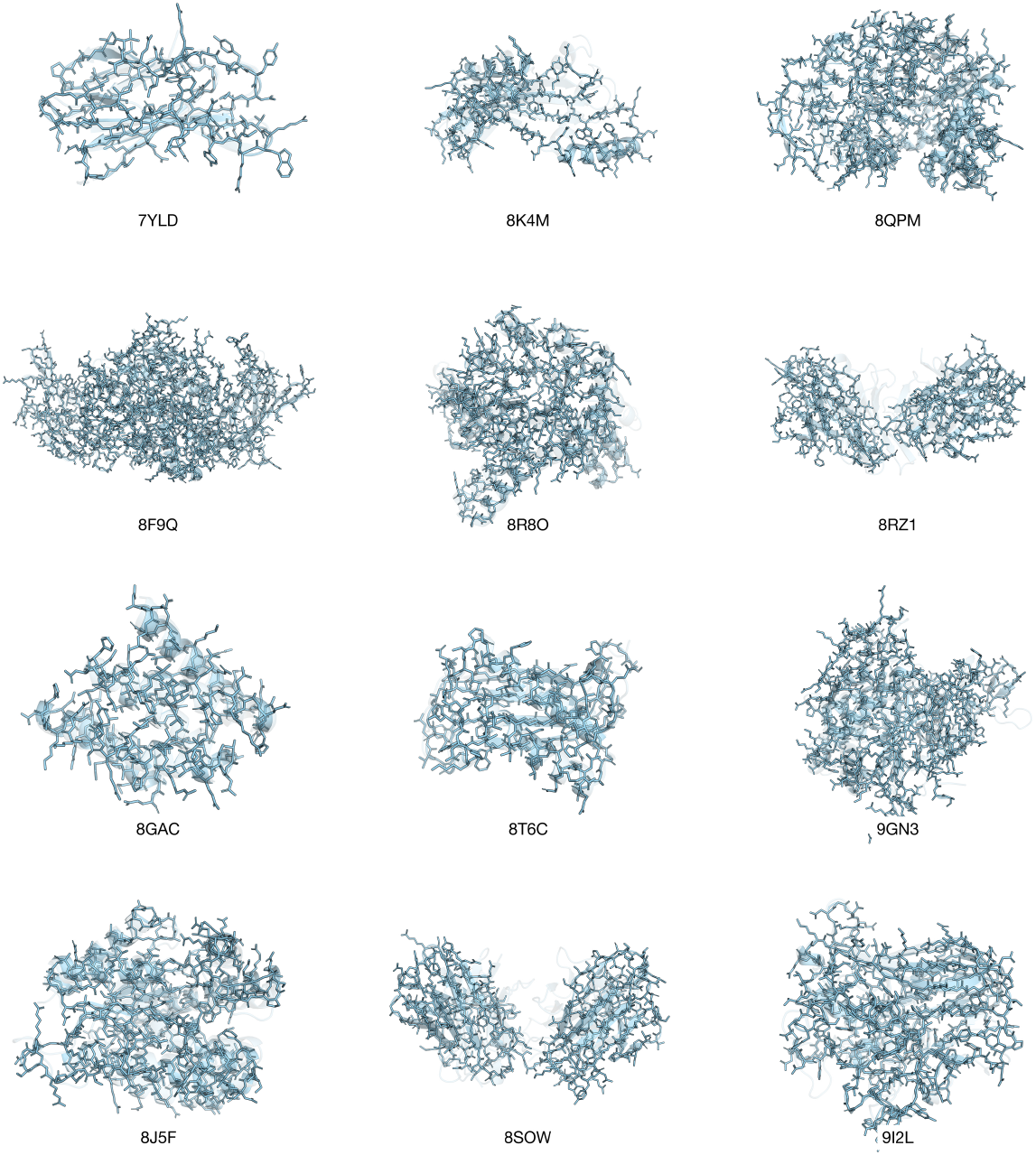
Sample reconstruction visualizations of the FSQ on the CAMEO dataset. The cartoon representation of the original protein is shown in gray with the cartoon and stick representations of the reconstructed structures after stage-2 training shown in light blue.

#### B Data details

We outline the per-track composition of the Odyssey pre-training dataset in the following sections. The pre-training dataset is sourced from UniRef [1], MERC [69], SRC [69], AntiRef [14], OMG [19], PDB, ESMAtlas [41], and AlphaFoldDB [33], as detailed in Table 2.

**Table 2:**
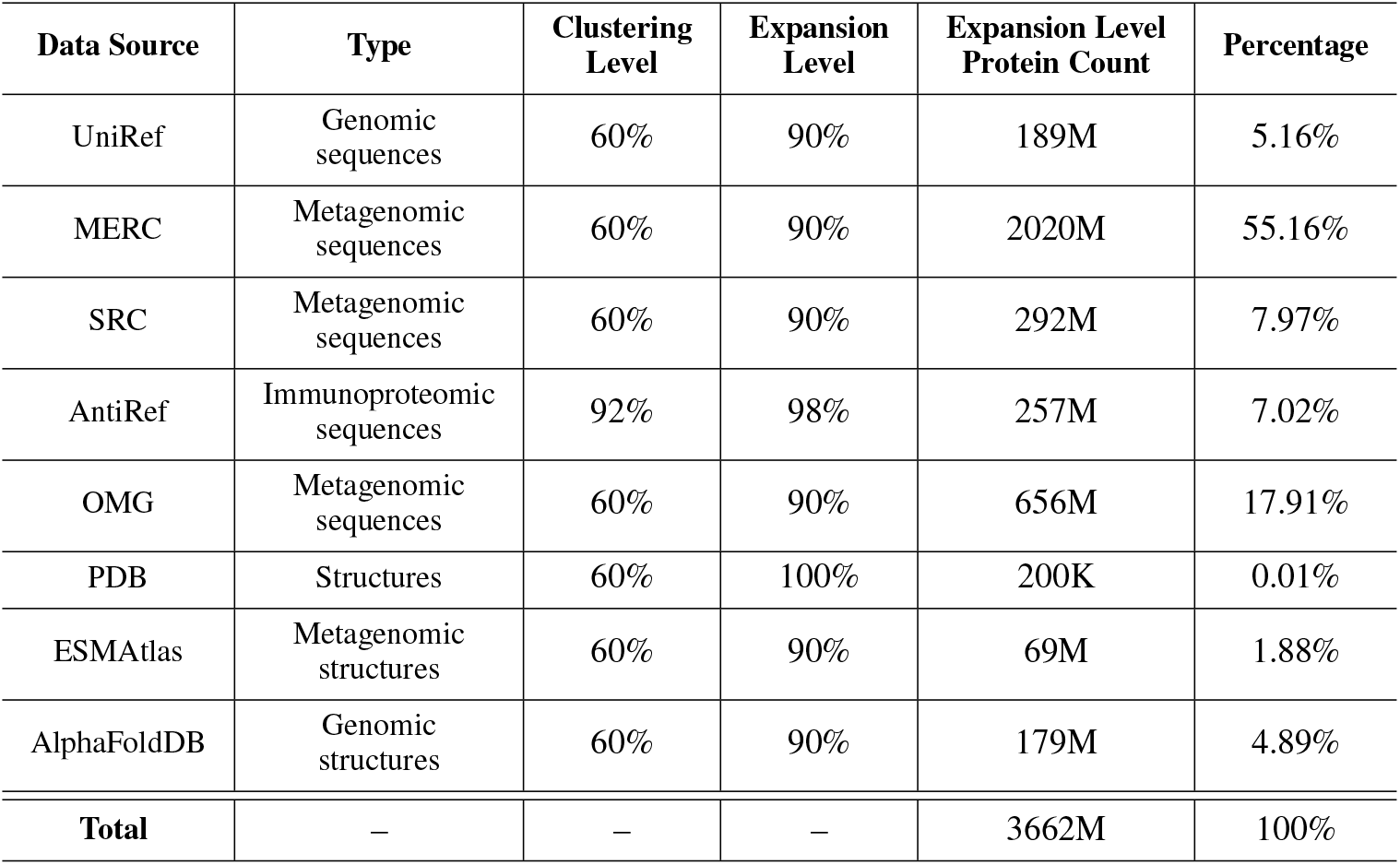
Composition of the Odyssey pre-training dataset.

#### B.1 Sequence

The sequences in Table 2 were downloaded and parsed with a strict cutoff date of January 1st, 2024 applied to all sequence data sources (the OMG dataset [released in 2024] solely comprises sequences from the JGI’s IMG database [46] and EMBL’s MGnify database [61]), with the latest snapshots occurring in August 2023 [19].

The sequences corresponding to the structural data from Section B.2 were also read and parsed. For pre-training, a hierarchal data strategy was utilized whereby all sequences (bar the antibodies and PDB structures) were clustered at 90%. These cluster representatives were then clustered down to 60%. The antibody sequences were clustered at 98% and 92%, respectively (to maintain CDR diversity), and the PDB structures were clustered at 100% and 60% due to their high quality, and the relatively low number of structures in this database. During pre-training, for each protein in the data loader, one of the 60% clusters was chosen with *n/*(1 + log(*n*)) probability (where *n* denotes the number of 90% clusters within the 60% cluster), and from this cluster, one of the 90% cluster representatives was randomly chosen. This approach balanced diversity with a controlled bias toward heavily represented protein families. All clustering was done with DIAMOND [15] linclust.

#### B.2 Atomic Coordinates

The structures in Table 2 were downloaded and parsed with strict cutoff date Aug 1st, 2022, enabling temporal holdout of the CASP and CAMEO datasets for FSQ benchmarking. For the PDB data, we used all structures prior to the cutoff date and with ≤ 5Å resolution, irrespective of imaging process. For FSQ training, all structures were clustered at 40%, with these representatives comprising the training dataset. During Odyssey pre-training, the clustering strategy detailed in B.1 was observed.

#### B.3 Orthologous groups, semantic descriptions, and per-residue domain annotations

Orthologous groups, semantic descriptions, and per-residue annotations comprise the set of ‘functional’ context tracks that Odyssey was pre-trained on. They were generated with eggNOG-mapper-v2 using DIAMOND as the underlying algorithm [16, 15, 32]. We used this approach for all expansion level proteins besides antibodies. The data was processed and tokenized via the scheme outlined in Sections C.5, C.6, and C.7.

#### B.4 Secondary structure and solvent-accessible surface area

Secondary structure was predicted utilizing DSSP 4 [29] for all proteins with an associated tertiary structure. Solvent-accessible surface area (SASA) was computed with the *biotite* python package [40] (based on the Shrake-Rupley algorithm [65]).

### C Tokenization

This section outlines the tokenization schemes leveraged for the eight data *tracks* comprising Odyssey. We reserve a track-agnostic inventory of special symbols used across all Odyssey tracks for corruption-based training and sequence delimitation, where *S* = |𝒱_special_| = 6. Accordingly:

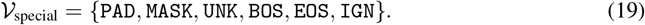

All track-specific content vocabularies, 𝒱, where *V* = |𝒱|, occupy the lowest indices in the global vocabulary, while the *S* special tokens are offset to occupy the range {*V*, …, *V* + *S* − 1}. Tracks are wrapped with `BOS` and `EOS` and then padded with `PAD` to their maximum length.

We distinguish *source* tracks (sequence and 3D atomic structure coordinates) from auxiliary *context* tracks (secondary structure, SASA, pLDDT, orthologous groups, semantic descriptions, and domains). Sequences are tokenized over a fixed 25-symbol alphabet (see Section C.1), and atomic coordinates are represented by 3-backbone, atom-14, or atom-37 layouts (see Section C.2) and discretized via finite scalar quantization (see Section F.2). Secondary structure is tokenized via a finite inventory (see Section C.3), SASA is discretized via quantile binning (see Section C.4), orthologous groups are formed via at most 32 OG-tags drawn from a filtered vocabulary (see Section C.5), free-text semantic descriptions are tokenized into a word-level vocabulary and truncated to 512 tokens (see Section C.6), domain annotations supply at most 4 per-residue tags from a frequency-filtered inventory (see Section C.7), and pLDDT is discretized via uniform binning (see Section C.8).

#### C.1 Sequence

We tokenize amino-acid sequences over a fixed alphabet where *V*_seq_ = |𝒱_seq_| = 25. Accordingly:

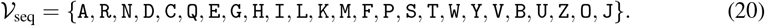

The sequence is a *source* track, where after tokenization and embedding, it is positionally summed with the tokenized and embedded structure track, and passed to the transformer stack.

##### Data dimensions

We denote the original sequence as *x*_seq_ ∈ {0, …, *V*_seq_ – 1} ^*B×L*^ and the tokenized sequence as 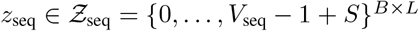, where *B* is the batch size (number of proteins in the batch), and *L* is the maximum sequence length (2048 total tokens, up to 2046 content tokens).

#### C.2 Atomic coordinates

We tokenize continuous atomic structure coordinates into discrete structure tokens using a finite scalar quantizer, yielding a vocabulary of size *V*_struct_ = |𝒱_struct_| = 4375. We outline this tokenization methodology in Section F.2. As a precursor, we discuss how we can efficiently and unambiguously record the positions of atoms in a protein, wherein we invoke a flattened batch with *B* = 1.

A protein can be rotated and translated arbitrarily in a given reference frame. For a protein containing *L* residues, suppose that we fix an arbitrary position and orientation. Accordingly, each atom can be represented by its Cartesian coordinates in ℝ^3^. The three primary backbone (“3-backbone”) atoms in each amino acid are *N, C*_*α*_, and *C*.

##### Definition C.1

(3-Backbone). *The 3-backbone representation of a residue corresponds to positions of its N, C*_*α*_, *and C atoms, and its representation is in* ℝ^3*×*3^.

Representing only these backbone atoms, we denote a residue, *i* ∈ *{*0, …, *L* − 1*}*, 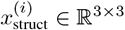 and the protein via *x*_struct_ ∈ ℝ^*L×*3*×*3^. In contrast to the backbone, representing side chains is more complex, since the number of atoms in a side chain is not fixed across amino acids. Moreover, even among amino acids with the same number of side-chain atoms, the constituent atom types may differ (for instance, cysteine differs from serine by replacing the hydroxyl group with a thiol). Furthermore, *even when both the number and types of atoms coincide*, their spatial arrangement can vary (as in the case of isoleucine and leucine, which differ only by a rearrangement of identical atoms).

Atom-14 [33] is one proposed method of recording the 3-backbone and heavy (i.e. non-hydrogen) side chain atoms. In atom-14, the atoms of a side chain are ordered via their proximity to the *C*_*β*_ atom. For example, the side-chain atoms can be ordered as {*C*_*γ*_, *S*_*δ*_, *C*_*ϵ*_} in methionine and {*C*_*γ*_, *C*_*δ*_, *C*_*ϵ*_, *N*_*ζ*_} in lysine. The largest amino acid, tryptophan, has three atoms in its 3-backbone and 11 heavy atoms in its side chain (for a total of 14 atoms). We denote a residue, *i* ∈ *{*0, …, *L* − 1*}*, as 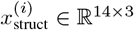 and a protein as *x*_struct_ ∈ ℝ^*L×*14*×*3^. For a residue with *K* atoms in its side chain, the final (14 *−* 3 *− K*) rows of *x*_struct_ can be set to all-zero coordinates. The central issue with the atom-14 representation is that distinct amino acid types have the same representation (or at least representations with all-zero coordinates in identical places — e.g., cysteine versus serine and leucine versus isoleucine). Thus, the side-chain cannot be identified from *x*_struct_ ∈ ℝ^*L×*14*×*3^ alone; in addition we require sequence information to uniquely determine the elemental identity of the side-chain atoms.

An alternative representation is the atom-37 scheme introduced by [33], which encodes side chains through a fixed vocabulary of canonical atom labels. While any given residue contains at most 11 heavy side-chain atoms, the union across the 20 amino acids comprises 34 distinct labeled heavy side-chain sites (e.g., 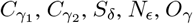. An order is fixed to the 37 total sites (three 3-backbone and 34 heavy side-chain). For any amino acid, all 3-backbone and heavy side-chain are recorded in a position pertaining to their site. We denote a residue, *i* ∈ *{*0, …, *L* − 1*}*, as 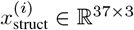 and a protein as *x*_struct_ ∈ ℝ^*L×*37*×*3^. Vacant atomic sites can be set to all-zero coordinates. While atom-37 is more data-intensive than atom-14, the ambiguity in identifying amino acids is resolved.

Accordingly, to fully represent the 3-backbone and heavy side chain atoms of a protein, we must invoke either the atom-37 representation or atom-14-plus-sequence representation. We note that the complete sequence of a protein can be obtained from the atom-37 coordinates (moreover, it can be obtained solely from the position of the all-zero coordinates). As an example, if an atom-37 tensor in ℝ^*L×*37*×*3^ contains non-zero coordinates in the row corresponding to *S*_*γ*_, then it must be cysteine, as no other residue has an *S*_*γ*_ atom. Similarly, leucine and isoleucine can be distinguished based on whether the all-zero coordinates correspond to 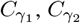, and 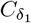, or to 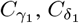, and 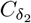. In atom-14, the full sequence of a protein can only be *partially* obtained from the coordinates provided. As an example, if a residue has all 14 sites provided, it cannot be aspartic acid (which has 3 side chain atoms), and if a residue only has 4 side chain atoms provided, it cannot be proline (which has 2 side chain atoms). However, residues with the same number of side chain atoms cannot be distinguished using atom-14. We summarize these representation schemes, including 3-backbone, in Table 6.

This raises a methodological challenge: we aim to treat sequence and structure as distinct modalities, enabling us to apply `MASK` to one while leaving the other fully observed. However, if either modality is uniquely determined by the other, the masking task becomes degenerate, undermining the rationale for maintaining separate modalities. We explore this further in Section G.1.

##### Data dimensions

We write the original structure as *x*_struct_ ∈ ℝ^*B×L×A×*3^, and tokenized structure as 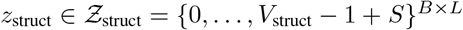, where *B* is batch size (number of proteins in a batch), *L* is the maximum sequence length (2048 total tokens, up to 2046 content tokens), *A* denotes the number of sites within each residue (can be 3, 14, or 37 for 3-backbone, atom-14, or atom-37 representations), and 3 denotes the number of coordinates comprising each atom in Cartesian space.

#### C.3 Secondary structure

We adopt the eight-state secondary structure (SS) inventory (C, E, B, T, S, H, G, I) and extend it with polyproline II (P), yielding *V*_ss_ = |𝒱_ss_| = 9:

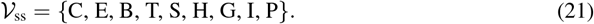

The secondary structure is a *context* track, where after tokenization and embedding, it cross-attends to the tokenized, embedded, and positionally summed sequence and structure tracks.

##### Data dimensions

We denote the original secondary structure as *x*_ss_ ∈ {0, …, *V*_ss_ *−* 1} ^*B×L*^ and its tokenized version as 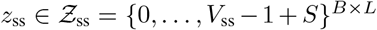 where *B* is batch size (number of proteins in a batch), and *L* is the maximum sequence length (2048 total tokens, up to 2046 content tokens).

#### C.4 Solvent-accessible surface area (SASA)

We discretize per-residue solvent-accessible surface area (SASA), *s*_*t*_ *∈* ℝ^+^, into quantile-based bins with approximately equal population, using thresholds estimated from a uniformly sampled subset of the Odyssey training dataset, wherein *V*_sasa_ = |𝒱_sasa_| = 16. Accordingly:

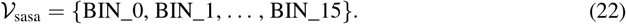

The solvent-accessible surface area is a *context* track, where after tokenization and embedding, it cross-attends to the tokenized, embedded, and positionally summed sequence and structure tracks.

##### Data dimensions

We denote the original SASA as 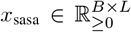 and the tokenized SASA as 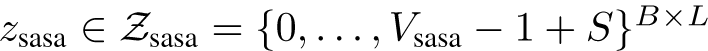, wherein *B* is the batch size (number of proteins in the batch), and *L* is the maximum sequence length (2048 total tokens, up to 2046 content tokens).

#### C.5 Orthologous groups

We construct an orthologous group vocabulary from a uniformly sampled subset of the Odyssey training set by counting OG-tag occurrences and retaining only those that appear at least 10 times, where *V*_*og*_ = |𝒱_*og*_|. For each protein, the orthologous group comprises a maximum of 32 OG-tags, each denoted as *og*.

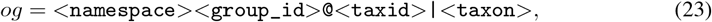

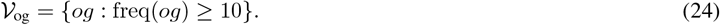

The orthologous groups track is a *context track*, where after tokenization and embedding, it cross-attends to the tokenized, embedded, and positionally summed sequence and structure tracks.

##### Data dimensions

We denote the original OG-tags as *x*_*og*_ ∈ *{*0, …, *V*_*og*_ *−* 1} ^*B×G*^, and the tokenized OG-tags as 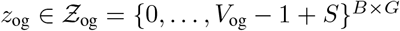, where B is batch size (number of proteins in a batch), and G is the maximum number of OG-tags (32 total tokens, up to 30 content tokens).

#### C.6 Semantic descriptions

We construct a word-level vocabulary from a uniformly sampled subset of the Odyssey training set by splitting free-text semantic descriptions on spaces and counting word frequencies, retaining only words that appear at least 5 times, where *V*_sd_ = |𝒱_sd_|. Each protein’s description is then represented as a sequence of space-delimited words, w, with length constrained to the first 512. Accordingly:

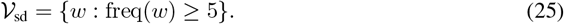

The semantic descriptions track is a *context* track, as after tokenization and embedding, it cross-attends to the tokenized, embedded, and positionally summed sequence and structure tracks.

##### Data dimensions

We write the original descriptions as *x*_sd_ ∈ {0, …, *V*_sd_ *−* 1} ^*B×H*^ and the tokenized descriptions as *z*_sd_ ∈ {0, …, *V*_sd_ 1 + *S*} ^*B×H*^, where B is the batch size (number of proteins in the batch), and *H* is the maximum number of descriptions (512 total tokens, up to 510 content tokens).

#### C.7 Per-residue domain annotations (Domains)

We represent domains as per–residue categorical annotations, where *V*_dom_ = |𝒱_dom_|. From a uniformly sampled subset of the Odyssey training set, we recorded a histogram of domain tags, *dom*, and retained only those with corpus frequency at least 4, forming the vocabulary

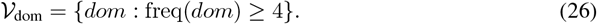

Each residue carries up to four in-vocabulary domain annotations. The domains track is a *context* track, where after tokenization and embedding, it cross-attends to the tokenized, embedded, and positionally summed sequence and structure tracks.

##### Data Dimensions

We write the original domains as *x*_dom_ ∈ {0, …, *V*_dom_ *–* 1} ^*B×L×K*^ and tokenized domains as 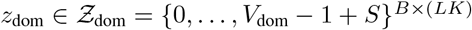, where *B* is the batch size (number of proteins in the batch), L is the maximum sequence length (2048 total tokens, up to 2046 content tokens), *and K* is the maximum number of domain tags per residue (4 [max] content tokens).

#### C.8 Structure confidence scores (pLDDT)

We discretize per-residue structure confidence scores (pLDDT), *p*_*t*_ ∈ [0, 1], into uniform bins, wherein *V*_plddt_ = |𝒱_plddt_| = 50. Higher *p*_*t*_ values monotonically indicate greater confidence in the correctness of the residue’s local geometry (its placement relative to nearby atoms). Accordingly:

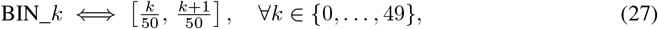

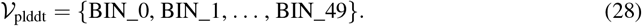

The pLDDT track is a context track, where after tokenization and embedding, it cross-attends to the tokenized, embedded, and positionally summed sequence and structure tracks.

##### Data dimensions

We write the original pLDDT as *x*_*plddt*_ ∈ [0, 1]^*B×L*^ and the tokenized pLDDT as 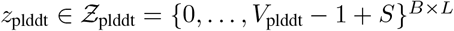, where *B* is batch size (number of proteins in a batch), and *L* is the maximum sequence length (2048 total tokens, up to 2046 content tokens).

### D Consensus

As we describe in Section F, the Odyssey architecture comprises transformer blocks, which leverage the *consensus* mechanism. Consensus is a drop-in replacement for attention, and seeks the same goal of cross-token communication, accomplishing it iteratively through repeated local *agreement* on a sparse graph. Each token is connected to a set of neighboring tokens, defined using a local window, and we propagate to nearby windows with every forward pass through the mechanism. Domains such as protein structures are often governed by local coherence, since nearby residues tend to agree. By casting this as a consensus problem where each token iteratively negotiates with neighbors, we embed the notion that the representation must be locally consistent before global propagation. We outline self-consensus in Section D.1, cross-consensus in Section D.2, multi-head consensus in Section D.3, and an adapted version of rotary positional embeddings for consensus in Section D.4. In the following exposition and derivations, we invoke a flattened batch where *B* = 1.

#### D.1 Self-consensus

Consider source embeddings *y* = [*y*^(0)^, …, *y*^(*L−*1)^]^*⊤*^ ∈ ℝ^*L×d*^, with *y*^(*i*)^ ∈ ℝ^*d*^, ∀*i* ∈ *{*0, …, *L* − 1*}*. The self-consensus mechanism produces outputs 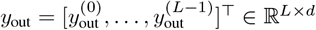, with 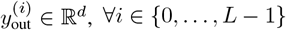. We describe the transformation of *y* into *y*_out_ via self-consensus as follows.

##### D.1.1 Initial projection

We first project the source into the *consensus feature space*, with ⟨*u*^(*i*)^⟩_(0)_ ∈ ℝ^*d*^, ∀*i* ∈ *{*0, …, *L* − 1*}*:

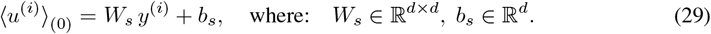

This linear projection maps each source embedding, *y*^(*i*)^, into an initial feature space, ⟨*u*^(*i*)^⟩_(0)_, that will undergo iterative refinement via the self-consensus gradient update in Section D.1.4.

##### D.1.2 Sparse graph construction

We define a directed edge set, *E* ⊂ *{*0, …, *L* − 1*}*^2^, using a *local window*: *E* = *{*(*i, j*) : 0 ≤ *i, j* ≤ *L* − 1, 0 *<* |*i* − *j*| ≤ *w}*. Each embedding has (at most) 2*w* outgoing edges. The outgoing neighbor set of embedding *y*^(*i*)^ is 𝒩^(*i*)^ = *{j* ∈ *{*0, …, *L* − 1*}* : (*i, j*) ∈ *E}*. By restricting to this sparse edge set, each embedding communicates with a limited set of peers rather than the entire sequence at once.

##### D.1.3 Consensus weight matrix

Let ∥ *·* ∥_*R*_ : ℝ^*r×d*^ → ℝ^*r×d*^ denote the row-normalization operator that scales each row of a matrix to unit *ℓ*_2_ norm (but maps zero rows to zero). For each directed edge, (*i, j*) ∈ *E*, we compute:

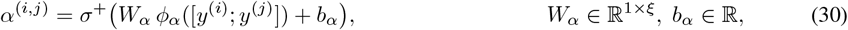

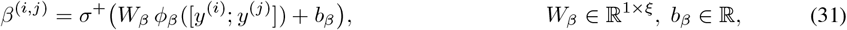

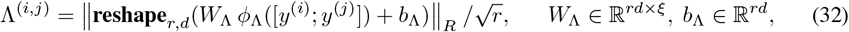

where *α*^(*i,j*)^, *β*^(*i,j*)^ ∈ ℝ_*>*0_, Λ^(*i,j*)^ ∈ ℝ^*r×d*^, *ϕ*_*α*_, *ϕ*_*β*_, *ϕ*_Λ_ : ℝ^2*d*^ → ℝ^*ξ*^ are MLPs, and *σ*^+^ : ℝ → ℝ is softplus activation. Index [*R*^(*i,j*)^]^(*a,b*)^ of the *consensus weight matrix* stores how much the difference in the *b*^th^ feature between embeddings *y*^(*i*)^ and *y*^(*j*)^ informs the update of the *a*^th^ feature of *y*^(*i*)^, with:

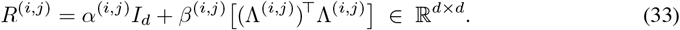

Above, *α*^(*i,j*)^ is a *scalar* weight inducing *isotropic* coupling, while *β*^(*i,j*)^(Λ^(*i,j*)^)^*⊤*^Λ^(*i,j*)^ is a low-rank positive semi-definite *matrix* whose column space identifies up to *r* principal directions of agreement. Thus, *R*^(*i,j*)^ is symmetric positive definite with *λ*_min_(*R*^(*i,j*)^) ≥ *α*^(*i,j*)^. If we omit *α*^(*i,j*)^*I*_*d*_, then *R*^(*i,j*)^ is positive semi-definite, so directions in the null-space of Λ^(*i,j*)^ receive no penalty and the energy loses strict convexity, allowing gradient updates to drift along flat directions. Conversely, a purely isotropic expression, *α*^(*i,j*)^*I*_*d*_, cannot capture structured, *anisotropic* relationships in the feature space. The factorization *β*^(*i,j*)^(Λ^(*i,j*)^)^*⊤*^Λ^(*i,j*)^ yields directional specificity at 𝒪 (*rd*) cost, and in the special case of *r* = 0, the formulation reduces exactly to the isotropic-only coupling, *R*^(*i,j*)^ = *α*^(*i,j*)^*I*_*d*_.

##### D.1.4 Consensus gradient updates

We derive the update by noting that self-consensus seeks to minimize the quadratic energy:

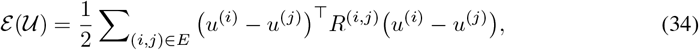

with respect to = 𝒰[*u*^(0)^, …, *u*^(*L−*1)^]^*⊤*^. Because *E* is a directed edge set, the gradient at node *I* receives contributions from its outgoing and incoming edges. The first-order optimality condition is:

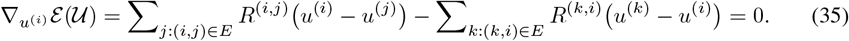

We move toward this optimum via a single explicit gradient descent step:

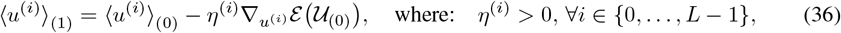

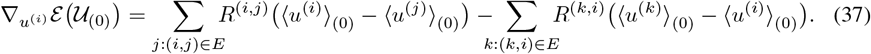

Intuitively, node *i* exchanges its current “disagreement” with each neighboring node, and moves by a weighted average of these differences. Because each *R*^(*i,j*)^ is symmetric positive definite, ℰ is convex and has a translation invariance obtained by adding the same vector, *c* ∈ ℝ^*d*^, to every node (i.e., the Hessian has rank *Ld − d*). Accordingly, ℰ is strictly convex on the orthogonal complement of this consensus subspace. With a suitable learned step size *η*^(*i*)^, the update decreases ℰ, and can accelerate mixing in smooth neighborhoods (large *η*^(*i*)^), while inhibiting it near sharp boundaries (small *η*^(*i*)^).

##### D.1.5 Output projection

Following the gradient update, we map back to the input space, where 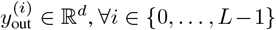:

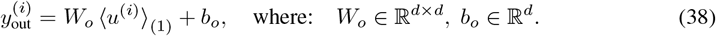

The aforementioned output is given by *y*_out_ ∈ ℝ^*L×d*^. Per edge, self-attention uses a *scalar* weight, while self-consensus uses a *matrix, R*^(*i,j*)^. Regarding complexity, the initial and output projections cost *O*(*Ld*^2^), computing all edge parameters once costs *O*(*Ldwrξ*), and the gradient update costs 𝒪(*Ldwr*). Thus, the overall complexity of the self-consensus mechanism is 𝒪(*Ld*^2^ + *Ldwrξ*).

#### D.2 Cross-consensus

Consider source embeddings *y* = [*y*^(0)^, …, *y*^(*L−*1)^]^*⊤*^ ∈ ℝ^*L×d*^ with *y*^(*i*)^ ∈ ℝ^*d*^, ∀*i* ∈ *{*0, …, *L* − 1*}*, and context embeddings *c* = [*c*^(0)^, …, *c*^(*J−*1)^]^*⊤*^ ∈ ℝ^*J×d*^ where *c*^(*i*)^ ∈ ℝ^*d*^, ∀*i* ∈ *{*0, …, *J* − 1*}*. The cross-consensus mechanism produces outputs 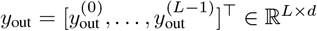, with 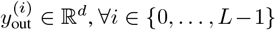, ∀*i* ∈ {0, …, *L* − 1}, imbuing the context embeddings into the source embeddings. We describe the transformation of *y* and *c* into *y*_out_ via cross-consensus as follows.

##### D.2.1 Initial projection

We first project the embedded source and context into the consensus feature space, with ⟨ *u*^(*i*)^⟩_(0)_ ∈ ℝ^*d*^, ∀*i* ∈ *{*0, …, *L* − 1*}*, and ⟨*v*^(*j*)^⟩ ∈ ℝ^*d*^, ∀*j* ∈ *{*0, …, *J* − 1*}*:

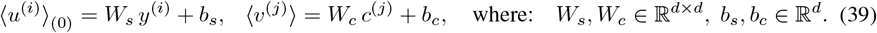

This linear projection maps each source embedding, *y*^(*i*)^, into an initial feature space, ⟨*u*^(*i*)^⟩_(0)_, that will undergo iterative refinement through the cross-consensus gradient update in Section D.1.4. The projection, ⟨*v*_*j*_ ⟩, serves as the fixed context for the cross-consensus gradient update.

##### D.2.2 Sparse graph construction

We define a directed bipartite edge set, *E*_×_ ⊂ {0, …, *L –* 1} *× {*0, …, *J −* 1}, via a local window. If the source and context embeddings are index-aligned (the ordering of embeddings in *y* and *c* refers to the same underlying axis), then (*i, j*) ∈ *E*_*×*_ whenever |*i − j*| ≤ *w* and *i* ≠ *j*. This local-window rule is *only valid under such index alignment*. Otherwise, locality in one sequence is not meaningful in the other. The outgoing neighbor set of embedding *y*^(*i*)^ is 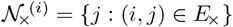.

##### D.2.3 Consensus weight matrix

Let ∥ *·* ∥_*R*_ : ℝ^*r×d*^ → ℝ^*r×d*^ denote the row-normalization operator that scales each row of a matrix to unit *ℓ*_2_ norm (mapping zero rows to zero). For each directed edge, (*i, j*) ∈ *E*_*×*_, we compute:

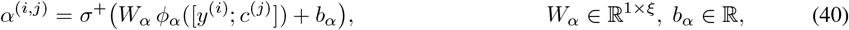

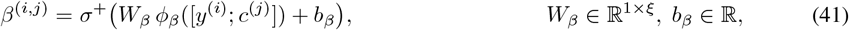

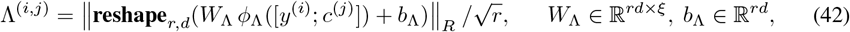

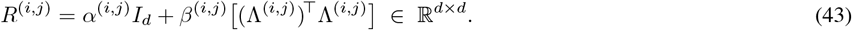

The construction of the consensus weight matrix, *R*^(*i,j*)^, parallels that of Section D.1.3. *R*^(*i,j*)^ conveys how strongly the source embedding *y*^(*i*)^ should “agree” with the context embedding *c*^(*j*)^.

##### D.2.4 Consensus gradient updates

Paralleling Section D.1.4, we note that cross-consensus seeks to minimize the quadratic energy:

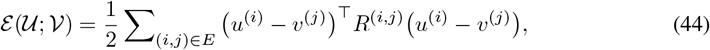

with respect to 𝒰 = [*u*^(0)^, …, *u*^(*L−*1)^]^*⊤*^, where 𝒱 = [*v*^(0)^, …, *v*^(*J−*1)^]^*⊤*^ is held fixed. Because *E*_*×*_ is directed from source to context, the gradient at source node *i* receives contributions only from its outgoing edges. The first-order optimality condition is:

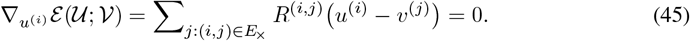

We move toward this optimum via a single explicit gradient descent step:

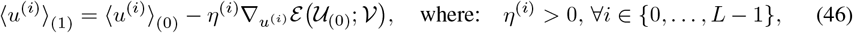

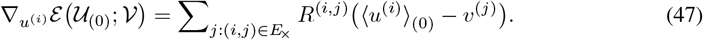

As in Section D.1.4, source node *i* exchanges current disagreements with context neighboring nodes, and moves by a weighted average of the differences. Since each *R*^(*i,j*)^ is symmetric positive definite and the context is held fixed, ℰ is strictly convex in 𝒰, wherein there is a unique minimizer and no translation invariance as in self-consensus. With a suitable step size, *η*^(*i*)^, this update decreases ℰ.

##### D.2.5 Output projection

Following the gradient update, we map back to the input space, where 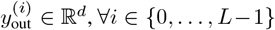:

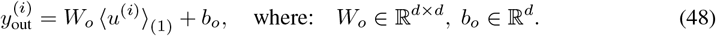

The aforementioned output is given by *y*_out_ ∈ ℝ^*L×d*^. Regarding complexity, the initial projections cost *O*((*L* + *J*)*d*^2^), the output projection costs *O*(*Ld*^2^), computing all edge parameters once costs *O*(*Ldwrξ*), and the gradient update costs 𝒪(*Ldwr*), yielding 𝒪((*L* + *J*)*d*^2^ + *Ldwrξ*) in total.

#### D.3 Multi-head consensus

Mirroring multi-head attention [74], consensus can be run with *H* parallel subspaces. Let *d* = *Hd*_*H*_. For self-consensus, each projected source embedding in the consensus feature space is given by Eq. (49), ∀*i* ∈ *{*0, …, *L* − 1*}*, and ∀*h* ∈ *{*0, …, *H* − 1*}*:

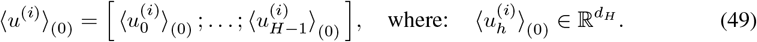

For cross-consensus, we additionally have each projected context embedding in the consensus feature space, which is given by Eq. (50), ∀*j* ∈ *{*0, …, *J* − 1*}*, and ∀*h* ∈ *{*0, …, *H* − 1*}*:

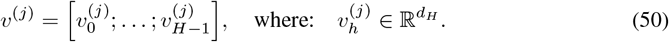

For every edge, (*i, j*), in the directed graph (*E* for self-consensus and *E*_*×*_ for cross-consensus), and for every head, *h*, we obtain the scalar, *α*^(*i,j*)^, and the low-rank matrix, 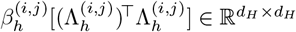 (see Section D.1.3 and Section D.2.3), wherein:

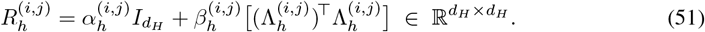

Each head performs the consensus gradient update on its own quadratic energy. We provide separate update rules for the two consensus modes. For self-consensus, we have that, ∀*h* ∈ *{*0, …, *H* − 1*}*:

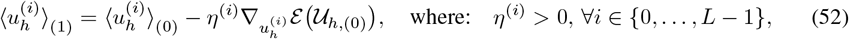

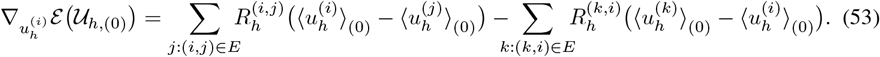

For cross-consensus, we have that, ∀*h* ∈ *{*0, …, *H* − 1*}*:

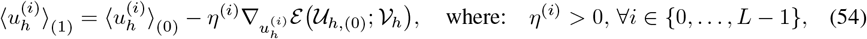

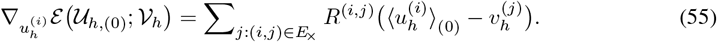

Following the update, the *H* head outputs are concatenated and mapped back to the input space:

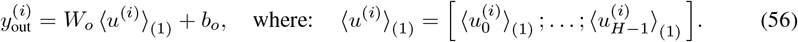

Since the *H* heads are processed in parallel, the cost of multi-head self-consensus is 𝒪(*Ld*^2^ +*Ldwrξ*) and of multi-head cross-consensus is 𝒪(*L* + *J*)*d*^2^ + *Ldwrξ*). Multi-head consensus thus preserves linear complexity in *L* while enabling different heads to capture complementary notions of agreement.

#### D.4 Rotary positional embeddings

In both self-consensus and cross-consensus, we inject absolute position information by rotating each endpoint of every edge before forming the edge difference. Suppose *d* is even, and consider a base, *B >* 0. For position index, *p* ∈ ℕ, we define the angle vector ***θ***(*p*) ∈ ℝ^*d/*2^, where:

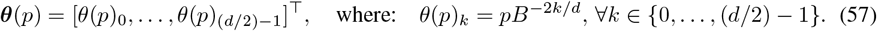

Let cos(***θ***(*p*)), sin(***θ***(*p*)) ∈ ℝ^*d*^ delineate the elementwise cosine and sine of ***θ***(*p*) repeated to length *d*. Define 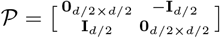 as the fixed permutation-sign matrix that exchanges two half-vectors. We now outline rotary positional embeddings (RoPE) [71] for self and cross-consensus.

##### D.4.1 RoPE for self-consensus

Before the consensus gradient update step (see Section D.1.4), for each directed edge (*i, j*) ∈ *E*, we rotate each endpoint, ⟨*u*^(*i*)^⟩_(0)_, ⟨*u*^(*j*)^⟩_(0)_, wherein:

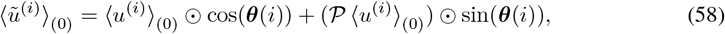

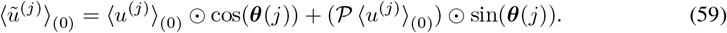

The edge disagreement from the consensus gradient update step is now 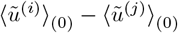 in place of ⟨*u*^(*i*)^⟩_(0)_ − ⟨*u*^(*j*)^⟩_(0)_, whereby we consider 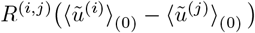.

##### D.4.2 RoPE for cross-consensus

Paralleling the self-consensus case, the consensus gradient update step (see Section D.2.4), for each directed edge (*i, j*) ∈ *E*_*×*_, we rotate each endpoint, ⟨*u*^(*i*)^⟩_(0)_, ⟨*v*^(*j*)^⟩_(0)_, wherein:

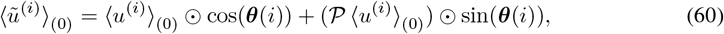

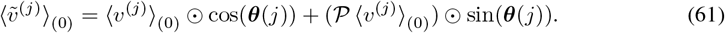

The edge disagreement from the consensus gradient update step is now 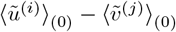 in place of ⟨*u*^(*i*)^⟩_(0)_ − ⟨*v*^(*j*)^⟩_(0)_, whereby we consider 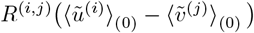.

### E Discrete diffusion

The Odyssey transformer stack learns and generates following the method of score-entropy discrete diffusion (SEDD) [44], in which tokenized data gets corrupted in a forward process and reconstructed in a reverse process. Throughout this section, we use a flattened batch where *B* = 1. We recall that the tokens of Table 3 are *special tokens* and that all other tokens are *content tokens*.

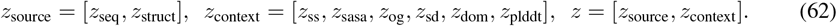

**Table 3:** Special token inventory and usage.

| ID | Token | Usage | Consensus Mechanism |
| --- | --- | --- | --- |
| 0 | PAD | Pads track to fixed maximum length | Excluded from consensus |
| 1 | MASK | Replaces content tokens during training | Included in consensus |
| 2 | UNK | Fallback for out-of-vocabulary or invalid symbols | Excluded from consensus |
| 3 | BOS | Left delimiter for variable-length tracks | Excluded from consensus |
| 4 | EOS | Right delimiter for variable length tracks | Excluded from consensus |
| 5 | IGN | Ignores empty tokens within context tracks | Excluded from consensus |

The tokens of *z* are indexed as *z*^(*i*)^. We denote the track of the *i*-th token of *z* by track(*i*), where track : {0, …, len(*z*) – 1} → {seq, struct, ss, sasa, og, sd, dom, plddt}, and len(*z*) = (5 + *K*)*L* + *G* + *H* is the total number of tokens in *z*. In the forward process, every content token of *z* evolves independently under a continuous-time Markov Chain (CTMC) which does not use the transformer stack. For the source tracks (sequence and structure), the CTMC may corrupt content data to a `MASK` token, and for context tracks (secondary structure, SASA, orthologous groups, semantic descriptions, domains, and pLDDT), the CTMC may corrupt content tokens to an `IGN` token. This transition from content to corruption tokens is the only (non-identity) transition permitted by the forward CTMCs.

The reverse process, simulated by the transformer stack, learns to undo the forward process through the simulation of its reverse, and is a collection of reverse CTMCs. The transition from `MASK` and `IGN` tokens to content tokens is the only (non-identity) transition permitted by the reverse CTMCs.

#### E.1 Forward process

Before tokens are supplied to the transformer stack, they are first evolved under the forward process, which we emphasize does not involve the transformer or its parameterization. In the forward process, tokenized data is corrupted, and some content tokens are replaced by `MASK`s (for sequence and structure tracks) or `IGN`s (for all other tracks). Special tokens are not overwritten and do not evolve under any CTMC. We begin the forward process by setting *z*_0_ = *z, t* = 0. Over time, *t* : 0 ≤ *t* ≤ *T*, each content token 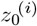 of *z*_0_ evolves under a CTMC (independently of every other token) to form 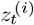. The state space of this CTMC is the union of the (track-specific) vocabulary (see Table 4) with the special tokens `MASK` and `IGN`, and hence the CTMC state space is dependent upon its position’s track. The composition of each of these independent forward CTMCs comprises the *forward process*. We first define the *noise schedule*:

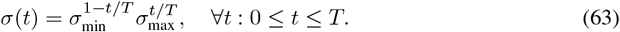

**Table 4:** Summarizing source and context tracks in Odyssey.

| Track | Raw Data | Tokenized Data | Per-Residue? | Track Type |
| --- | --- | --- | --- | --- |
| Sequence | $\{0, \dots, 24\}$ | $\{0, \dots, V_{\text{seq}} - 1 + S\}$ | Yes | Source |
| Atomic coordinates | $\mathbb{R}^{A \times 3}$ | $\{0, \dots, V_{\text{struct}} - 1 + S\}$ | Yes | Source |
| Secondary structure | $\{0, \dots, 8\}$ | $\{0, \dots, V_{\text{ss}} - 1 + S\}$ | Yes | Context |
| SASA | $\mathbb{R}^+$ | $\{0, \dots, V_{\text{sasa}} - 1 + S\}$ | Yes | Context |
| Orthologous groups | OG-tags | $\{0, \dots, V_{\text{og}} - 1 + S\}^{32}$ | No | Context |
| Semantic descriptions | Words | $\{0, \dots, V_{\text{sd}} - 1 + S\}^{512}$ | No | Context |
| Domains | Words | $\{0, \dots, V_{\text{dom}} - 1 + S\}^4$ | Yes | Context |
| pLDDT | $[0, 1]$ | $\{0, \dots, V_{\text{plddt}} - 1 + S\}$ | Yes | Context |

Per [44], for track-specific vocabulary size, *V* ^(*i*)^ = *V*_track(*i*)_ (see Table 4), we define:

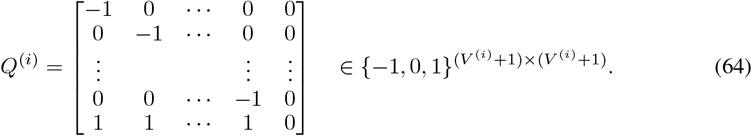

We further define the time-dependent generator matrix as:

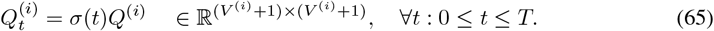

During the forward process, each content position, *i*, evolves under the CTMC defined by the time-dependent infinitesimal generator matrix, 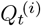. The state of the CTMC (starting from *z*_0_^(*i*)^) at time, *t*, is given by *z*_*t*_^(*i*)^, where *i* ∈ *{*0, …, len(*z*) − 1*}*. While each content position, *i*, of protein *z*_0_ evolves independently of all other content positions, all do not follow the same CTMC. We add one to each track-specific vocabulary size to enable the CTMC to transition to the `MASK` token state.

Per Eq. (64) and Eq. (65), in the forward CTMC, only transitions from a content token to a `MASK` or `IGN` token are permitted. A content token may not transition to another content token, and a content token that transitions to a `MASK` or `IGN` never transitions a second time. The `BOS, EOS, PAD` and `UNK` tokens of Table 3 do not evolve and are held constant for all time, *t* : 0 ≤ *t* ≤ *T*. The forward process is the composition of all tokens’ forward CTMCs. The time-dependent tokenized input is given by:

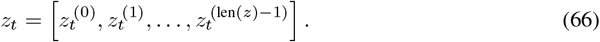

We define *p*_0_ to be the probability distribution over the tokenized input, *z*_0_, at *t* = 0, and we assume our training dataset consists of independent and identically distributed samples from *p*_0_. We further define *p*_*t*_ to be the probability distribution over all *t*-corrupted proteins, *z*_*t*_. Since each content position evolves independently, the transition kernel *p*_*t* | 0_*(z*_*t*_ | *z*_0_) can be factorized over content positions, *i*. For a protein, *z*_0_, with only content tokens:

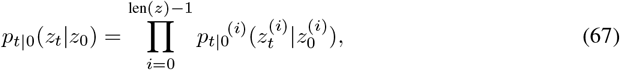

where each factor *p*_*t*|0_^*(i)*^ is a solution to:

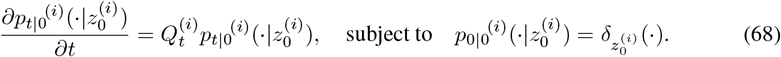

In order to analytically express *p*_*t*|0_^*(i)*^ of Eq. (67), we first introduce the *cumulative noise*:

##### Remark E.1.

*For the geometric noise schedule of Eq. (63), the cumulative noise at time, t, is:*

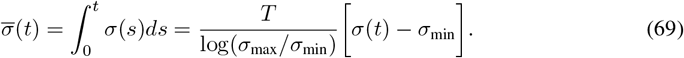

*We provide the complete derivation in Derivation J.1 of Section J*.

We utilize the cumulative noise to express the forward CTMC conditional density:

##### Remark E.2.

*The p*_*t*|0_^*(i)*^ *(which is a solution to Eq. (68)) can be expressed via the following, where* 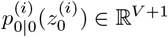 *is the one-hot vector corresponding to* 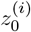:

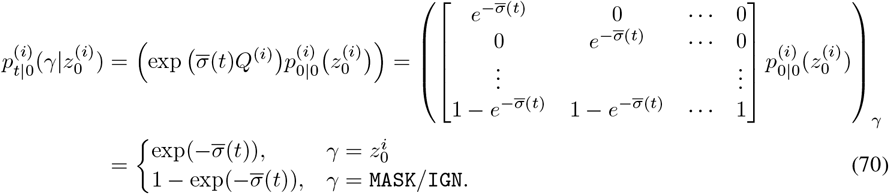

*We provide the complete derivation in Derivation J.4 of Section J*.

If we were to let *T* = ∞, then the limiting distribution starting from state 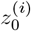 is given by 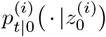 for arbitrarily large *t*, wherein it follows that:

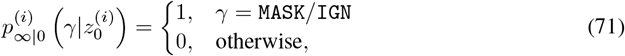

for a protein *z*_0_ with only content tokens. As this limiting distribution holds for all starting states, we know the stationary distribution is also:

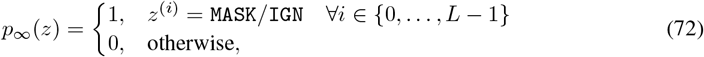

for *z*_0_ with only content tokens. The forward process applies `MASK` and `IGN` tokens to content tokens over len(*z*) independent CTMCs (excluding the positions of special tokens). The transformer does not affect the model-free forward process, but subsequently aims to replace `MASK`s and `IGN`s with content tokens in the *reverse process*.

#### E.2 Reverse process

During the reverse process, the transformer replaces `MASK`s in the sequence and structure tracks with content tokens — the opposite of the forward process’ substitution. Consequently, the transformer endeavors to generate a reverse path, (*ω*_source,*T* −*t*_)_0≤*t*≤*T*_, that approximates as closely as possible (*z*_source,*t*_)_0≤*t*≤*T*_. For our exposition of the forward process, the true reverse process is given by:

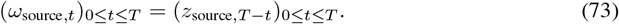

Denoting *p*_*t*_ as the probability distribution over *z*_*t*_, the probability distribution over *ω*_source,*t*_ is:

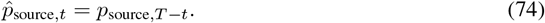

The same convention applies in defining 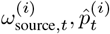. In the forward process, only content positions are free to transition states, and may transition only to `MASK` and `IGN` (states that can solely absorb). Conversely, in the reverse process, only `MASK` and `IGN` may transition (as these states are exit-only) and the only permitted destination states are content tokens. In both the forward and reverse processes, `BOS`, `EOS`, `PAD`, and `UNK` tokens remain unaffected. We recall that forward process is formulated as independent CTMCs for each token position, and that the generator 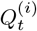 does not depend on the state at other token positions. However, in the reverse process, the infinitesimal generator 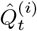 depends on the whole input *z*_*t*_ and is related to *Q*^*(i)*^ by the following expression:

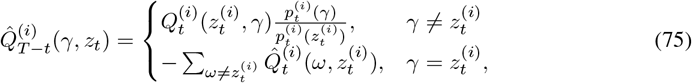

where we define 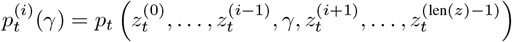.

Thus, an estimate of 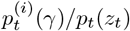 for *γ*∈ { 0, …, *V* ^(*i)*^} is sufficient to simulate the reverse-time CTMC and thus the reverse process. We define the true per-residue probability ratios, 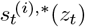 as:

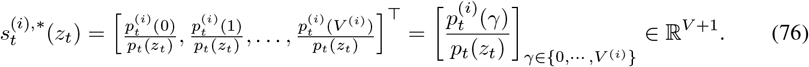

The Odyssey transformer stack is trained to represent a function, 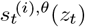, *parameterized* by *θ* that approximates 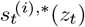. We now express 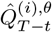 using this approximation:

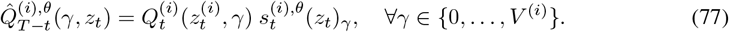

As we outline in Section F.3, the Odyssey transformer stack outputs two distinct sets of estimated probability ratios, which correspond to the sequence and structure tracks:

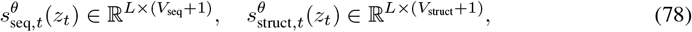

where 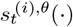 is equivalent to the *i*-th row of 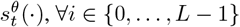. We recall the CTMCs of the forward process evolve independently of one another. However, in the reverse process, though each initially `MASK`ed position maintains its own time-dependent state, and though the transformer’s estimated probability ratios, *s*_*t*_^(*i*),*θ*^(*z*_*t*_), are sampled independently across the various token positions, it is the case that any given transformer output ratio is conditioned upon the *entire transformer input z*_*t*_, *across all tracks*. Therefore the transition in the reverse CTMC at position *i* at time *t* : 0 ≤ *t* ≤ *T can influence the* transition in position *j*’s CTMC at time *t*^*′*^ : *t*^*′*^ *> t*, ∀*i, j* ∈ {0, …, *L* − 1}.

### F Architecture

The Odyssey architecture comprises two primary components — a *finite scalar quantizer* (FSQ) for tokenizing protein structure, and a *transformer stack* for multimodal representation learning. The FSQ quantizes continuous atomic coordinates into discrete structure tokens, which can be embedded and summed with the embedded sequence tokens (with the embedded context tracks introduced through cross-consensus and cross-attention). The transformer stack operates on this joint representation. We outline the FSQ in Section F.2, and the transformer stack in Section F.3. For all the experimental results of this paper, the model configurations and training hyperparameters are given in Section K.3.

#### F.1 Blocks

Odyssey leverages recurrent blocks for localized mixing over structure coordinates and for processing embedded tracks. These include *convolutional blocks*, which comprise the FSQ, and *self-consensus blocks*, which are used by the FSQ and the transformer stack. We delineate the components comprising the convolutional and self-consensus blocks below (see Figures 10, 11 and 12 for schematics).

**Figure 10:**
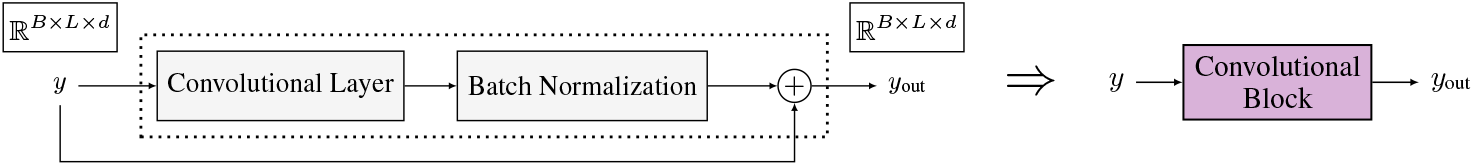
Convolutional block schematic.

**Figure 11:**
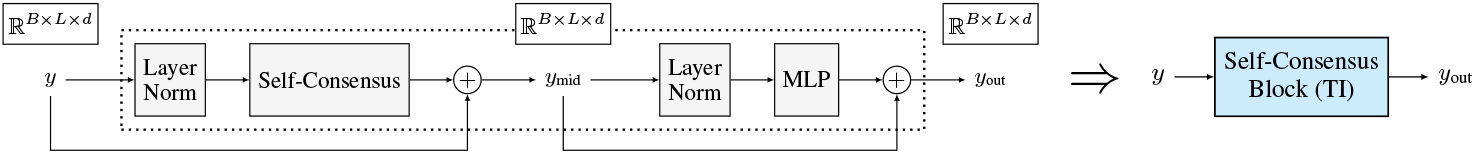
Time-independent (TI) self-consensus block schematic.

**Figure 12:**
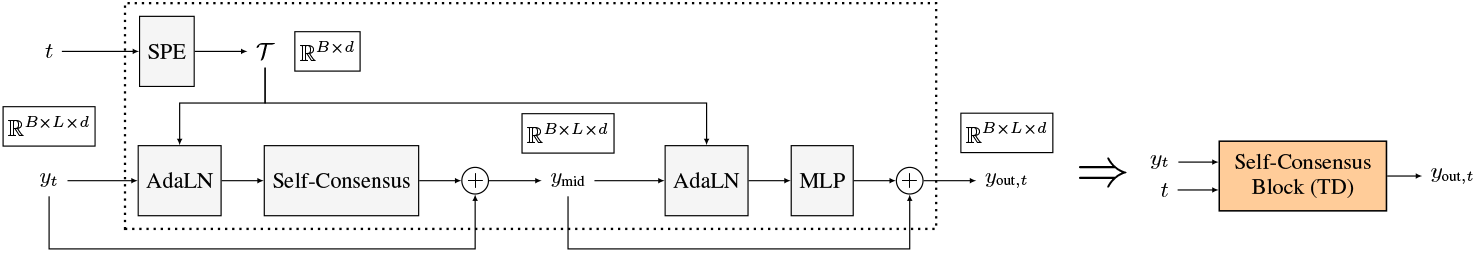
Time-dependent (TD) self-consensus block schematic.

The convolutional block has convolutional and batch normalization layers with a residual connection. We first transpose the input from ℝ^*B×L×d*^ to ℝ^*B×d×L*^ to match the expected channel-first convention. The convolutional layer applies a learnable bank of *d* × *d* filters along the length dimension to aggregate local neighborhoods and linearly mix channels at every position. Next, the batch normalization layer normalizes each channel (across the flattened “batch-and-length” axes) to have approximately zero mean and unit variance during training, and applies a learned per-channel scale and shift.

We employ two variants of the self-consensus block that differ in their normalization and conditioning. In the time-independent self-consensus block, we consider a pair of sublayers: (i) layer normalization followed by a multi-headed self-consensus mechanism with rotary positional embeddings (detailed in Section D.1) using a residual connection, and (ii) layer normalization followed by a MLP with a residual connection. In the time-dependent self-consensus block, the layer normalization is replaced with adaptive layer normalization (adaLN-zero) across both sublayers, which modulates normalized activations through a time embedding (we implement the approaches introduced by [68] and [54]) — this time embedding is obtained from sinusoidal positional embeddings (SPE) applied to a discrete time step index, *t* ∈ {0, …, *T/*Δ*t* − 1}^*B*^, to produce 𝒯 ∈ ℝ^*B×d*^, which conditions adaLN in each sublayer. We delineate this time conditioning procedure in Section F.3.1. As a precursor, we begin by outlining the FSQ architecture as follows in Section F.2.

#### F.2 Finite scalar quantizer

To discretize continuous atomic structure coordinates into structure tokens, we utilize the finite scalar quantizer (FSQ) [48] to avoid codebook collapse and the auxiliary machinery characteristic of the vector-quantized variational autoencoder (VQ-VAE) [73]. The FSQ partitions the coordinate space into a fixed lattice, where each coordinate is mapped to an index in a structured codebook, ℋ. This construction yields a vocabulary, 𝒱_struct_, which is comparable to the sequence alphabet, 𝒱_seq_. As it pertains to Odyssey, we summarize this construction below:

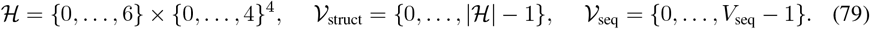

We now outline the FSQ encoder in Section F.2.1 and the FSQ decoder in Section F.2.2.

##### F.2.1 Encoder

The FSQ encoder produces discrete structure tokens, *z*_struct_ ∈ {0, …, |ℋ | − 1}^*B×L*^, from 3-backbone continuous atomic structure coordinate inputs, *x*_struct_ ∈ ℝ^*B×L*×3×3^, via the following steps:

1. **Projector**: ℝ^*B×L*×3×3^ → ℝ^*B×L×*|ℋ |^. The structure coordinates, *x*_struct_, are flattened and projected to a *d*-dimensional space, passing through two convolutional blocks and 16 time-independent self-consensus blocks, before being projected to the -dimensional vector, *x*_proj_ ∈ ℝ^*B×L×*| ℋ|^. We schematize this projector in Figure 13.
2. **Quantizer**: ℝ^*B×L×*| ℋ|^ → ℋ^*B×L*^. The FSQ maps each |ℋ|*-*dimensional vector, *x*_proj_, to a tuple, *x*_quant_ ∈ ℋ^*B×L*^ using an axis-aligned lattice. We detail the quantizer in Section F.2.4.
3. **Indicizer**: ℋ ^*B×L*^ → { 0, …, |ℋ |− 1} ^*B×L*^. Each tuple, *x*_quant_, is mapped to an integer token via lexicographic indexing to form the discrete structure tokens *z*_struct_ ∈ {0, …, ℋ − 1 ^*B×L*^}. We further delineate the indicizer in Section F.2.3.

**Figure 13:**
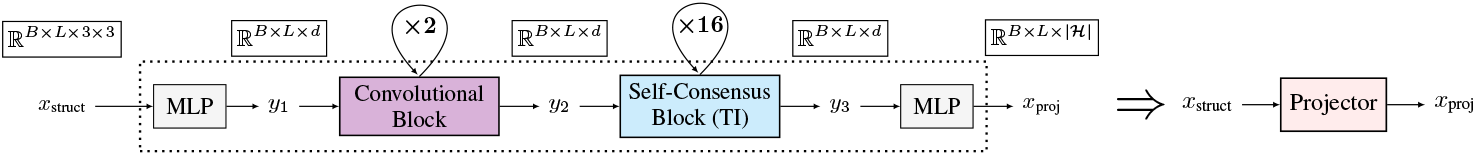
FSQ projector schematic. The projector has convolutional and self-consensus blocks.

We illustrate the FSQ encoder in Figure 14, alongside its compressed form (Enc), which will be used in future schematics. The encoder output, *z*_struct_, is embedded and positionally summed with the tokenized and embedded sequence, *y*_seq_, in the transformer stack (see Section F.3).

**Figure 14:**
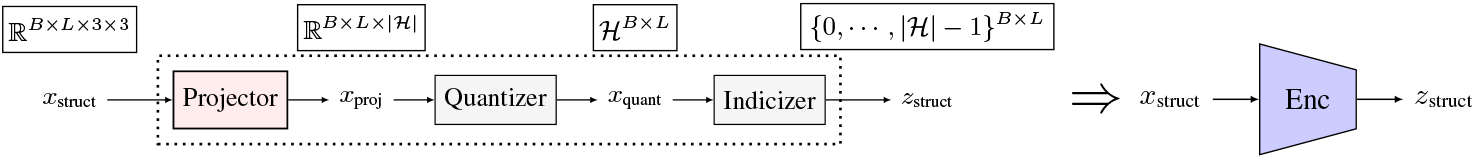
FSQ encoder schematic. The encoder produces discrete-valued structure tokens.

##### F.2.2 Decoder

The FSQ decoder produces reconstructed structure coordinates, 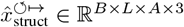, from discrete structure token inputs, *z*_struct_. Per Section G.1, we employ the FSQ encoder from Section F.2.1 with decoders trained in two stages. In stage-1, we train the encoder jointly with a stage-1 decoder (Dec_(1)_) to reconstruct the 3-backbone representation, where *A* = 3. In stage-2, we freeze the trained stage-1 encoder and, using the embedded sequence tokens, *y*_seq_, as input, train a stage-2 decoder (Dec_(2)_) to reconstruct the atom-14 representation, where *A* = 14. The stage-1 and 2 FSQ decoders consists of:

1. **Codifier**: {0, …, |ℋ| − 1} ^*B×L*^ → ℋ^*B×L*^. Each integer structure token, *z*_struct_, is mapped through inverse lexicographic indexing to recover the quantized tuple, *x*_cod_ ∈ ℋ ^*B×L*^. We further outline the codifier in Section F.2.3.
2. **Dequantizer**: ℋ ^*B×L*^ → ℝ^*B×L×*| ℋ|^. *The FSQ* maps each tuple, *x*_cod_, to an |ℋ |*-*dimensional vector, *x*_deq_ ∈ ℝ^*B×L×*| ℋ|^, using the same lattice. We detail the dequantizer in Section F.2.4.
3. **Expander**: ℝ^*B×L×*| ℋ|^ → ℝ^*B×L×A×*3^. The |ℋ| -dimensional vector, *x*_deq_, is projected onto a *d*-dimensional space, passing through 16 time-independent self-consensus blocks in the stage-1 decoder (and 48 in the stage-2 decoder), followed by two convolutional blocks and a map to the reconstructed structure coordinates, 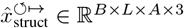. Figure 15 depicts the stage-1 expander (from Dec_(1)_), and Figure 16 depicts the stage-2 expander (from Dec_(2)_).

We illustrate the stage-1 FSQ decoder in Figure 17, alongside its compressed form (Dec_(1)_), and the stage-2 FSQ decoder in Figure 18, alongside its compressed form (Dec_(2)_).

**Figure 15:**
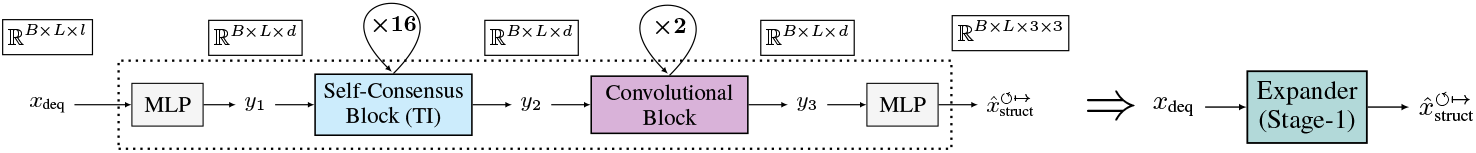
FSQ stage-1 expander schematic. The expander comprises the stage-1 decoder (Dec_(1)_).

**Figure 16:**
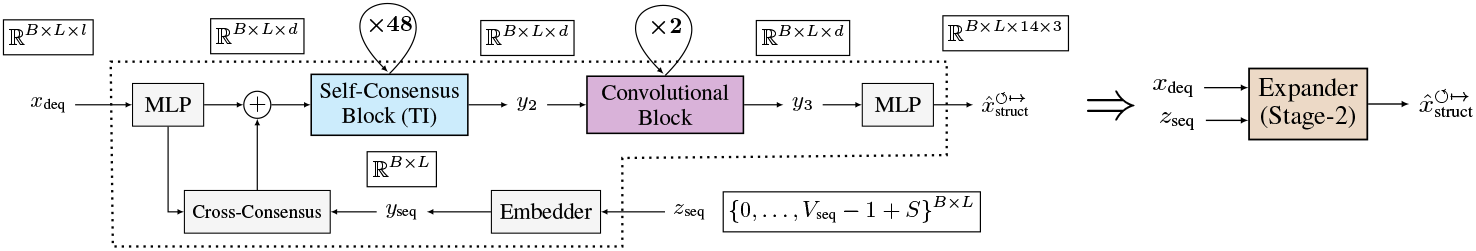
FSQ stage-2 expander schematic. The expander comprises the stage-2 decoder (Dec_(2)_).

**Figure 17:**
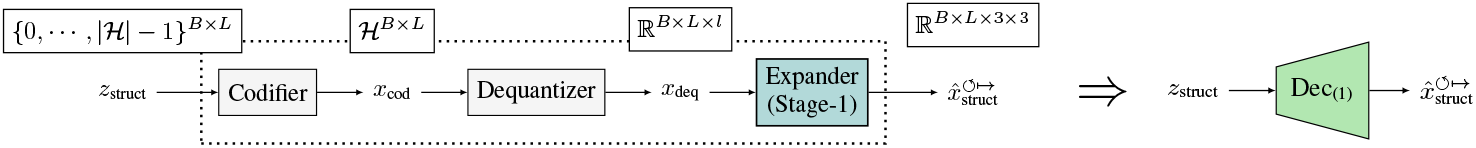
FSQ stage-1 decoder schematic. The decoder reconstructs the 3-backbone representation.

**Figure 18:**
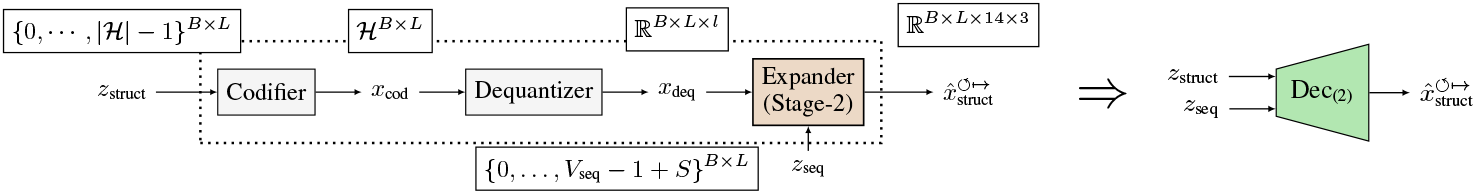
FSQ stage-2 decoder schematic. The decoder reconstructs the atom-14 representation.

##### F.2.3 Indicizer and codifier

For a codebook of size (𝔥_0_, …, 𝔥_| ℋ|−1_) (which is (7, 5, 5, 5, 5) for the codebook of Eq. (79)), a code, 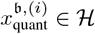, where 𝔟 ∈ {0, …, *B* − 1} and *i* ∈ {0, …, *L* − 1}, is given by:

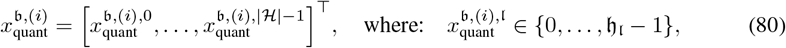

∀𝔩 ∈ {0, …, |ℋ |−1}. The indicizer yields indices, *z*_struct_ ∈ {0, …, |ℋ|− 1}^*B×L*^, from codes, *x*_quant_, by Eq. (81), and the codifier yields codes from indices by Eq. (82), where 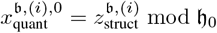.

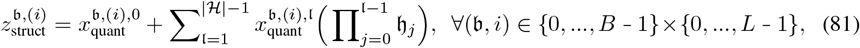

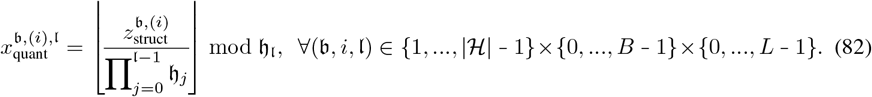

##### F.2.4 Quantizer and dequantizer

In the quantizer, each vector, *x*_proj_ ∈ *H* ℝ^*B×L×*|ℋ |^, is mapped axis-wise onto the finite lattice, ℋ, by an affine transform, clipped, and rounded [48]. For per-axis offset *m*_*l*_ *> 0, and scale* 𝔫 _𝔩_:

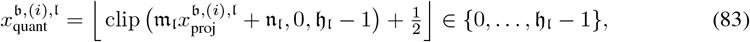

∀(𝔟, *i, l)* ∈ {0, …, *B* − 1}*×*{0, …, *L* − 1}*×*{0, …, |ℋ| − 1}. For backpropagation, the quantizer leverages the straight-through estimator (STE) [7]. In the dequantizer, given integer tuples, *x*_cod_ ∈ ℋ^*B×L*^, the real-valued vectors are recovered axis-wise by inverting the affine map:

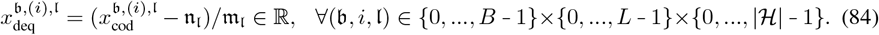

#### F.3 Transformer stack

The Odyssey transformer stack is a multimodal architecture for joint sequence and structure generation, trained using score-entropy discrete diffusion (SEDD). Odyssey treats the tokenized amino-acid sequence and FSQ-encoded structure as synchronized views of the same protein, and learns to keep them mutually consistent at every residue, *i* ∈ {0, …, *L* − 1}. The source embedding comprises the positionally summed sequence and structure embeddings, which we enrich with biological context so the model can reason about residue-level structural constraints together with global functional insight. During generation, the transformer stack outputs probability ratios at `MASK`ed positions, for which the sequence and structure tokens are filled over *T/*Δ*t* steps. After the sequence and structure are generated, the structure tokens pass through the stage-2 FSQ to obtain the atom-14 representation.

For *t* ∈ {0, …, *T/*Δ*t* − 1}^*B*^, the tokenized tracks and their embeddings are summarized as follows. We note that embedded sequence *y*_seq,*t*_ and structure *y*_*struct,t*_ are summed to form *y*_source,*t*_ *= y*_*seq,t*_ *+ y*_*struct,t*_. We note *y*_dom,*t*_ is formed by first average pooling 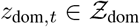 across *K*.

Context is injected into *y*_source,*t*_ in the following order, where each step is applied only for batches with at least one valid context position and added residually: (1) secondary structure, (2) SASA, (3) orthologous groups, (4) semantic descriptions, (5) domains, and (6) pLDDT. Specifically, secondary structure, SASA, domains, and pLDDT are injected by cross-consensus (since they are positionally aligned with the sequence and structure), whereas orthologous groups and semantic descriptions are imbued by cross-attention. We denote *y*_*t*_ ∈ ℝ^*B×L×d*^ as the output after all six residual updates, and summarize this injection step and its compressed representation below in Figure 19.

**Figure 19:**
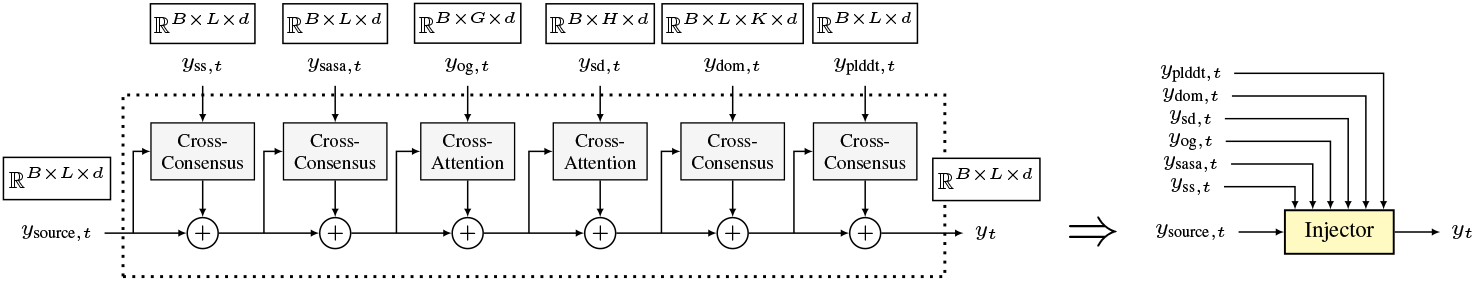
Injector schematic. The injector imbues embedded context into the embedded source.

The embedding output, *y*_*t*_, then passes through *N* repeated time-dependent self-consensus transformer blocks, where normalization is modulated by the discrete time step, *t* ∈ {0, …, *T/*Δ*t* − 1}^*B*^. We outline this time conditioning step in Section F.3.1. A final adaptive layer-normalization and linear projection outputs per-residue probability ratios, 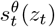, for the sequence vocabulary and structure vocabulary. We illustrate the transformer stack processing pipeline below in Figure 20.

**Figure 20:**
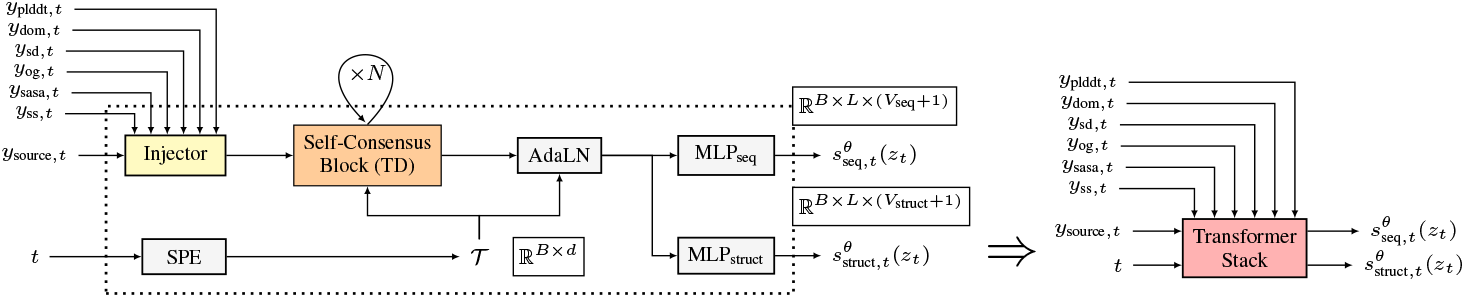
Transformer stack schematic. The transformer stack predicts per-residue probability ratios.

##### F.3.1 Time conditioning

Let *t* ∈ {0, …, *T/*Δ*t* − 1}^*B*^ be the discrete time step and *y*_*t*_ ∈ ℝ^*B×L×d*^ as the embedding output. Following the guidelines of [30], we form the sinusoidal positional embedding of *t*, 𝒯 ∈ ℝ^*B×d*^, and pass it through an MLP to obtain per-channel scale and shift, 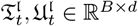 for 𝔩 ∈ {0, …, *d* − 1}. Further adapting [68], we first layer-normalize *y*_*t*_, and modulate it via adaptive layer normalization with zero-initialized modulation (adaLN-zero) [54]:

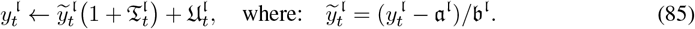

Above, 𝔞, 𝔟 ∈ ℝ^*B×L*^ are the mean and standard deviation, broadcast along 𝔩. In adaLN-zero, the map from 𝒯 to 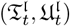 is initialized to zeros so the network begins with the identity mapping.

### G Training

In this section, we detail the methods of training the FSQ and transformer stack comprising Odyssey. More information regarding the training of models in Section F are provided in Section K. For clarity of exposition, we restrict all formulas of this section to the case where the batch size, *B* = 1.

#### G.1 Finite scalar quantizer training

We first outline how the Odyssey FSQ is trained to discretize atomic structure coordinates. The FSQ is an autoencoder architecture, and its training leverages a reconstruction loss. Unlike the output of a vanilla autoencoder, the output of our FSQ is a geometric object, and it is possible that the FSQ can induce translations or rotations in its reconstruction without inducing distortion or reflection. Thus, to make our loss invariant to translations or rotations, we first translate and rotate the reconstructed coordinates to best align them to the input atomic structure coordinates before we calculate losses.

We split the training into two stages — in stage one (stage-1) training, an encoder and decoder are trained with masked coordinates, with masks sampled according to the discrete diffusion corruption schedule from Section E. In stage two (stage-2) training, a new decoder is paired with the encoder of stage one training (now frozen) with unmasked coordinates.

Throughout this section, we shall refer to atomic coordinates as *x*_struct_ ∈ ℝ^*L*×14×3^. When we wish to refer only to the first *A* atomic coordinates per residue, we shall denote as [*x*_struct_]^:,*:A*,:^ ∈ ℝ^*L×A*×3^ *(i.e*., 3-backbone(*x*_struct_) = [*x*_struct_]^:,:3,:^ ∈ ℝ^*L*×3×3^).

For some portions of this section, masks are applied to the input atomic coordinates. A single time step *t ∼* 𝒰 ([0, *T*]) is chosen uniformly, for which the corresponding mask is denoted as 𝕄_*t*_ ∈ { 0, 1} ^*L*^, with zeros comprising masked positions and ones in all other positions. For some equations, we mask *x*_struct_ via *x*_struct_ ⊙ 𝕄_*t*_ ∈ ℝ^*L×A×*3^, where the multiplication is element-wise and broadcasts over all *L* residues, and all 3 dimensions of Euclidean space. For other expressions, we preserve only the Σ 𝕄_*t*_ residues of *x*_*struct*_ corresponding to unmasked coordinates, and denote this by 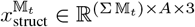. We emphasize that *x*_struct_ ⊙ 𝕄_*t*_ and 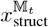 are not equal in dimension.

##### G.1.1 SE(3)-invariant loss

Among vanilla autoencoders [5], which we delineate as 𝒜, the mean-squared error (MSE) loss is the de-facto loss metric of choice:

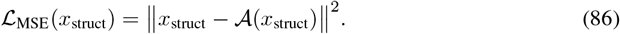

Though the FSQ is a particular type of autoencoder, it is nonetheless one that can induce translations and rotations in its output. For FSQ input *x*_struct_, we denote the output 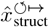, with the superscripts emphasizing the possible translational and rotational artifacts remaining after FSQ reconstruction. The reconstructed 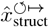 could be an exact clone of *x*_struct_ after a translation and rotation, but under ℒ_MSE_, this exact replica protein structure would yield a significant and positive loss.

Thus, we seek a loss metric that is invariant to translations and rotations (*SE*(3)-invariance). To do so, we translate and rotate the reconstructed coordinates to offset any translation introduced by the FSQ. We begin by defining the 3-centroid of the protein:

###### Definition G.1

(3-centroid). *We define the* 3*-centroid of a protein of length N to be the geometric centroid of the first* 3*-backbone atoms (see Definition C.1):*

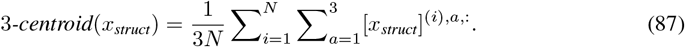

We now introduce a modified version of the Kabsch algorithm [34] in Algorithm 1. Since *x*_struct_ is translated so that its 3-centroid is aligned with the origin prior to being fed into the model, the Kabsch algorithm similarly translates 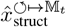 to form 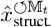 with the origin-aligned 3-centroid. Next, the algorithm finds the matrix, *R* _KABSCH_, which rotates 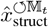 to best align with 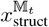 in the first *A*^↺^ atomic positions. For both the translation and rotation steps, only unmasked residues are considered.

###### Algorithm 1 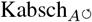 Algorithm

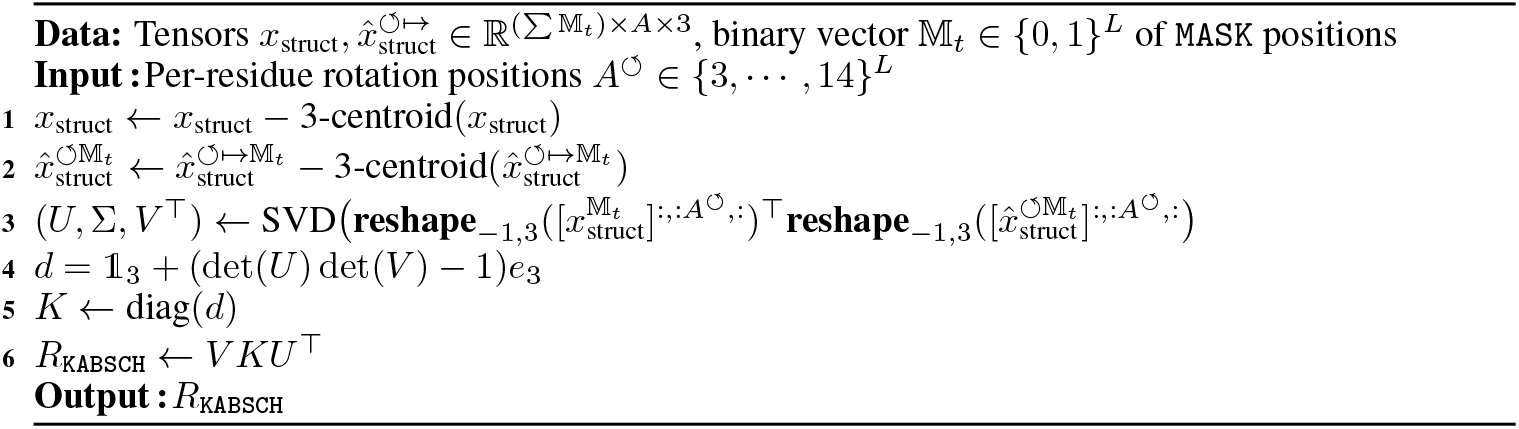

Following [34], we next offer a proof of the optimality of the matrix of Algorithm 1:

###### Theorem G.2

(Kabsch Algorithm). *For* 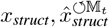 *with origin 3-centroids, and for a binary vector indicating non-*`MASK` *positions* 𝕄_*t*_ ∈ { 0, 1} ^*L*^, *let R*_KABSCH_ *denote the output of Algorithm 1. Suppose SO*(3) *denotes the set of rotation matrices in* ℝ^3*×*3^. *Then:*

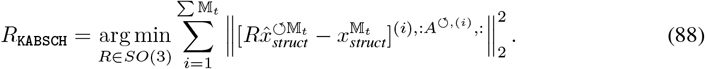

*We provide the complete proof in Proof J.5 of Section J*.

##### G.1.2 Stage one training

The focus of stage one is the training of the FSQ encoder from Section F.2.1. While the FSQ does not perform generative infilling, we nonetheless expose the FSQ encoder to masked atomic coordinates during training. This ensures that masked coordinates encountered at inference time fall within the FSQ encoder’s training distribution. Without this exposure, heavily masked inputs at inference time would produce out-of-distribution structure tokens for the transformer, degrading generation.

Suppose ℳ is a probability distribution supported on {0, 1} ^*L*^. For any protein, we sample 𝕄_*t*_^†^*∼* ℳ. *The sampled* 𝕄_*t*_^†^ will contain zeros in desired `MASK` positions and ones in all other positions. Although we assume *B* = 1 in this section, we note that the sampling of 𝕄_*t*_ is not necessarily independent and identically distributed (i.i.d.) across distinct proteins in or across training batches but *is* i.i.d. across residues within a given protein. As the other fully-tokenized per-residue tracks will contain `BOS`, `EOS`, `PAD` *and* `UNK` *tokens, we modify* 𝕄_*t*_^†^ so that `MASK`s will not be placed in these positions. For a given protein *x*_struct_, let S(*x*_struct_) 0, ∈ {0, 1} ^*L*^ be a vector containing zeros in the positions of `BOS`, `EOS`, `PAD`, and `UNK` positions and ones in all other positions. Then:

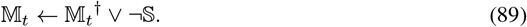

We then pass *x*_struct_ ⊙𝕄 _*t*_ through the encoder to obtain 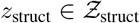, then pass *z*_struct_ ⊙ 𝕄_*t*_ through the decoder to obtain 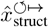, and we apply Algorithm 1 to produce 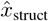. We illustrate the stage one training procedure in Figure 21 below:

**Figure 21:**
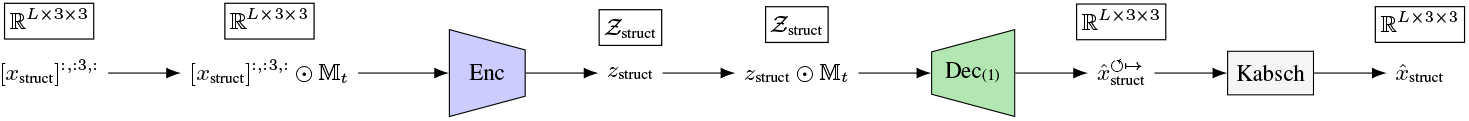
Stage one FSQ training.

The encoder and decoder are simultaneously trained to minimize the objective ℒ_stage-1_ on expectation over our distribution of proteins, where:

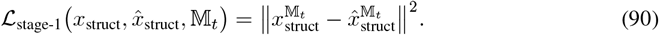

We outline the optimization procedure in Algorithm 2, where **3**_*L*_ is an *L-*dimensional vector of threes:

###### Algorithm 2 FSQ Stage One Training Algorithm

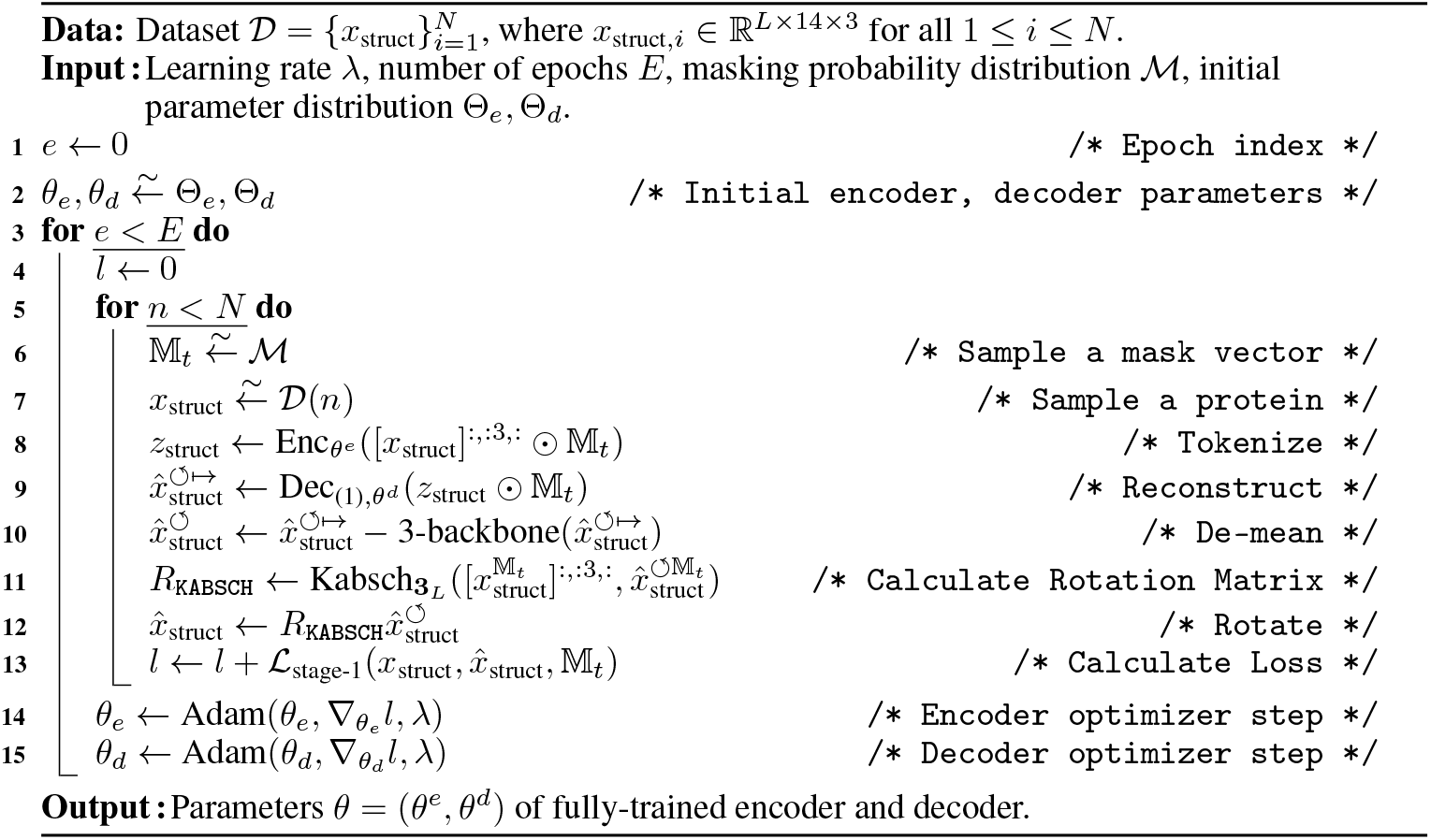

##### G.1.3 Stage two training

The fully-trained encoder from stage one training is then ported and paired with a new decoder. In stage two training, the parameters of the encoder are frozen, and only the FSQ decoder parameters are updated. As the FSQ decoder will not encounter `MASK` tokens at inference time (since only completely infilled structure tokens are fed through the stage 2 decoder), we do not expose the stage 2 FSQ to `MASK`ed atomic structure coordinates during training.

The FSQ decoder outputs an atom-14 (see Section B) representation of the residues, however, most residues do not contain 14 atoms, and the ordering of the identities of the atoms within the atom-14 representation varies between the different amino acid types. Thus, it is necessary for the decoder to receive as input the sequence of the protein so that it knows the ordering and identities of the various side-chain atoms. The total number of atoms in a given protein is given by:

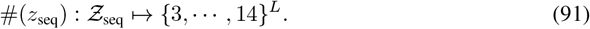

We schematize the stage two training procedure in Figure 22 below:

**Figure 22:**
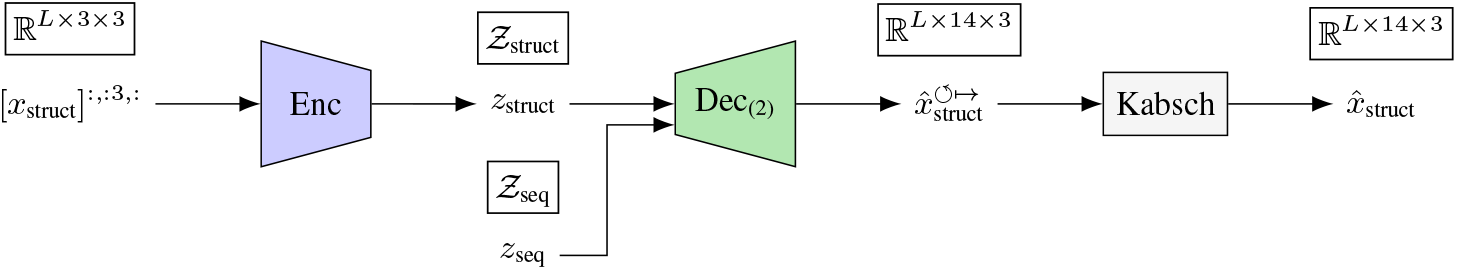
Stage two FSQ training.

During stage two training, the encoder is frozen. The decoder is trained to minimize the objective ℒ _stage-2_ on expectation over our distribution of proteins, where:

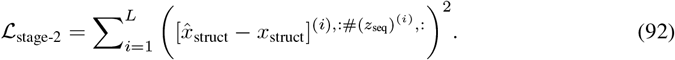

The complete stage two training procedure is summarized in Algorithm 3.

###### Algorithm 3 FSQ Stage Two Training Algorithm

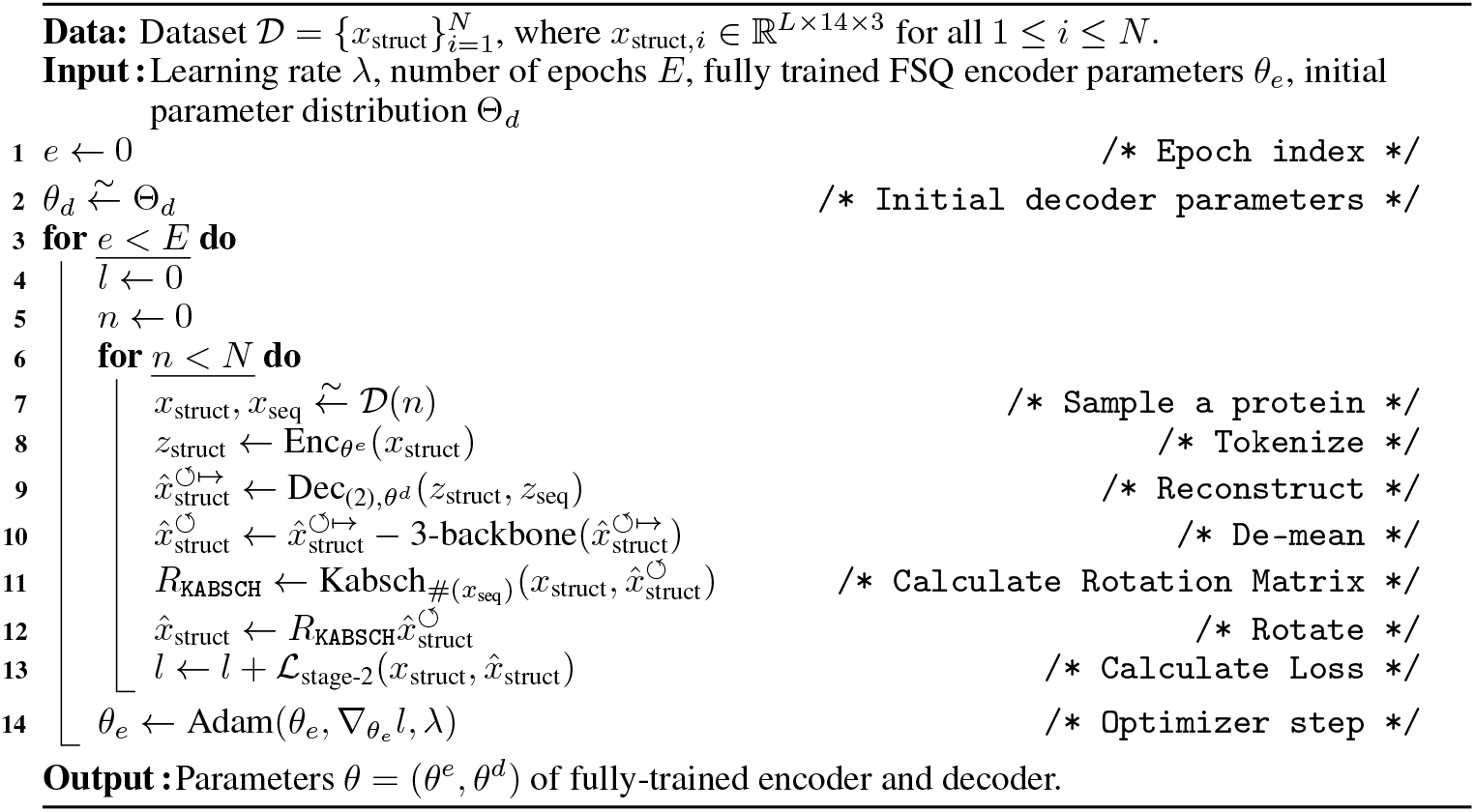

#### G.2 Transformer stack training

Fully-tokenized tracks undergo the forward process (see Section E.1) before being fed to the transformer stack. During the forward process, some tokens will be overwritten with `MASK`*/*`IGN` tokens. For each protein supplied to the model, a single time step *t* ∼ 𝒰 ([0, *T*]) is chosen uniformly (although this *t* may vary between proteins in a common batch). Various proteins in a batch may have different values of *t*, but all tracks across a single protein share a common *t*. On expectation, the proteins with higher values of *t* will have a greater proportion of tokens overwritten with `MASK`s (see Remark E.2). The special tokens of Table 3 are preserved and never overwritten by `MASK`*/*`IGN` tokens.

Following Eq. (62), the tokenized sequence post-forward-process is denoted by *z*_*t*_. We define 𝕄_*t*_ to be the binary vector with zeros in the positions of `MASK`*/*`IGN` *tokens* and ones in all others:

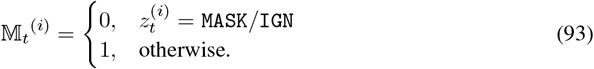

Following [44], we introduce the diffusion-weighted denoising score entropy, which serves as our training objective for the transformer stack:

##### Definition G.3

(Diffusion-Weighted Denoising Score Entropy (DWDSE)). *For track* ∈ {*seq, struct*}:

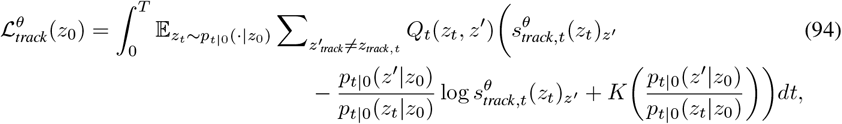

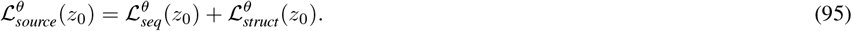

We now present a theorem following Theorem 3.6 from [44] which relates the perplexity of the learned 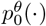 to the objective 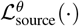:□

##### Theorem G.4

(Score-Entropy and Perplexity Bound). *For* 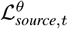 *given in Definition G.3*,

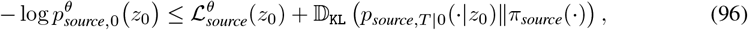

*where* 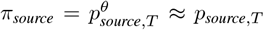 *is the initial distribution during generation. We provide the complete proof in Proof J.6 of Section J*.

Accordingly, the upper-bound on the negative log-likelihood of the ground-truth token can be denoted as the sum of the loss term, 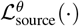, and a term independent of *θ*. By optimizing *θ* to minimize the average 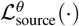, we minimize the negative log-likelihood (and the perplexity of our model). While 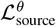 comprises an expectation over the forward process and is inconvenient to calculate, we use a per-track estimator, 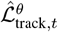, instead. We note that 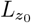 denotes the length of each uncorrupted protein.

##### Definition G.5

(DWDSE Estimator). *For track* ∈ {*seq, struct*}, *we define:*

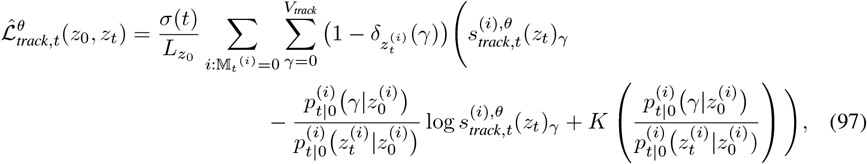

*where K*(*a*) = *a*(log(*a*) − *1) and δ is the Kronecker delta function*.

##### Remark G.6.

*If t* ∼ 𝒰 *([0, T]), and z*_*t*_ *drawn following Remark E.2, then the estimator*, 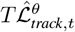 *of Definition G.5, is an estimator of the* 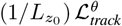.

*We provide the complete proof in Proof J.7 of Section J*.

The estimator of Definition G.5 is a calculable metric for each track ∈ {seq, struct}. We weight and combine the two source tracks to arrive at the final objective for transformer track training:

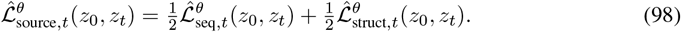

##### Algorithm 4 Transformer Stack Training Algorithm

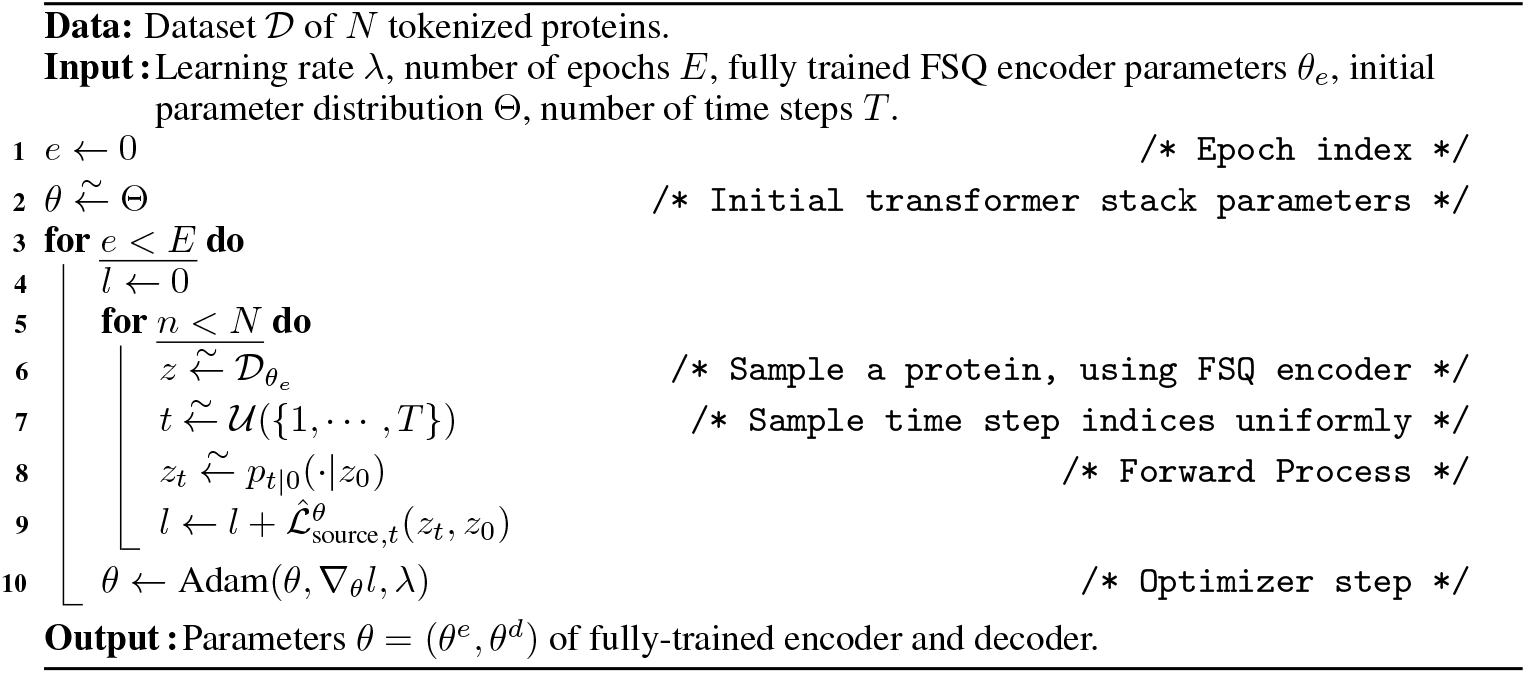

### H Generation

The generation of novel proteins with Odyssey begins with an *original protein* — a known, existing protein upon which Odyssey can improve. Across all tracks, the designer chooses which residues to keep and which to replace with a special token so that Odyssey can infill. For the sequence and structure tracks, this special token is the `MASK`, and for all other tracks it is the `IGN`. *This designer’s* `MASK`*/*`IGN`-corrupted protein is denoted by *z*_*T*_. While the designer plays the role of a forward process, it is the transformer stack that will simulate the reverse process and generate *z*_source,*t*_.

Let 𝕄 be a binary tensor containing ones in the positions of *z*_0_ that the designer intends to preserve and zeros in all other positions. Then, *z*_*T*_ can be written as:

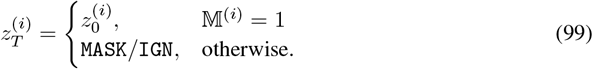

Across all positions, *i*, such that 𝕄^(*i*)^ = 0, the process of replacing 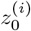 *by* `MASK` can be viewed as sampling from 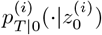 (see Eq. (71)) for an arbitrarily large value of *T*, where the distribution, 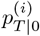, converges to the limiting distribution (see Eq. (71)), equivalent to its stationary distribution (see Eq. (72)), which places all probability at the `MASK`/`IGN` token.

For sufficiently large *T*, 𝔻_KL_(*p*_∞_(·) *p*_*T* |0_(· | *z*_0_)) is small, so replacing content tokens by `MASK`s/`IGN`s is approximately equivalent to sampling *p*_*T* | 0_(| *z*_0_) via the forward process. Thus, the action of the designer plays the role of the forward process for reasonable choices of *T*, and the starting point of the reverse CTMC can be viewed as a random draw from the forward CTMC’s stationary distribution. From this starting point, we simulate the reverse process. While the true reverse process is continuous in time, we choose to simulate by dividing the time range [0, *T*] into discrete steps. We follow the time-dependent update rule provided below in Remark H:

#### Remark H

(Reverse Timestep using Tweedie’s Method and Tau Leaping). *The following discrete-time update rule numerically approximates the continuous-time reverse process:*

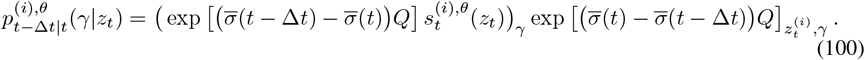

*We provide the complete derivation in Derivation J.8 of Section J*.

We denote 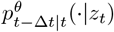 as the probability transition kernel for which 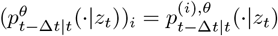 for 0 ≤ *i* < len(z). We emphasize that although the conditional probability of Remark H serves as an update rule only for the token at position *i*, it is conditioned upon the entire protein, *z*_*t*_, as the entire protein is specified as input for 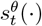. Using this update rule, we simulate the reverse CTMC across all positions in Algorithm 5, which mirrors Algorithm 3 from [44]:

#### Algorithm 5 Protein Generation

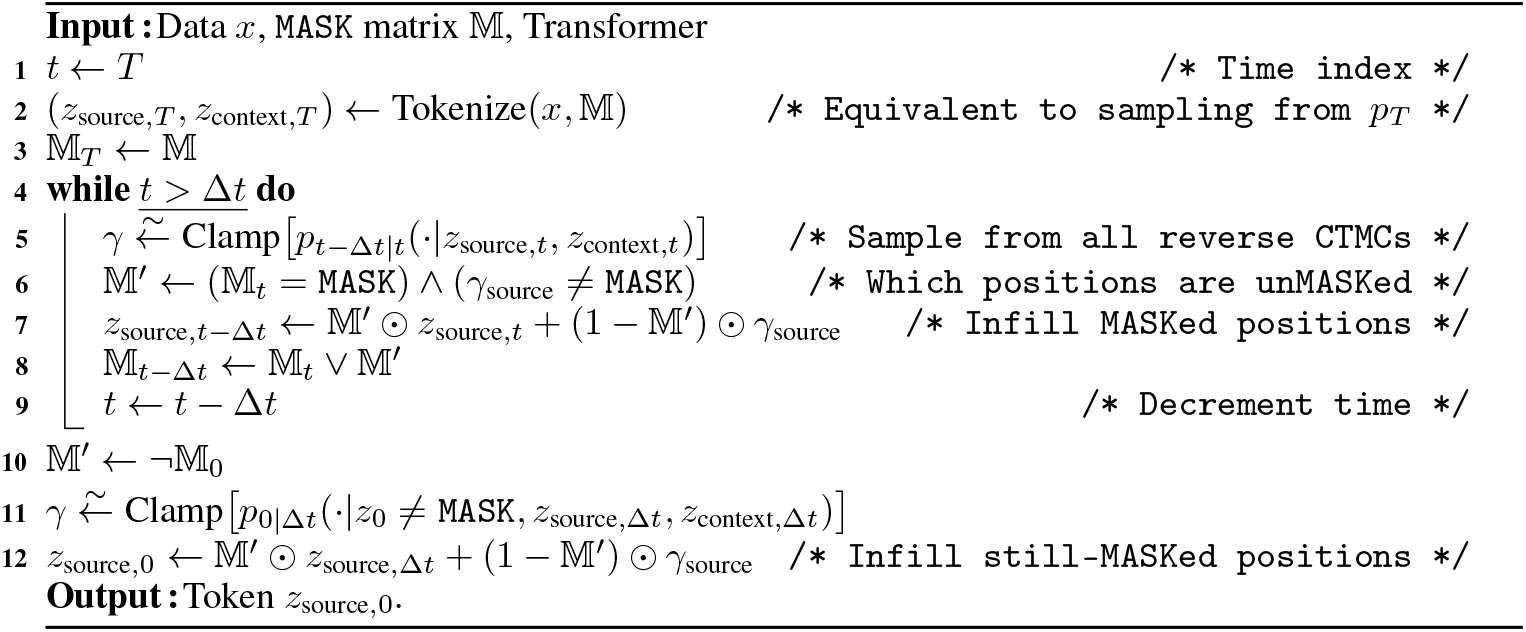

As noted by [44], the numerical estimation of *p*_*t* − Δ*t*|*t*_*(·|z*_*t*_) can produce negative values; in this event we subtract from this probability vector its most negative element in order to make sampling well-defined. Furthermore, if any `MASK` tokens remain in the protein *z*_*t*_ when *t =* Δ*t*, we sample from the conditional distribution 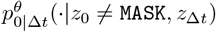 in the update step of Algorithm 5.

After *z*_0_ has been fully sampled by the transformer stack, the tokens corresponding to structure are passed through the stage two FSQ decoder to recover new structure coordinates, 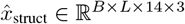. The tokens corresponding to sequence are mapped back to their amino acid counterparts.

### I Alignment

Recent works exploring the alignment of protein transformers have targeted masked language [28] and autoregressive models [10] through direct preference optimization (DPO) [58] and its extension, iterative reasoning preference optimization (IRPO) [53]. While DPO has been validated in the setting of continuous diffusion [75] (for steering denoisers using pairwise preferences), DPO for discrete diffusion (D2-DPO) is a recent formalization [12]. We adapt D2-DPO for our setting as follows.

#### I.1 Alignment Training

We align only the sequence reverse kernel while treating all other tracks as conditioning. Let *z*_seq,*t*_ be the tokenized sequence at time *t* : 0 ≤ *t* ≤ *T*, which comprises *z*_*t*_ *(see Eq*. (62)). For a given position, *i* ∈ { 0, …, *L* − 1}, the infinitesimal generator matrix of the reverse CTMC is estimated by the model via probability ratios (see Section E.2). Sampling replaces `MASK` with content tokens while special tokens remain fixed. The probability ratios are estimated by:

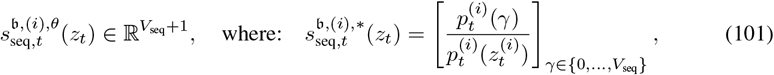

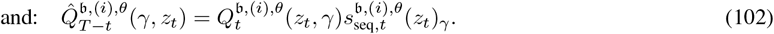

To form preference pairs, we use a batch comprising a wild type, *z*_seq,0_, *and B* mutant sequences with known scalar scores. We enumerate within-batch pairs, retain those with positive score gaps, sort by gap magnitude, and select the top-*k* (winner, loser) preference pairs, wherein the *j*-th pair is given by 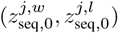 for *j* ∈ {0, …, *k* − 1}, and *k* ≤ *B(B* − *1)/2*. For our setting, as in [12], we compute the following D2-DPO loss over the (winner, loser) preference pairs, using the reference model, *θ*, and aligned model, *θ*^*′*^, where *P* denotes the distribution over winner-loser pairs:

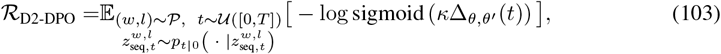

where we define the preference margin, 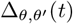, as:

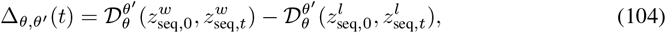

with the preference difference given by:

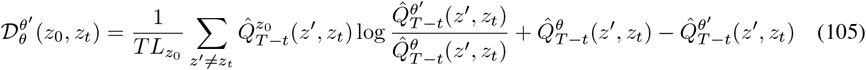

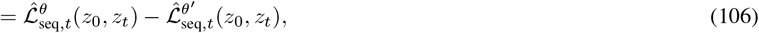

where 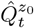 and 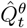 are defined as in Proof J.6. The hyperparameter *κ* regularizes the loss by implicitly penalizing the KL-divergence of alignment model to the reference model. Remarkably, we note that after some manipulation we recover that the D2-DPO preference margin can be written as the sums and differences of the estimators for the sequence and structure DWDSE losses.

#### I.2 Post-alignment Validation

Post-alignment via D2-DPO, we score the proteins generated by the aligned model (parameterized by *θ*^*′*^) by treating each generated sequence, 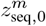, as a mutant, and the original unmasked sequence, *z*_seq,0_, as the wild type. It is also possible to use the same scoring to perform validation of the aligned model on any mutant protein 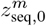 against a known wild type *z*_seq,0_. We evaluate our diffusion model by the following metric inspired by MLM infilling tasks and normalize by the number of masks.

Let 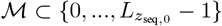, be the set of mutated positions (i.e., sequence positions wherein the residues differ between 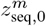 and 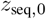). We compute a time *t*_infer_ to condition on via the equation:

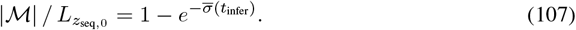

Let *z*^corr^ denote the mutant sequence, 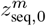, masked at ℳ, which also equals the wild type 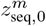 masked at ℳ. The estimated per-pair validation score is given by:

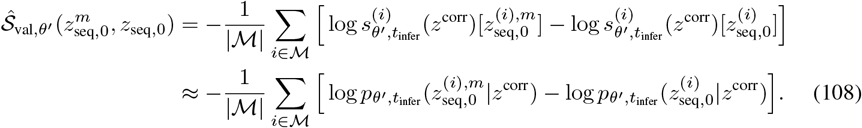

These scores measure the reference-calibrated preference for a mutant sequence over the wild type, aggregated across all mutated positions, which is standard for protein benchmarks [51].

### J Proofs

#### Remark E.1.

*For the geometric noise schedule of Eq. (63), the cumulative noise at time, t, is:*

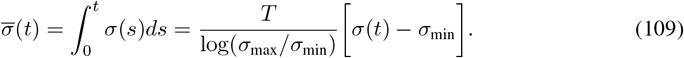

#### Derivation J.1.

To take the integral, we permit a continuous extension of the discrete-valued “stair-case” *σ*(*t*). We rearrange this function:

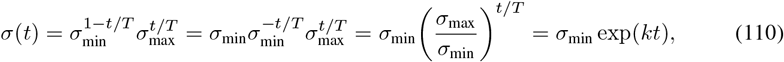

where *k* = log(*σ*_max_*/σ*_min_)*/T*. Thus:

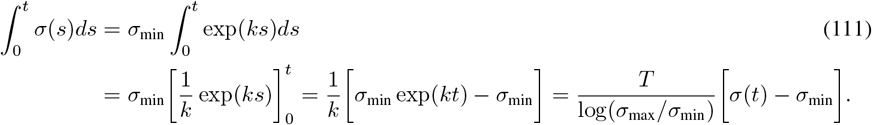

□

#### Lemma J.2

(Projective Matrix Exponent). *If P* ^2^ = *P, then* exp(*αP*) = *I* − *P + e*^*α*^*P*.

*Proof J.3.* Writing out the Taylor expansion:

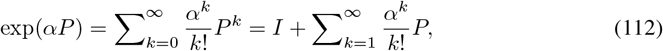

since *P* ^*k*^ *= P* for all *k* > 1. Accordingly, we have that:

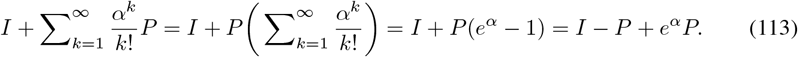

□

#### Remark E.2.

*The p*_*t*|0_^(*i*)^ *(which is a solution to Eq. (68)) can be expressed via the following, where* 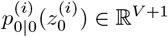 *is the one-hot vector corresponding to* 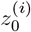:

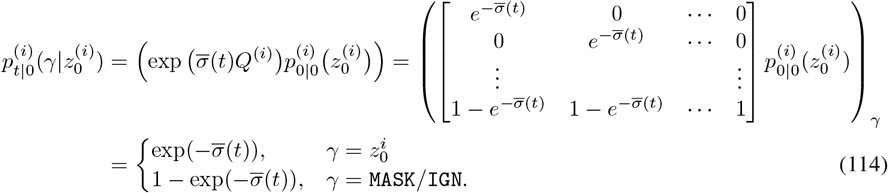

#### Derivation J.4.

Let *A* be a matrix with zeros in all but the last row, where:

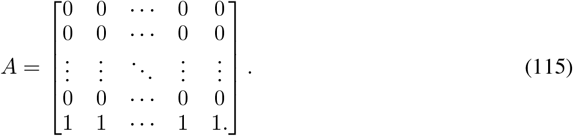

We express *Q*_absorb_ = *A* − *I. Noting that A*^2^ *= A, I*^2^ = *I* and applying Lemma J.2:

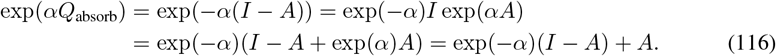

□

#### Theorem G.2

(Kabsch Algorithm). *For* 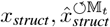 *with origin 3-centroids, and for a binary vector indicating non-*`MASK` *positions* 𝕄_*t*_ ∈ {0, 1} ^*L*^, *let R*_KABSCH_ *denote the output of Algorithm 1. Suppose SO*(3) *denotes the set of rotation matrices in* ℝ^3*×*3^. *Then:*

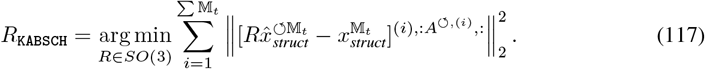

*Proof J.5*. The following proof is adapted from [34]. We define the index set 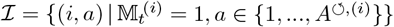, *with N* = |ℐ|. Ordering ℐ lexicographically by (*i, a*), we form:

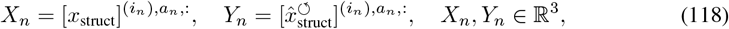

where (*i*_*n*_, *a*_*n*_) denotes the *n-th* element of ℐ. Stacking the rows, *(X*_*n*_)_*n*_, *(Y*_*n*_)_*n*_, into matrices, *X, Y* ∈ ℝ^*N ×*3^, we now rewrite the Kabsch objective in terms of *X* and *Y* as follows:

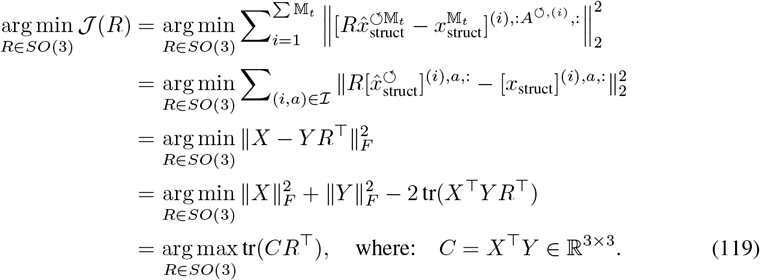

We perform a singular value decomposition (SVD) of *C* to obtain *C* = *U* Σ*V* ^⊤^, where *U, V* ∈ *O(3)*, Σ *= diag(σ*_1_, *σ*_2_, *σ*_3_), *σ*_1_ ≥ *σ*_2_ ≥ *σ*_3_ ≥ 0. Then, we have that:

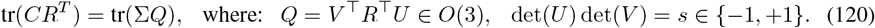

From von Neumann’s trace inequality, tr(Σ*Q*) ≤ *σ*_1_ + *σ*_2_ + *sσ*_3_, with equality at *Q* = diag(1, 1, *s*). Accordingly, it follows that:

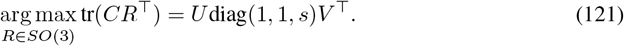

Algorithm 1 takes SVD of **reshape** 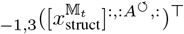 **reshape**

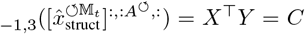, and returns *R*_KABSCH_ = *UKV* ^⊤^, where *K* = diag(1, 1, *s*). Thus, *R*_KABSCH_ = arg min_R∈SO(3)_ 𝒥 *(R)*.

#### Theorem G.4

(Score-Entropy and Perplexity Bound). *For* 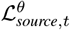 *given in Definition G.3*,

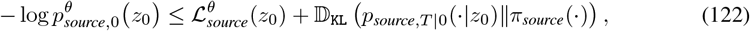

*Where* 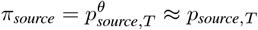 *is the initial distribution during generation*.

*Proof J.6*. The following is an adapted version of the corresponding proof in Theorem 3.6 of [44]. We reproduce it here for completeness.

Let 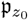 be the path measure of the CTMC generated by *Q*_*t*_ starting from 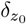, and note the generator for the reverse process 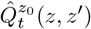 is given by:

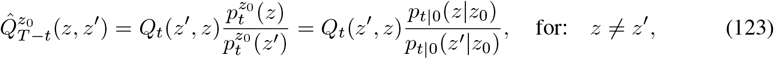

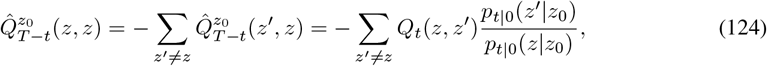

where 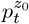 is the marginal distribution of 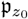 at time *t*. Next, let 𝔭^*θ*^ be the path measure of the learned reverse-time CTMC starting from *π* at time *T*, with generator 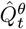. Recall that 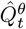 is defined by:

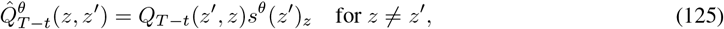

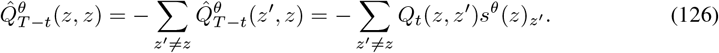

We can write down the following log-likelihood bound and manipulate using the data processing inequality:

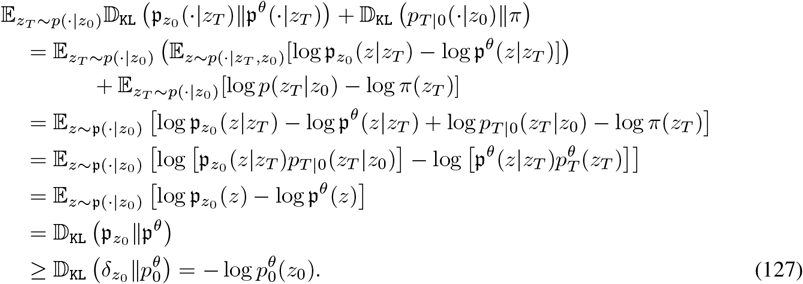

To obtain the theorem, it remains to show that ℒ_DWDSE_*(z*_0_) equals the quantity:

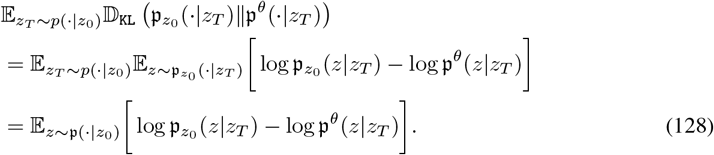

We fix a state *z*_*T*_ at time T and view it as the initial state for the reverse processes. We write the path log-likelihoods in terms of jumps and intensities, following Equation 2 of [59]:

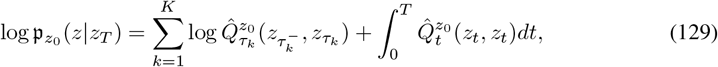

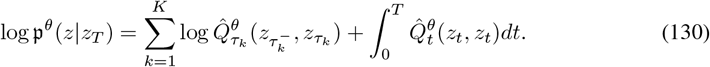

Then, Eq. (128) is equivalent to:

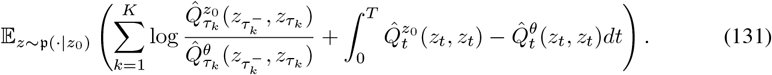

Applying Dynkin’s formula [23] to the first term, we obtain:

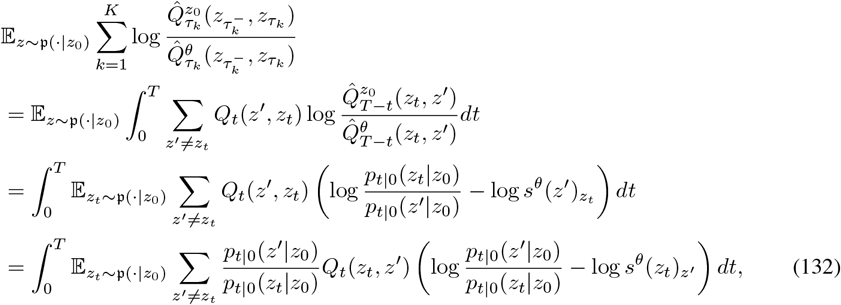

where in the third equality we leveraged the key identity:

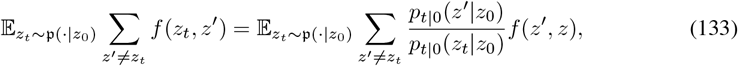

for any function *f* (*z, z*^*′*^). Regarding the second term of Eq. (131), we have that:

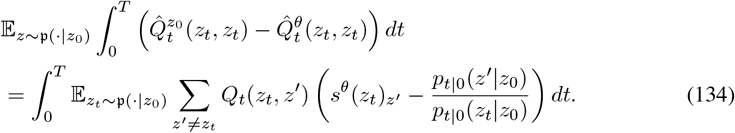

Combining the results of Eq. (132) and Eq. (134), we obtain:

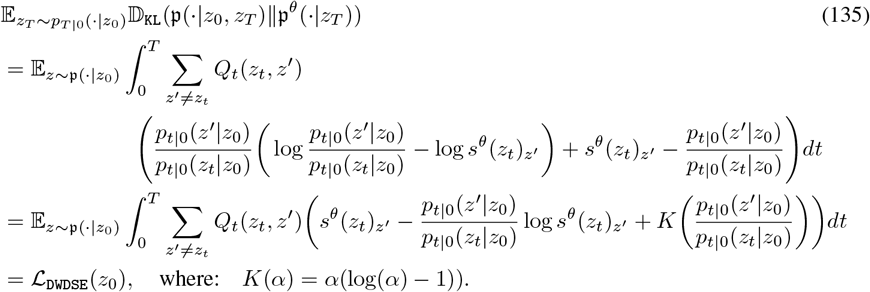

□

#### Remark G.6.

*If t* ∼ 𝒰 *([0, T]), and z*_*t*_ *drawn following Remark E.2, then the estimator*, 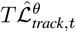 *of Definition G.5, is an estimator of the* 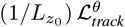.

*Proof J.7*. The proof below adapts the theory of [44]. We first rewrite the *t*-integral as an expectation:

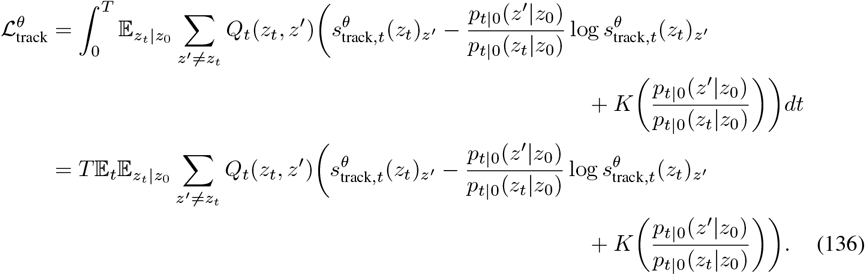

*For sequences z*_*t*_ ≠ *z*^*′*^, the generator *Q*_*t*_*(z*_*t*_, *z*^*′*^) is nonzero only if *z*_*t*_ and *z*^*′*^ differ by exactly one token at an index *i* and *z*_*t*_ is masked at *i;* that is,

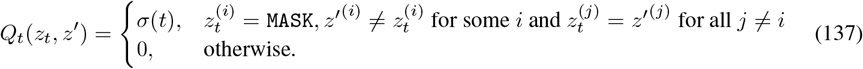

Consequently, we have that:

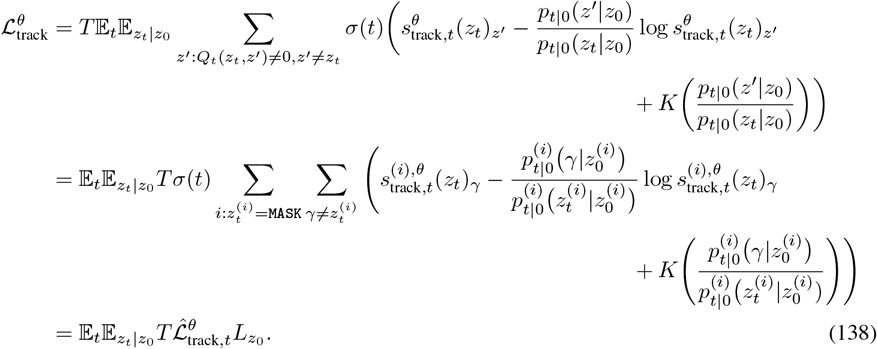

Therefore, 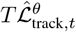 is an unbiased estimator of 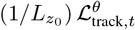 under the stated sampling scheme.□

#### Remark H

(Reverse Timestep using Tweedie’s Method and Tau Leaping). *The following update rule numerically approximates the reverse process:*

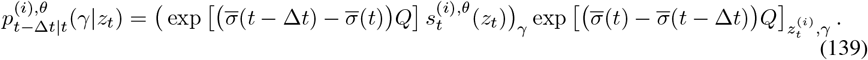

#### Derivation J.8.

The following is an adapted version of the corresponding proof in Theorem 4.2 of [44]. We reproduce it here for completeness.

For non-negative vectors *p*_0_, *p*_*t*_ ∈ ℝ^*V +1*^ with sum one, and following the ODE:

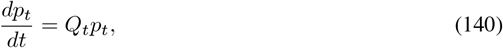

we have that:

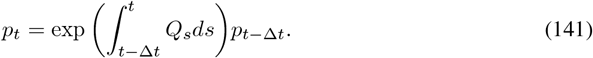

Moreover, with Q_t_ = σ(t)Q, we have that:

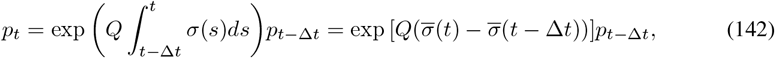

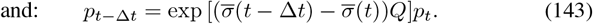

Next, we consider:

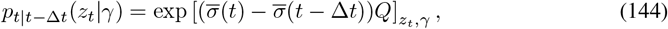

since exp 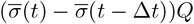 is a transition matrix. By Bayes’ rule:

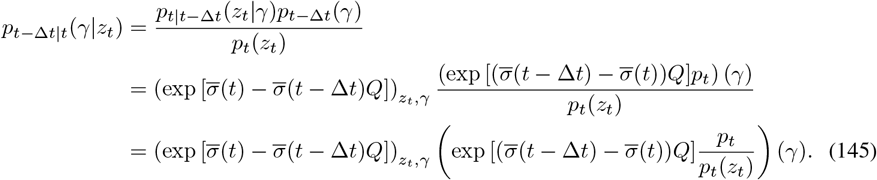

In the exposition of Eq. (145), we note that each *z*_*t*_ is a full sequence. For large *L*, it is intractable to estimate *p*_*t*_(*γ*)*/p*_*t*_(*z*_*t*_)) for all (*V* + 1)^*L*^ outcomes of *γ*. Our transformer output 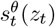 only models the probability ratios between sequences that differ in one position. For positive Δ*t*, however, it is possible that the true reverse process will produce a *z*_*t−*Δ*t*_ that has a Hamming distance greater than one to *z*_*t*_. For small Δ*t*, 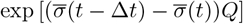 does not generate these transitions with high probability, and hence we may make the approximation to ignore entries of 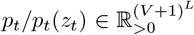 other than those predicted by 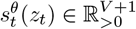.

The transition of Eq. (139) is an approximation to the true reverse process. In our implementation, we choose a small value of Δ*t* to make this approximation precise. For a more rigorous treatment of tau-leaping, we refer the reader to [44]. □

### K Additional empirical results

#### K.1 Benchmarking consensus versus attention

As an addendum to Section 3.2, we provide detailed sequence and structure breakdowns for scaled masked language models trained via consensus and attention in Figures 23 and 24. These figures broadly follow the same pattern discussed in the main text, wherein models trained with consensus are significantly more stable at higher learning rates. Moreover, consensus exhibits a broader interval, *W*, than self-attention, a pattern that is is much more pronounced with larger model sizes.

**Figure 23:**
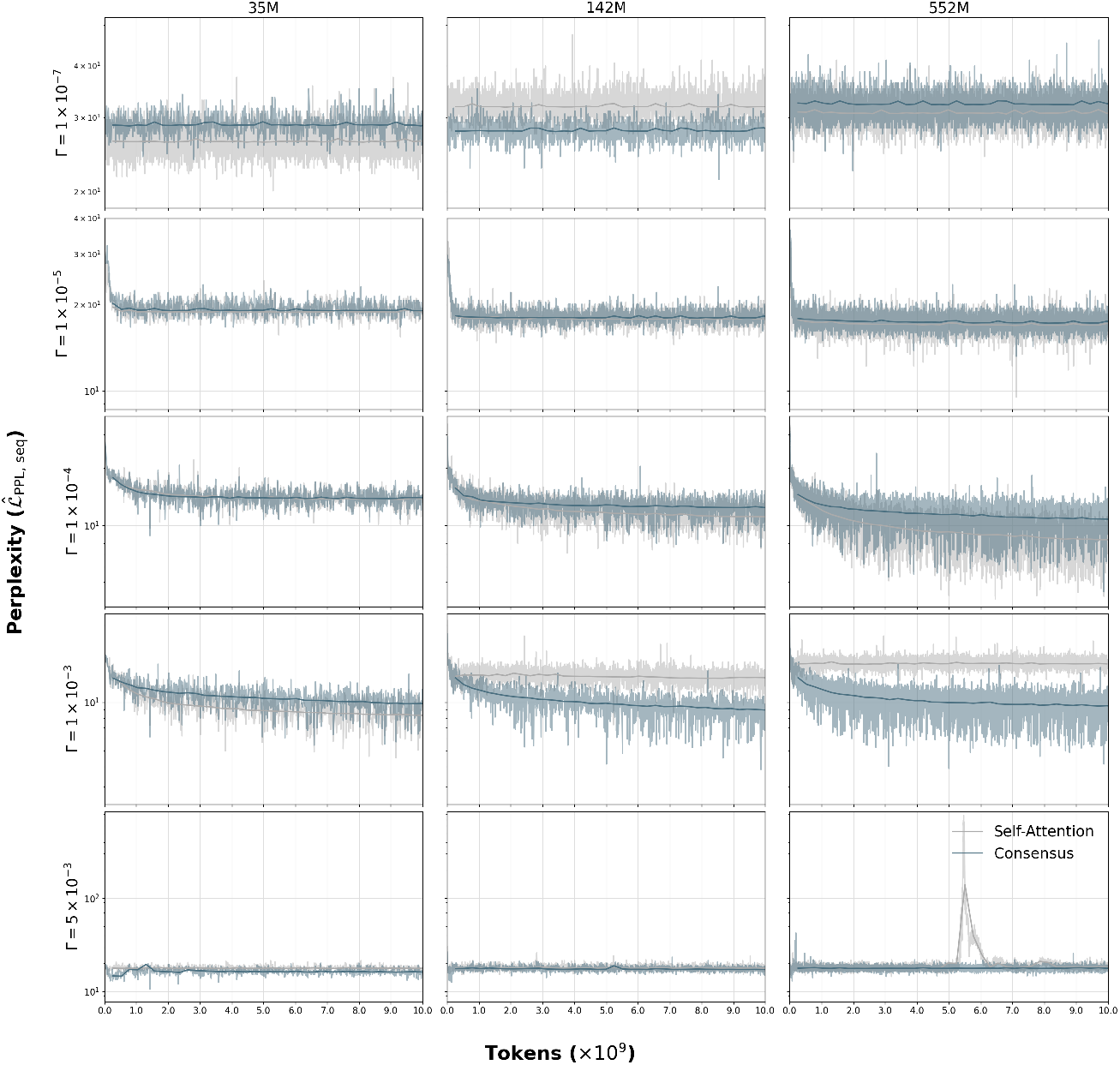
We train several masked language models (35M, 142M, 552M parameters) with consensus (blue) or attention (gray) via simple masking. The sequence training and validation perplexities form the background and solid lines in the foreground, respectively. For variable learning rate, Γ, we see improved robustness using consensus vs attention, with heightened effects at larger model sizes.

**Figure 24:**
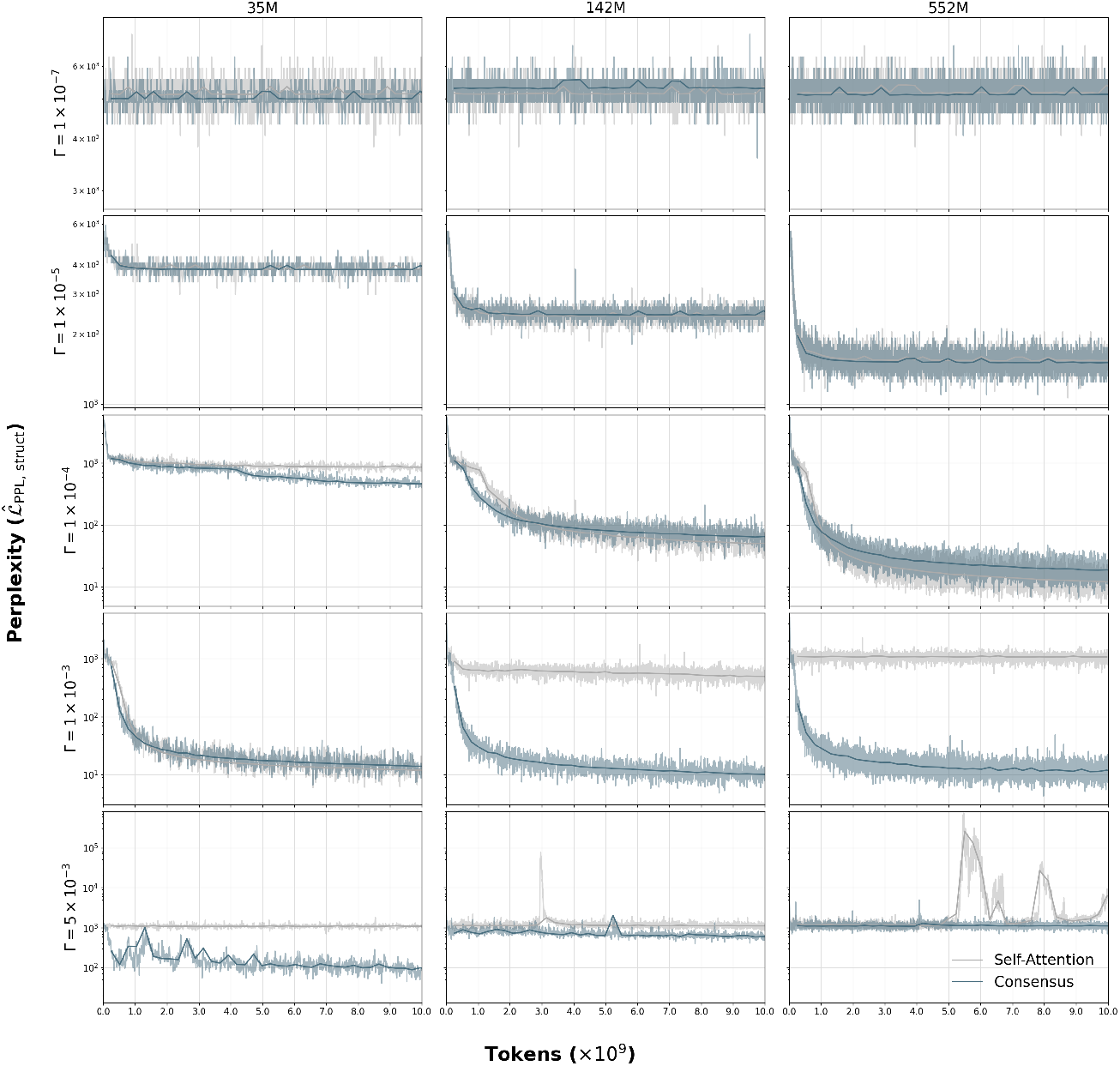
We train several masked language models (35M, 142M, 552M parameters) with consensus (blue) or attention (gray) via simple masking. The structure training and validation perplexities form the background and solid lines in the foreground, respectively. For variable learning rate, Γ, we see improved robustness using consensus vs attention, with heightened effects at larger model sizes.

#### K.2 Tracking alignment

We track and visualize the training progress of D2-DPO alignment (from Section 4) for enzymes with PDB IDs 1BS9, 1CWY, and 8PCH (see Figures 25, 26, 27). We see consistent increases in Spearman correlation, *ρ*, between true scores (composite score of two measures of protein fitness) and predicted multi-objective scores (as defined by our alignment validation metric in Eq. (108)) with more training epochs. This demonstrates the effectiveness and reliability of D2-DPO alignment for taking unaligned reference models, whose predicted scores have low correlation to protein fitness, and steering toward alignment for predicting protein fitness, across proteins not seen during pre-training.

**Figure 25:**
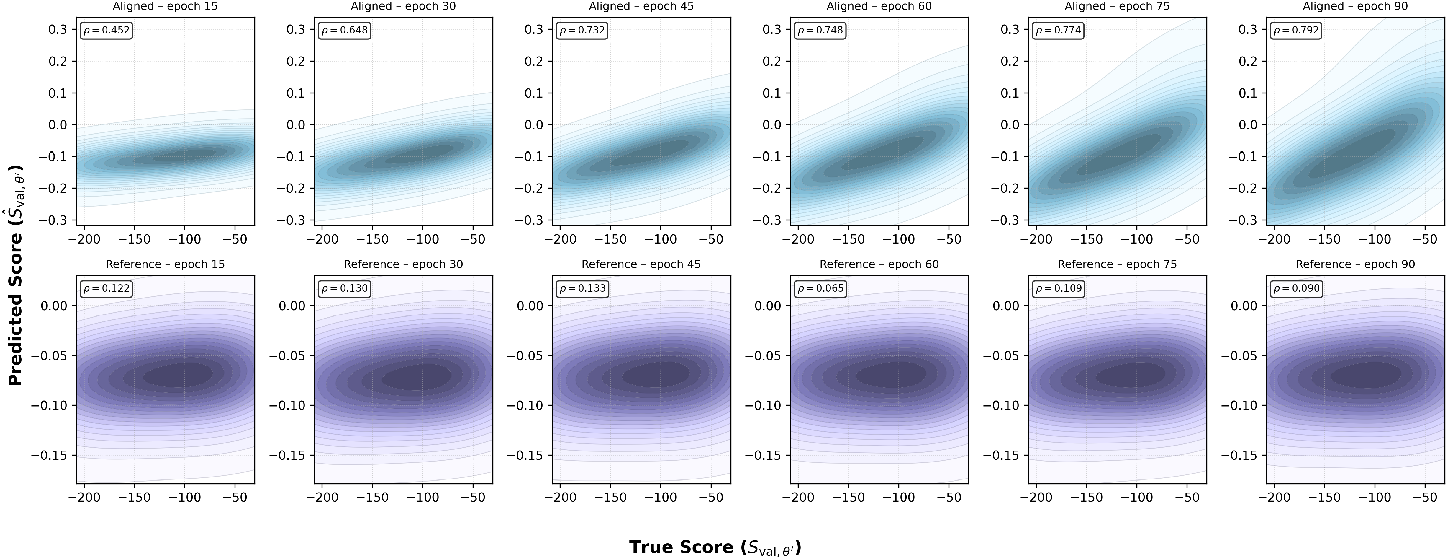
Alignment across training epochs for 1BS9.

**Figure 26:**
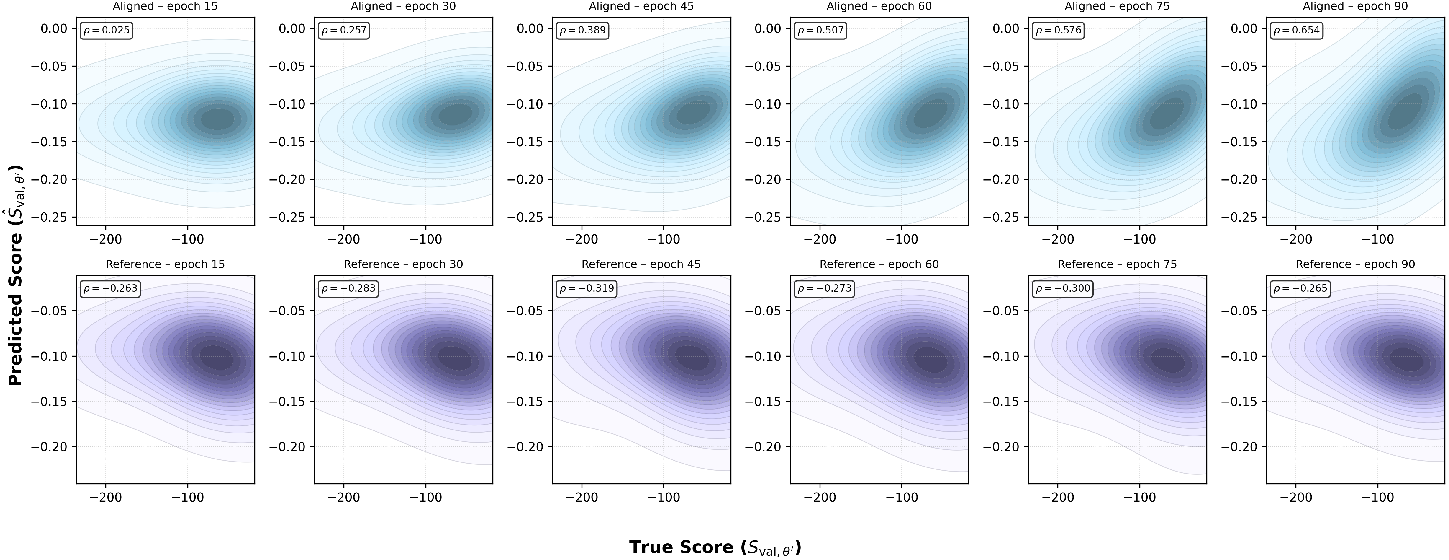
Alignment across training epochs for 1CWY.

**Figure 27:**
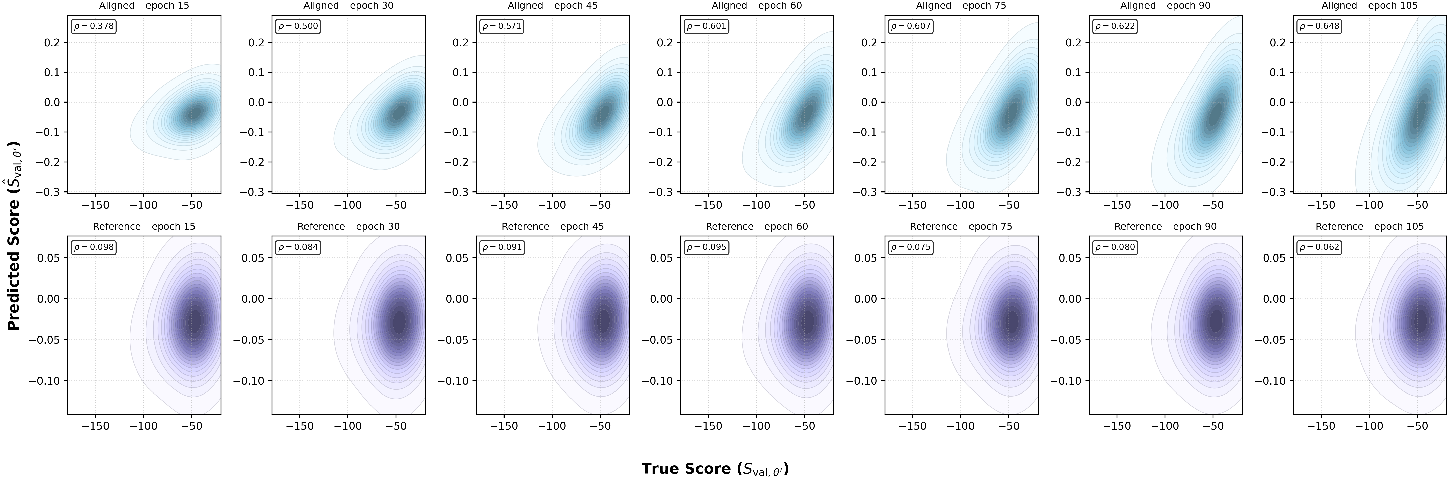
Alignment across training epochs for 8PCH.

#### K.3 Model configurations and training hyperparameters

To train the FSQ and all transformer stack variants, we utilize a fixed learning-rate schedule, which comprises a warm-up period over the first 10% of total training steps to the peak learning rate [36], a linear decay to the minimum learning rate over the next 40% of training steps [8], and a constant leveling at the minimum learning rate for the remaining 50% of training steps. Table 10 summarizes the model configurations and training hyperparameters used for our production-ready models. We use the AdamW optimizer to train all models [43]. The full tabulation of considered model configurations and training hyperparameters is provided in Table 10.

**Table 5:** Sequence tokenization and mapping scheme.

| ID | Symbol | Name | ID | Symbol | Name |
| --- | --- | --- | --- | --- | --- |
| 0 | A | Alanine | 13 | F | Phenylalanine |
| 1 | R | Arginine | 14 | P | Proline |
| 2 | N | Asparagine | 15 | S | Serine |
| 3 | D | Aspartic acid | 16 | T | Threonine |
| 4 | C | Cysteine | 17 | W | Tryptophan |
| 5 | Q | Glutamine | 18 | Y | Tyrosine |
| 6 | E | Glutamic acid | 19 | V | Valine |
| 7 | G | Glycine | 20 | B | Asparagine or Aspartic acid |
| 8 | H | Histidine | 21 | U | Selenocysteine |
| 9 | I | Isoleucine | 22 | Z | Glutamic acid or Glutamine |
| 10 | L | Leucine | 23 | O | Ornithine |
| 11 | K | Lysine | 24 | J | Leucine or Isoleucine |
| 12 | M | Methionine | — | — | — |

**Table 6:**
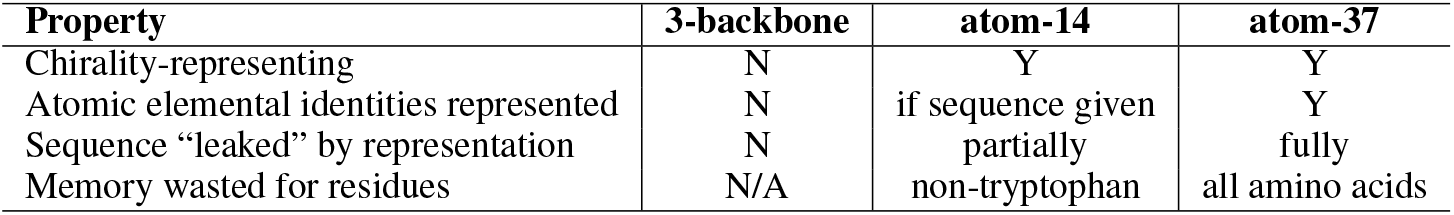
Comparison of atom coordinate representation choices.

**Table 7:** Secondary structure tokenization and mapping scheme.

| ID | Symbol | Name | Description |
| --- | --- | --- | --- |
| 0 | C | Coil / Loop | Non-regular backbone geometry |
| 1 | E | Extended strand | β-sheet ladder participation |
| 2 | B | Isolated β-bridge | Single-residue $\beta$ bridge |
| 3 | T | Turn | Hydrogen-bonded reversal |
| 4 | S | Bend | Geometric bend (non H-bonded) |
| 5 | H | α-helix | Canonical helix (i→i+4) |
| 6 | G | $3_{10}$ helix | Helix (i→i+3) |
| 7 | I | π-helix | Helix (i→i+5; rare) |
| 8 | P | Polyproline II | Left-handed PPII (non-DSSP extension) |

**Table 8:** Solvent-accessible surface area tokenization and mapping scheme.

| ID | Bin | Range ($\text{\AA}^2$) | ID | Bin | Range ($\text{\AA}^2$) |
| --- | --- | --- | --- | --- | --- |
| 0 | BIN_0 | 0.00–0.14 | 8 | BIN_8 | 47.43–56.62 |
| 1 | BIN_1 | 0.14–2.09 | 9 | BIN_9 | 56.62–65.50 |
| 2 | BIN_2 | 2.09–6.49 | 10 | BIN_10 | 65.50–76.08 |
| 3 | BIN_3 | 6.49–12.69 | 11 | BIN_11 | 76.08–86.71 |
| 4 | BIN_4 | 12.69–20.23 | 12 | BIN_12 | 86.71–99.48 |
| 5 | BIN_5 | 20.23–29.03 | 13 | BIN_13 | 99.48–115.35 |
| 6 | BIN_6 | 29.03–38.17 | 14 | BIN_14 | 115.35–137.27 |
| 7 | BIN_7 | 38.17–47.43 | 15 | BIN_15 | $\geq 137.27$ |

**Table 9:**
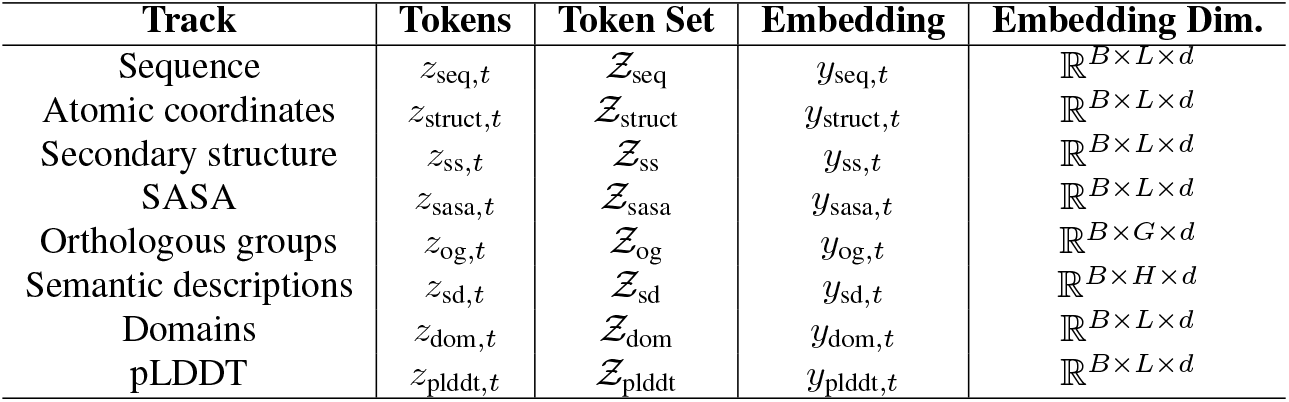
Tokenization and embedding dimensions of Odyssey tracks.

| Track | Tokens | Token Set | Embedding | Embedding Dim. |
| --- | --- | --- | --- | --- |
| Sequence | $z_{\text{seq},t}$ | $\mathcal{Z}_{\text{seq}}$ | $y_{\text{seq},t}$ | $\mathbb{R}^{B \times L \times d}$ |
| Atomic coordinates | $z_{\text{struct},t}$ | $\mathcal{Z}_{\text{struct}}$ | $y_{\text{struct},t}$ | $\mathbb{R}^{B \times L \times d}$ |
| Secondary structure | $z_{\text{ss},t}$ | $\mathcal{Z}_{\text{ss}}$ | $y_{\text{ss},t}$ | $\mathbb{R}^{B \times L \times d}$ |
| SASA | $z_{\text{sasa},t}$ | $\mathcal{Z}_{\text{sasa}}$ | $y_{\text{sasa},t}$ | $\mathbb{R}^{B \times L \times d}$ |
| Orthologous groups | $z_{\text{og},t}$ | $\mathcal{Z}_{\text{og}}$ | $y_{\text{og},t}$ | $\mathbb{R}^{B \times G \times d}$ |
| Semantic descriptions | $z_{\text{sd},t}$ | $\mathcal{Z}_{\text{sd}}$ | $y_{\text{sd},t}$ | $\mathbb{R}^{B \times H \times d}$ |
| Domains | $z_{\text{dom},t}$ | $\mathcal{Z}_{\text{dom}}$ | $y_{\text{dom},t}$ | $\mathbb{R}^{B \times L \times d}$ |
| pLDDT | $z_{\text{plddt},t}$ | $\mathcal{Z}_{\text{plddt}}$ | $y_{\text{plddt},t}$ | $\mathbb{R}^{B \times L \times d}$ |

**Table 10:** Production-ready model configurations and training hyperparameters.

| Model | # Params<br>($\mathcal{K}$) | # Layers<br>($N$) | # Heads<br>($H$) | Model Dim.<br>($d$) | Peak LR<br>($\Gamma_{\max}$) | Min LR<br>($\Gamma_{\min}$) |
| --- | --- | --- | --- | --- | --- | --- |
| FSQ Stage-1 | 50M | 16 | 6 | 384 | 3e-5 | 1e-6 |
| FSQ Stage-2 | 500M | 48 | 16 | 1024 | 3e-5 | 1e-6 |
| Odyssey-35M | 35M | 12 | 4 | 256 | 1e-4 | 1e-5 |
| Odyssey-142M | 142M | 24 | 8 | 512 | 1e-4 | 1e-5 |
| Odyssey-1.2B | 1.2B | 32 | 24 | 1536 | 4e-4 | 4e-5 |
| Odyssey-12B | 12B | 48 | 64 | 4096 | 2.5e-4 | 2.5e-5 |
| Odyssey-102B | 102B | 106 | 128 | 8192 | 6e-4 | 6e-5 |

In the scaling experiments, we train each transformer stack using a constant learning rate and discrete diffusion masking schedule. We add warmup steps for larger models to ensure stable training. These parameters are detailed in Table 11. In our ablation study benchmarking consensus versus attention across learning rate, we use configurations detailed in Table 12. For all model sizes, we use a constant learning rate and 300 warmup steps. In addition, we only compare a simple masking schedule when testing Attention and Consensus mechanisms. In our ablation study benchmarking discrete diffusion versus masked language modeling, we train with a flat learning rate scheduler and 300 warmup steps, using the hyperparameters depicted in Table 13.

**Table 11:** Model and training parameters for scaling experiments.

| # Params | # Layers<br>($N$) | # Heads<br>($H$) | Model Dim.<br>($d$) | LR<br>($\Gamma$) | # Warmup<br>Steps |
| --- | --- | --- | --- | --- | --- |
| 142M | 24 | 8 | 512 | 7.5e-5 | 0 |
| 284M | 24 | 12 | 768 | 1e-4 | 0 |
| 592M | 32 | 16 | 1024 | 2e-4 | 0 |
| 1.2B | 32 | 24 | 1536 | 4e-4 | 300 |

**Table 12:** Model and training parameters for consensus vs attention ablation study.

| # Params | # Layers<br>($N$) | # Heads<br>($H$) | Model Dim.<br>($d$) | Learning Rates ($\Gamma$) | Model<br>Mechanism |
| --- | --- | --- | --- | --- | --- |
| 35M | 12 | 4 | 256 | 1e-7, 1e-5, 1e-4, 1e-3, 5e-3 | Attention |
| 35M | 12 | 4 | 256 | 1e-7, 1e-5, 1e-4, 1e-3, 5e-3 | Consensus |
| 142M | 24 | 8 | 512 | 1e-7, 1e-5, 1e-4, 1e-3, 5e-3 | Attention |
| 142M | 24 | 8 | 512 | 1e-7, 1e-5, 1e-4, 1e-3, 5e-3 | Consensus |
| 552M | 32 | 16 | 1024 | 1e-7, 1e-5, 1e-4, 1e-3, 5e-3 | Attention |
| 552M | 32 | 16 | 1024 | 1e-7, 1e-5, 1e-4, 1e-3, 5e-3 | Consensus |

**Table 13:** Model and training parameters for discrete diffusion vs masked language modeling study.

| # Params | # Layers<br>($N$) | # Heads<br>($H$) | Model Dim.<br>($d$) | LR<br>($\Gamma$) | Masking Scheme |
| --- | --- | --- | --- | --- | --- |
| 35M | 12 | 4 | 256 | 1e-4 | Simple/Complex Masking<br>Discrete Diffusion |
| 142M | 24 | 8 | 512 | 1e-4 | Simple/Complex Masking<br>Discrete Diffusion |
| 1.2B | 32 | 24 | 1536 | 4e-4 | Simple/Complex Masking<br>Discrete Diffusion |

